# Effects of technical noise on bulk RNA-seq differential gene expression inference

**DOI:** 10.1101/843789

**Authors:** Dylan Sheerin, Daniel O’Connor, Andrew J Pollard, Irina Mohorianu

## Abstract

**Motivation:** Inconsistent, analytical noise introduced either by the sequencing technology or by the choice of read-processing tools can bias bulk RNA-seq analyses by shifting the focus to the variation in expression of low-abundance transcripts; as a consequence these highly-variable genes are often included the differential expression (DE) call and impact the interpretation of results.

**Results:** To illustrate the effects of “noise”, we present simulated datasets following closely the characteristics of a *H.sapiens* and a *M.musculus* dataset, respectively, highlighting the extent of technical-noise in both a high inter-individual variability (*H. sapiens*) and reduced variability (*M. Musculus*) setup. The sequencing-induced noise is assessed using correlations of distributions of expression across transcripts; analytical noise is evaluated through side-by-side comparisons of several standard choices. The proportion of genes in the noise-range differs for each tool combi-nation. Data-driven, sample-specific noise-thresholds were applied to reduce the impact of low-level variation. Noise-adjustment reduced the number of significantly DE genes and gave rise to convergent calls across tool combinations.

**Availability:** The code for determining the sequence-derived noise is available for download from: https://github.com/yry/noiseAnalysis/tree/master/noiseDetection_mRNA; the code for running the analysis is available for download from: https://github.com/sheerind/noise_detection.

## 1 Introduction

The increase in sequencing depth for RNA sequencing (RNA-seq) experiments *(1)* allows a higher sensitivity for the detection of perturbations in gene expression levels between samples *(2)*. This increased accuracy facilitated the characterisation of differential expression (DE) at tissue and cellular levels *(3)*. However, non-biological/technical variation of signal can be introduced at different stages of the RNA-seq library preparation and sequencing and varies from amplification/sequencing bias *(4)* to random hexamer priming during the sequencing reaction *(5)*; these technical alterations of signal can affect the precision/accuracy of the downstream DE call. Statistical methods focused mainly on batch/background correction, normalisation, and evaluation of DE have been developed to mitigate the impact of these biases on DE analyses *(6)*; however these approaches are not designed to identify and assess the impact of genes showing random, low-level variation (noise), some of which are included in the DE call and thus impact the interpretation of the DE analysis. Additionally, the choices of pre-processing tools also influence the output (and relative quantification accuracy) of gene expression *(7)*. These analytical biases mainly arise from differences in the detection and handling of isoforms or processing of unmapped and multi-mapping reads *(3)*. In the present study, we compare a selection of commonly used alignment and transcript-quantification tools to assess the impact of computational choices, corroborated with the evaluation of technical noise, to the downstream DE call.

A recent benchmarking study of widely used RNA-seq aligners *(8)* compared their relative accuracy when applied to simulated data of varying sequence complexity. The results suggest a poor correlation between the performance of these tools and their relative frequency of use i.e. widely-cited tools appear to underperform based on the metrics used in the study, particularly when default parameters are applied. Several other benchmarking studies have also evaluated the landscape of RNA-seq aligners under different dataset-specific criteria *(9–12)* allowing the users to make informed decisions as to which tool to use based on the particularities of their datasets. In practice, transcript- and gene-level estimation tools are often used in combination with HISAT2 *(13)* and STAR *(14)*. The HISAT and STAR alignment tools differ mainly in their use of compression; HISAT utilises an extension of the Burrows-Wheeler transform compression algorithm *(15)*, reducing its memory requirement to <8GB, while STAR is more RAM-intensive due to its use of uncompressed suffix arrays for ultrafast alignment. However, speed and memory usage are not the sole considerations when choosing an alignment tool; an important criterion when deciding which of the ever-growing number of read alignment tools to use is the ability to integrate it into pipelines for downstream estimation of expression and subsequent call of DE.

StringTie *(16)* performs a transcript-level quantification using a network flow algorithm, in combination with optional *de novo* assembly methods; it is a recommended tool for quantifying HISAT2-aligned reads *(17)*. Genomic overlap between isoforms hinders the StringTie quantification on this particular set of transcripts; in comparison the gene-level estimation (e.g. union exon methods) are based on the simpler merging overlapping exons and quantification based on algebraic sum of incident reads. However, union-exon methods result in significant underestimation of gene expression levels when compared with transcript-based approaches *(7)*. HTseq (htseq-count) *(18)* and featureCounts *(19)* are two similar union exon-based transcript quantification tools; featureCounts has been shown to out-perform the htseq-count in speed and accuracy *(19)*. The use of the *–quantMode GeneCounts* option in STAR produces and identical output as the one from htseq-count. As the accuracy of quantification relies heavily on the identification of the transcripts from which reads originate, more than the precise characteristics of the alignment of these reads to the transcript, recently developed pseudoalignment tools, such as kallisto *(20)*, utilize hash tables to build the index, bypass the alignment step, and base transcript identification on the length of k-mers and the mean of the distribution of fragments, increasing computational speed while maintaining accuracy.

In the present study, we incorporate a statistical method to assess the variability of signal across transcripts *(21)* as part of standard DE pipelines and thus evaluate the combined effect of sequencing noise and analytical noise on the DE call. This method offers a data-driven alternative to the *ad hoc* (and sample-independent) filtration of low abundance genes, based on the counts per million (CPM) *(22, 23)*. We use the inferred thresholds to compare the proportion of “noisy” genes for combinations of the read alignment and quantification tools and assess the impact on the downstream DE call. Finally, we show that the variation derived from protocol-specific features can be reduced by the exclusion of genes with abundance below user-defined noise thresholds (specific for individual samples), and lead to the convergence in the identity of differentially expressed genes (DEGs) between method combinations.

## 2 Materials and Methods

The *H sapiens* comprises of 30 paired (control vs treatment) samples. The *M musculus* comprises 18 unpaired samples. The simulated datasets followed the sequencing depth, nucleotide composition and sequencing bias of real datasets.

The files corresponding to the *H sapiens* GRCh38.97 and *M musculus* GRCm38.97 genomes were downloaded from www.ensembl.org; the genome sequence, corresponding annotations (GRCh38.97.gtf and GRCm38.97.gtf) and the nucleotide sequences for the protein coding genes were used for alignments and gene quantifications.

### 2.1 Preprocessing of reads

HISAT2 v2.1.0, *(13)* was for the probabilistic alignment of reads to the reference genome index with default parameters. Samtools v0.1.18, *(24)* was used to convert the SAM output to a coordinate-sorted BAM file for quantification. Alignments generated by STAR (v2.4.0.1, *(14)*) are fully deterministic but the order of alignments in the output are only deterministic when run with one single thread and default parameters.

Transcript-based quantification was performed with StringTie v1.3.5, *(16)* estimating transcripts from the reference annotation file. For union exon-based quantification, htseq-count v0.10.0, *(18)* was used to count reads at the exon level. The STAR --quantMode GeneCount option was used to output a ReadsPerGene count file, similar to that of htseq-count, for each sample. Finally, featureCounts (v1.6.2, *(19)*) was used to provide an additional union-exon quantification.

### 2.2 Noise detection

Using the transcript coordinates of the aligned reads as input, the expression profile for each individual transcript is built as an algebraic point sum of the abundances of reads incident to any given position; if the alignment is performed per read, the corresponding abundance for every entry is set to +1. For each sample j, and for each transcript T, the point-to-point Pearson Correlation between the expression profile in j and the one in all other samples is calculated. The noise detection is based on the relative location of the distribution of the point-to-point Pearson Correlation Coefficient (p2pPCC, *(21)*) versus the abundances of genes and is specific for each individual sample. For low abundance transcripts the stochastic distribution of reads across the transcript leads to a low p2pPCC; the aim of the approach is to determine the range where the distribution of correlation coefficients (used as proxy for the distribution of reads across a transcript) are above a user-defined threshold; to approximate the signal-to-noise threshold a binning on the abundances is performed. For all examples presented in this study, the binning is done on log_2_ ranges; the signal-to-noise thresholds were defined as the abundance at which the median of the p2pPCC distribution within a bin exceeds 0.7. A range for the signal-to-noise can be defined on: [lower bound] the first occurrence of a distribution for which the median exceeds 0.7 and [upper bound] the last distribution for which the median is lower than 0.7. The correlation threshold, the localization of the distribution and the binning range are user-defined parameters. The localization parameter can be chosen as the lower/upper quartile (25/75%) or top 5%, 95%. We do not recommend the usage of local extremes (min/max) due to the high inter-sample variability. We also recommend a log-base binning approach that will assign at least 25 transcripts per bin for the bins around the mod of the abundance distribution.

### 2.3 Differential gene expression call

Read count matrices corresponding to all combinations of alignment and gene quantification tools were used for the downstream analysis and duplicated to facilitate the side-by-side comparison. Counts per million (CPM) filtering was applied to one matrix, applying a threshold proportional to the sequencing depth (a gene count of 10/minimum library size in millions); both count matrices were normalised using the trimmed mean of m-values (TMM) method *(22)*. The matrices were again duplicated, and the inferred noise thresholds were applied to one set of matrices from each pair; briefly, within a sample j, all entries below the signal-to-noise threshold were set to the threshold level. Entries with no value above the signal-to-noise thresholds, in any sample, were excluded from further analyses. The subsequent DE analysis was conducted using the Bioconductor package limma *(25)*. Weights were calculated using limma *voom (26)*. The signal-to-noise thresholds obtained on the raw data were factor-transformed to correspond to the TMM normalised distribution of values; all entries for which the normalised gene expression was below the threshold were set to the threshold value and values equivalent to the sum of the factor-transformed noise thresholds were removed from the analysis. Linear models were fitted to the data using the *lmfit* function, and a trended empirical Bayes, *eBayes*, method was applied to account for the mean-variance relationship of the data and improve accuracy of the log_2_ fold change (LFC) estimations *(25)*. Significantly DEGs were defined as those with a false-discovery rate (FDR) adjusted p-value, calculated using the Benjamini-Hochberg method *(27)*, of < 0.01, and |LFC| > 1.

## 3 Results

### 3.1 Variation in the quantification of gene expression for combinations of pre-processing tools is attributable to the choice of transcript quantification tool

Run summaries were generated for the input data and on the read alignment output comprising of total and unique number of reads, for the former, and of percentages of mapped reads for each method, for the latter (Supplementary Table 1). The proportions of uniquely mapping reads varied in the 87.69–97.78% (median 91.63%) range for STAR and 83.24–90.23% (median 87.5%) range for HISAT2. STAR captured a greater number of reads mapping to the reference genome, with as few as 2.28–6.62% (median 3.14%) not aligned compared with 7.09-12.94% for HISAT2 (median 8.07%), but also retrieved a higher percentage of multi-mapping reads (3.55–9.23%, median 5.03%) than HISAT2 (2.59–6.77%, median 3.72%). The distributions of the mapping proportions for H sapiens and M Musculus datasets are presented in Fig. 1A–C.

**Fig. 1.**
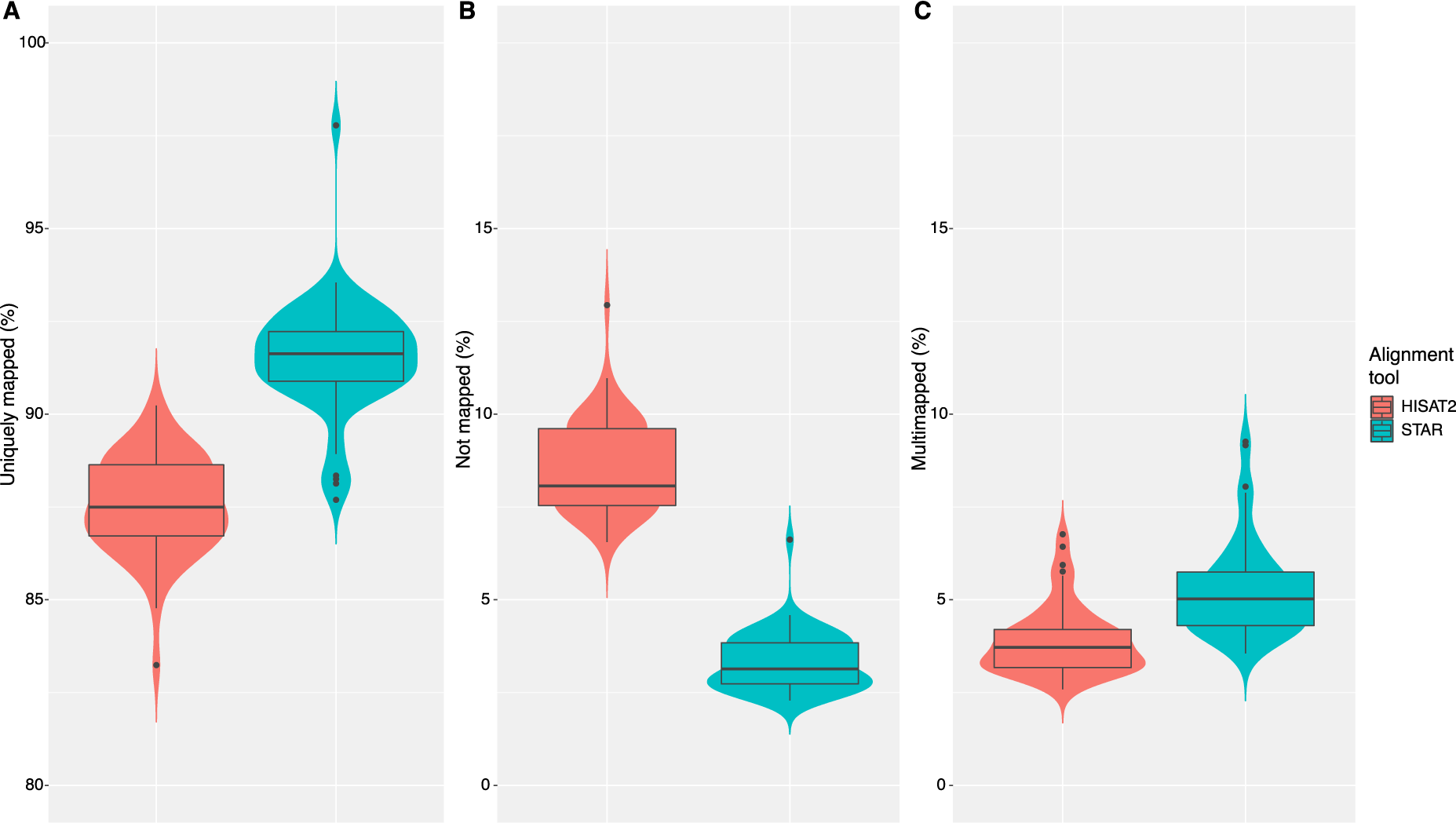
Run summaries for HISAT2 and STAR alignment tools. (A) results for H. sapiens, (B) results for M. musculus. The violin plots show the distribution of percentages of total reads mapping to the reference genome **A.** uniquely mapped, **B.** not mapped, and **C.** multi-mapped. The overlaying boxplots show the median and interquartile ranges of the distributions. STAR produces higher proportions of uniquely mapping reads, after successfully shifting these from the un-mapped category.

The total number of genes called (with non-zero expression level) by each alignment/quantification combination was consistent between HISAT2- and STAR-aligned outputs (37814–39370, median of absolute differences: 222 genes, Supplementary Table 2). The variation in total numbers of expressed genes appears to be attributable mainly to the choice of quantification tool since it remains relatively consistent between combinations with the same quantification tool even when different alignment tools are used (e.g. 39370 and 39592 for HISAT2-HTSeq and STAR-HTSeq options, respectively). A density plot of the abundances of genes uniquely called as expressed (i.e. called as expressed by one method only) by each tool combination shows that the genes in the specific differences are predominantly low abundance genes (Fig. 2A, 83.44% genes with abundance less than the median [blue line], 16.56% genes with abundance greater than the median).

**Fig. 2.**
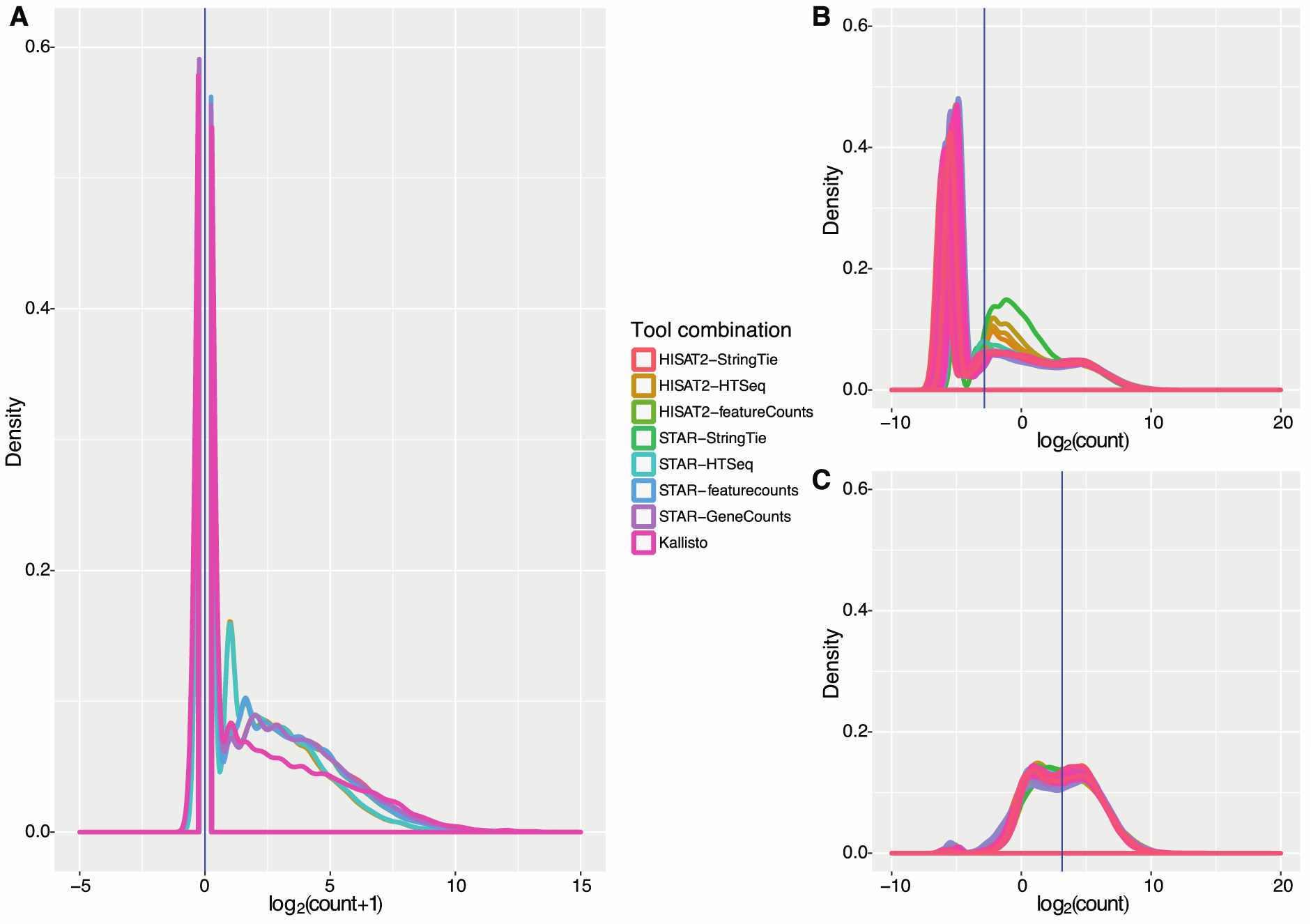
Gene count density plots. **A.** Distribution of expression levels, based on initial pre-normalised pseudocounts from non-zero matrices of expressed, unique genes as called by each read alignment/quantification tool combination for a randomly selected sample. **B.** Distribution of expression levels of unfiltered, TMM-normalised, *voom*-transformed counts for each sample processed by a single tool combination (HISAT2-StringTie); we observe that this combination of tools generates some anomalies (samples coloured in orange, ochre, mustard, and green), all of which correspond to D1 (post-immunisation) samples, that present a specific subset of genes expressed at medium abundance. **C.** Expression levels of CPM-filtered, TMM-normalised, *voom*-transformed counts for each sample processed by the same tool combination (HISAT2-StringTie); here we observe the effect of the CPM filtration, in the context of high-depth sequencing samples (14.2 × 10^6^ – 35.9 × 10^6^, median 22.6 × 10^6^) in excluding the anomalies visible in B. The horizontal blue lines indicate the median log_2_ (pseudo)count.

The MA plots of genes called as expressed by all methods are presented in Supplementary Fig. 1; in black we represent boxplots of the differences between expression values of genes for a given abundance (log2 scale); the central black line indicates a 0 log-ratio i.e. identical values in the compared samples; the two blue lines indicate a log2-ratio of ±1. Tight MA plots along the median line indicate consistency in gene expression values between tool combinations (the analysis was conducted per sample; the plots shown correspond to a single sample, randomly chosen). A good concordance was observed between combinations employing similar read quantification tools, i.e. HTseq, featureCounts and GeneCounts, even when a combination with a different alignment tool is considered. When the similarity analysis is restricted to the top 100 most variable genes, based on the sample variance for genes in the pairwise intersection of tool combinations (Jaccard approach), the consistency in response drops (0–5%) as shown in the heatmaps in Supplementary Fig. 2. The similarities between method combinations, as computed for a hierarchical clustering on expression vectors indicate a grouping by read alignment tool for similar read quantification tools, whereas the transcript-level estimation (StringTie) combinations cluster separately. These findings suggest that the choice of read quantification method contributes more to quantified variation in gene expression levels than the choice of read alignment tool. Density plots of gene expression, for each tool combination prior to normalisation (Supplementary Fig. 3) highlights the high shape-similarity of distributions of abundances for all tested combinations. CPM filtering (genes with counts above a threshold proportional to the sequencing depth) marked for exclusion from the subsequent steps of the analysis) was performed to compare the effect of the noise adjustment pipeline. This filtering step reduced the number of expressed genes included in the analysis by approximately 50% (16953–20199, median 19589). Corresponding TMM-normalised and voom-transformed density plots are presented in Supplementary Fig. 5–6 and 7–8, respectively. Representative density plots of unfiltered and CPM-filtered data following TMM-normalisation and voom-transformation are presented in Fig. 2B and 2C, respectively. These analyses indicate that the resulting distributions of gene counts are relatively consistent between methods.

### 3.2 Computational choices influence the number of genes with expression in the user-defined noise range

User-defined thresholds were inferred based on the distribution of point-to-point Pearson correlations versus the abundance of genes, for each sample. An example of three expression profiles, generated from paired pre- and post-vaccination samples so as not to bias for similar conditions, for a high abundance transcript (RPUSD1-201) that would generate a high value for the p2pPCC is shown in Supplementary Fig. 9A; an example of a low abundance transcript from a similar set of samples is shown in Supplementary Fig. 9B (SLC7A9-201), for the latter the distribution of signal across the transcript is not consistent, due to the stochasticity of the sequencing process. A good sample is characterised by high variability in the p2pPCC values for low abundance genes and a consistently high and tight distribution of p2pPCC values for medium and high abundances. An example of a good sample, with a consistently high distribution of correlation values is presented in Fig. 3A; the example presented in Fig. 3B shows a sample with a drop in p2pPCC at the high abundances indicating an inconsistency in distribution of signal that could be due to heavy alternative splicing or significant differential expression of a high number of highly abundant genes; Fig. 3C shows a high proportion of low p2pPCC transcripts over a wide range of medium abundances suggesting the occurrence of a fundamental change in the distribution of signal, across the transcript. The latter can also indicate the presence of technical issues with that particular sample that lead to a generalised pattern. Further investigation into the identity and overarching properties of the transcripts with low p2pPCC value (Fig. 3B and Fig. 3C) revealed little in the way of shared gene ontology between these transcripts.

**Fig. 3.**
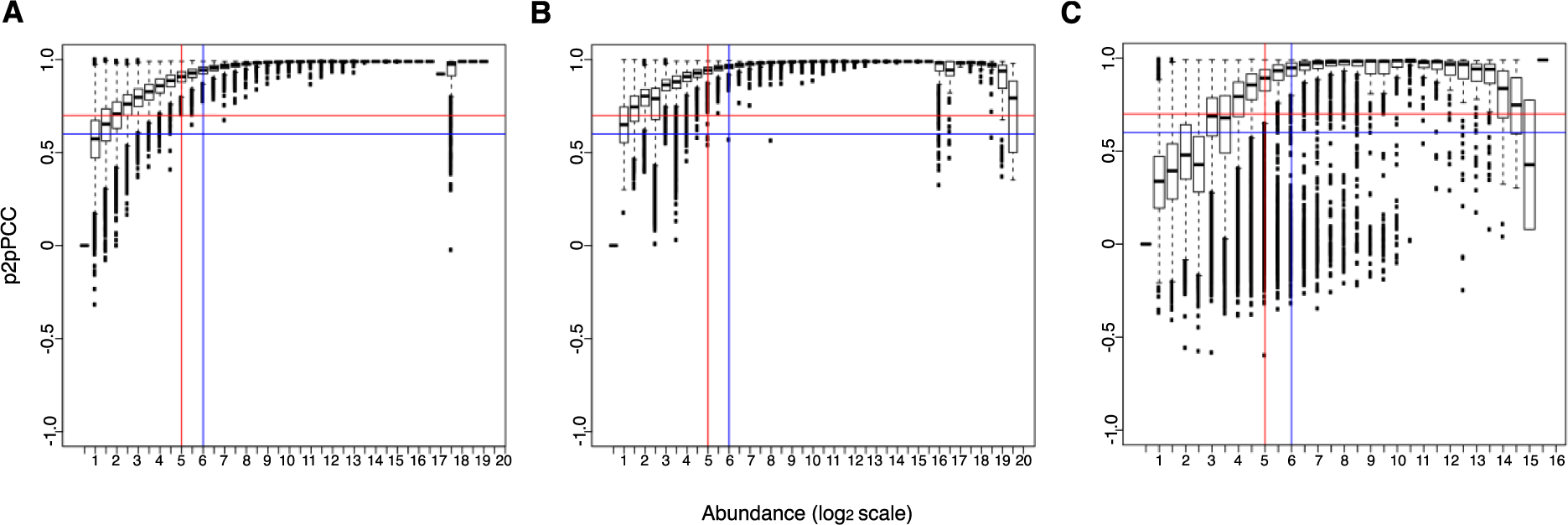
Distributions of point-to-point Pearson correlation coefficient (p2pPCC) against average abundance. In these plots we represent the distributions of p2pPCC values (y-axis) for bins of abundances, on log2 scale (x-axis). **A.** Example of a good sample for which the distribution of p2pPCCs increases steadily, proportional with the increase in abundances to a plateau that is maintained all the way through to the highest abundances. **B.** Example of a sub-optimal distribution of p2pPCCs where the distributions of correlation coefficients drop at the highest abundances indicating either extensive degradation or significant alternative splicing. **C.** Example of a poor distribution of p2pPCCs with numerous outliers across the abundance range. For these examples the signal-to-noise threshold was defined on the abundance bin for which the median of the p2pPCC was above the selected threshold. The red lines indicate a lower user-defined noise threshold, defined as the abundance at which the median of the p2pPCC distribution exceeds 0.7. The blue lines indicate a higher user-defined noise threshold, with a user-defined threshold of 0.9. Both thresholds are “first-occurrence” i.e. the first abundance for which the distribution of p2pPCCs is above the user-defined threshold. The implemented method provides the option of “last-occurrence” threshold i.e. the last abundance for which the distribution of p2pPCC is below the user-defined threshold.

The proportions of genes in the noise range are consistent between alignment tools (18581–25357 [median 20317] and 18619–25096 [median 20362] for HISAT2 and STAR alignments, respectively) but vary depending on the quantification tool used with approximately 34% of genes falling in the noise range for StringTie (25357 for HISAT2-StringTie and 25096 for STAR-StringTie combinations) and ∽56% (28960) when using the STAR --quantMode GeneCounts function (see Fig. 4A). This observation indicates that more computational noise is introduced by the choice of the gene/transcript estimation tool than by the choice of read alignment tool. Density plots of genes with expression levels below the user-defined noise threshold are shown in Fig. 4B. The expression values of “noisy” genes were increased to the signal-to-noise threshold, corresponding to each sample to reduce the number of DE calls for genes with values below the noise threshold in a subset of sample. Genes with all expression values below the noise thresholds, for each sample, were excluded from the analysis (see Supplementary Fig. 10). Noise adjustment reduced the number of expressed genes included in the analysis by 5.1–40.6% (22627–35902 genes remaining, median 33108), depending on the relative proportion of noise associated with a particular tool combination, with the remaining lowly expressed genes treated in a sample-specific manner. The resultant noise-adjusted, *voom*-transformed count density plot for noise adjusted and above-the-threshold genes is shown in Fig. 4C. The full set of density plots for all noise-adjusted, *voom*-transformed samples and tool combinations, with and without prior CPM-filtration, are presented in Supplementary Fig. 11 and 12, respectively. The former (Supplementary Fig. 11) exhibits greater variation in between density peaks below the median across all samples due to sample-specific noise adjustment, while in the latter (Supplementary Fig. 12) these differences are less apparent; the density distributions are nicely aligned above the median for both methods.

**Fig. 4.**
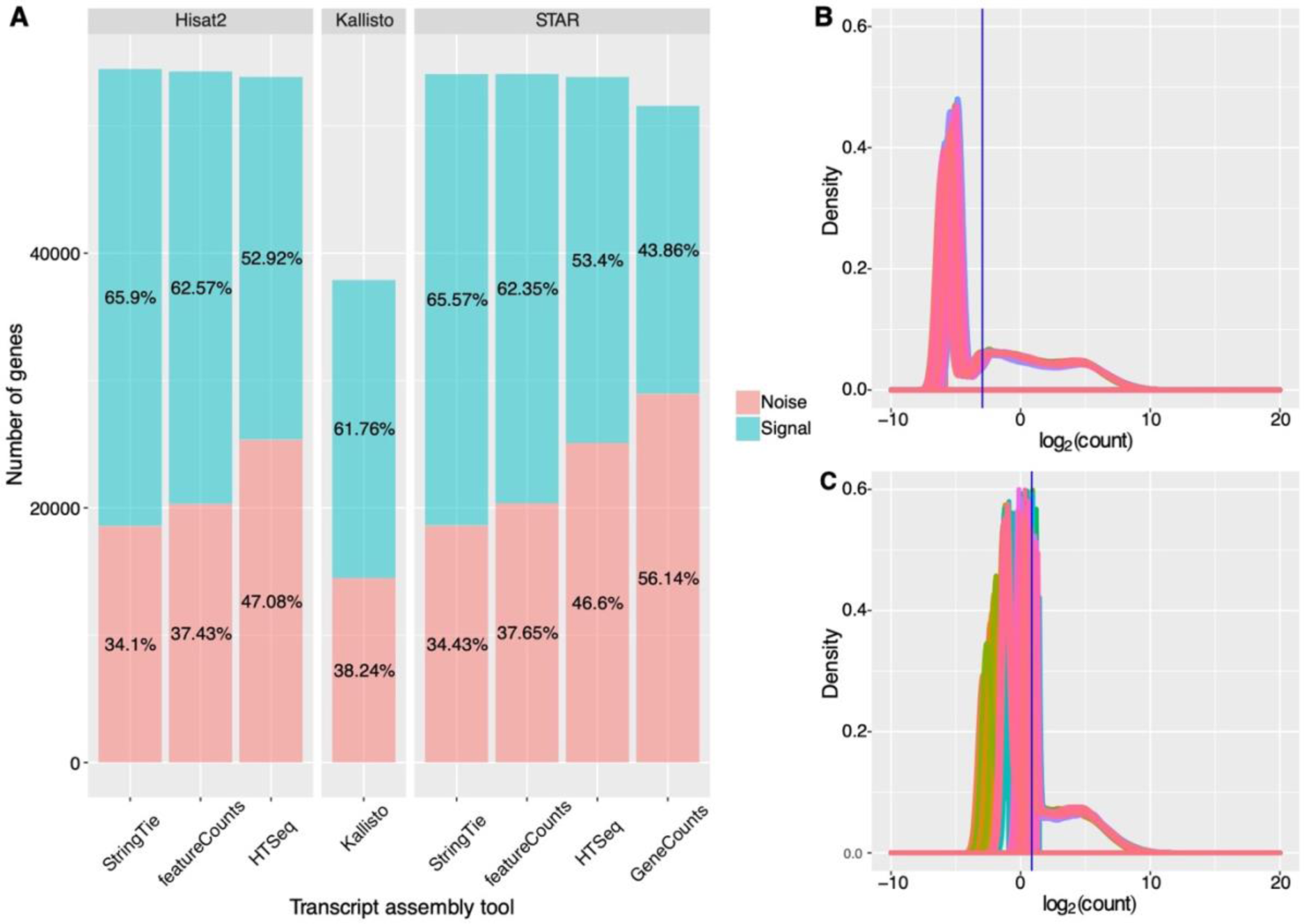
Effect of noise thresholds on the number and distribution of abundances of the selected genes. **A.** Stacked bar plot showing the relative proportion of genes with expression levels above/below the signal-to-noise for each read alignment/quantification tool combination. Proportions are presented as percentages of total expressed genes for each tool combination. StringTie generates a lower number of entries for the noisy range (18851 and 18619 for HISAT2 and STAR combinations, respectively), whereas HTseq and GeneConts produce a higher number of genes with low, and highly variable abundances (for the latter >50% of expressed genes) **B.** Density plots of median gene abundances for genes with abundances below the user-defined noise threshold for a single tool combination (HISAT2-StringTie). The expression values of these genes are subsequently increased to the sample specific signal/noise threshold and genes with expression values less than the signal-to-noise thresholds in all samples are removed. **C.** Density plot of noise-adjusted, *voom*-transformed read counts for the same tool combination (HISAT2-StringTie). The vertical blue lines indicate the median log_2_ count across all samples.

### 3.3 Removing genes in the noise range reduces the effect of pipeline-specific differences on the final set of differentially expressed genes

MA plots generated after the DE call were generated for each tool combination (Fig. 5A–D and Supplementary Fig. 13), with/without CPM-filtration and with/without noise-adjustment. CPM-filtration without noise-adjustment (Fig. 5B) gave rise to a greater number of low abundance, high LFC outliers – genes with highly variable distribution of signal across the transcript and for which the calculated LFC on the low number of incident reads does not reflect a transcript of biological relevance. These outliers (artefacts of differential expression analyses derived from low abundance transcripts) were absent from the corresponding noise-adjusted MA plot and reduced the call as DE by 250–1168 (median 509) transcripts, with LFC values varying between −1.016653 and 6.717757 (|median| 2.31) and abundances in the 1.3261–74120.038 (median of medians: 30.9832) range. The CPM-filtered, noise-adjusted method gave rise to the same distribution of abundances as the unfiltered, noise-adjusted MA plot, suggesting that the noise-adjustment performs equally well on filtered and non-filtered data. The noise-adjusted MA plots showed a more consistent distribution of abundances across the entire range compared with their noise-included counterparts, highlighting a high degree of variability at the lower abundances. This suggests a converging effect of the noise adjustment on a variety of analytical backgrounds. The pipeline was applied to a single combination of tools (STAR-quantMode GeneCounts) for the *M. musculus* dataset, as the primary focus was to assess the impact of analytical noise on the final DEG call. 19,898 genes included in the final analysis following noise removal, compared with 15,096 genes following CPM filtration. The density plots generated for this tool combination yielded similar distributions to those of the *H. sapiens* tool combination counterpart (Supplementary Fig. 14).

**Figure 5.**
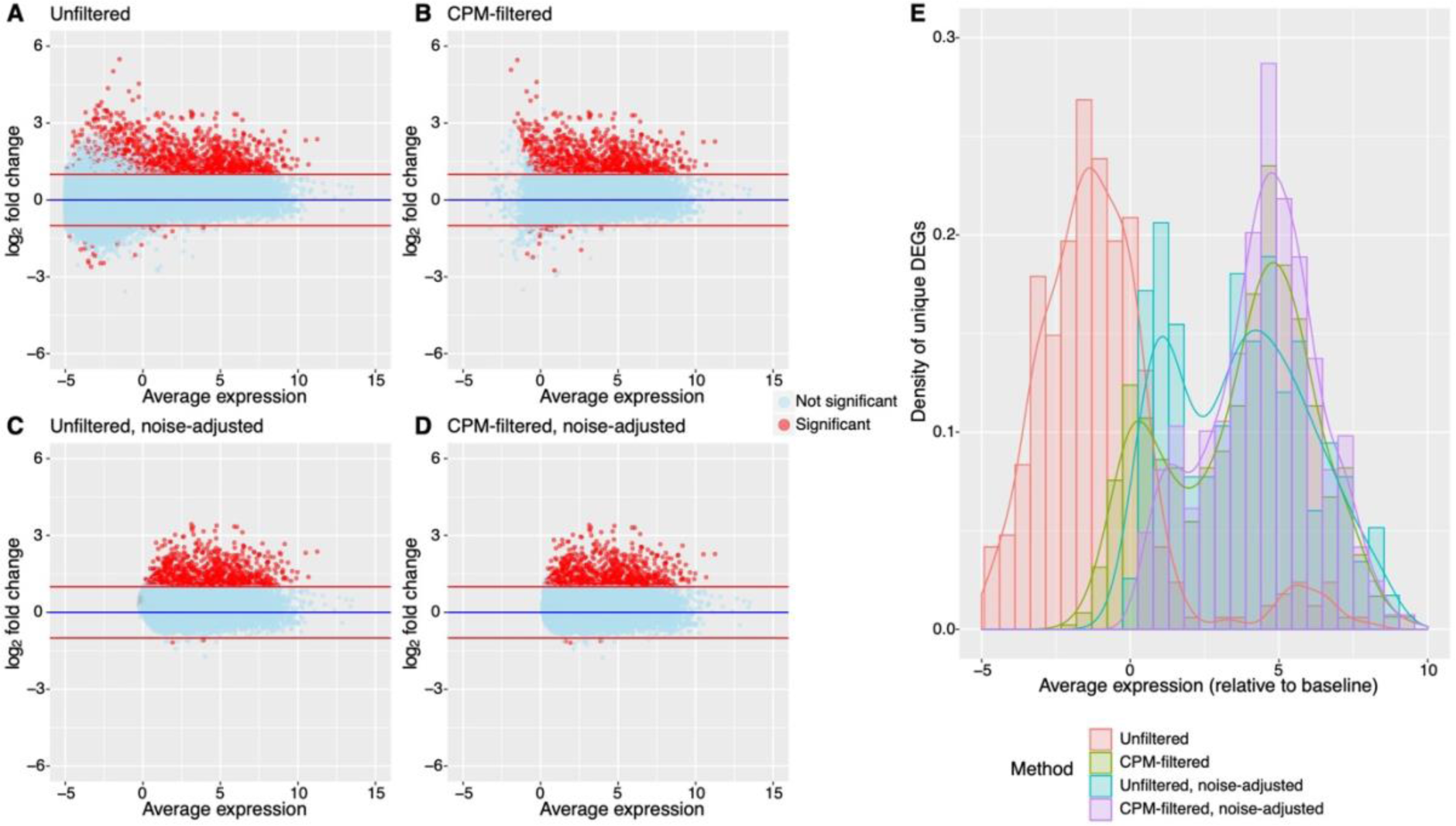
Distributions and properties of genes called differentially expressed subsequent to each filtration method. The MA plots show the pairwise comparison between the pre-vaccination and the post-vaccination samples; the significantly differentially expressed genes (DEGs, FDR < 0.1, absolute LFC > 1) are highlighted in red. The input for each MA plot is: **A.** unfiltered input, **B.** CPM-filtered input, no noise adjustment, **C.** unfiltered, noise-adjusted, and **D.** CPM-filtered, noise-adjusted. For each plot, the horizontal blues line corresponds to a log_2_ fold change of 0; the horizontal red lines correspond to a log_2_ fold change of +/- 1. Due to the nature of the compared datasets and the significant biological changes occurring after vaccination, the predominant identification of up-regulated genes is correct. **E.** Density histograms of average expression of genes uniquely called as significantly differentially expressed (DE) for each filtration method on the same tool combination. This plot indicates that the unfiltered data leads to more DEGs with low expression values being called significantly DE compared with the same data with CPM-filtered and/or noise adjusted data. CPM-filtered, noise-adjusted data call fewer lowly expressed genes as significantly DE but capture more significantly DEGs with high average expression values than other methods.

Applying the noise adjustment pipeline to this dataset also confirmed the effects of noise adjustment on reducing the overall DE call by 123 and 1164 transcripts relative to the CPM-filtered data for vaccine one and vaccine two, respectively, by removing low abundance outliers. This resulted in greater consistency between the MA plots (Supplementary Fig. 15).

Finally, histograms of the unique DEGs (FDR < 0.01, |LFC| > 1) associated with each filtration method for each tool combination revealed a low density of overlapping medium abundance genes between the CPM-filtered methods (see Fig. 5E and Supplementary Fig. 16). Conversely, the unfiltered methods had a higher density peak associated with low-abundance genes (see Supplementary Fig. 16). These plots highlight the specific differences remaining between the different tool combinations following DE analysis.

## 4 Discussion

This analysis highlights the specific differences between eight different read processing pipelines (as combinations of frequently used pre-processing tools) also in relation to empirically-determined noise thresholds; it also illustrates the effect of a novel data-driven filtration method for the removal of outlying low abundance, highly variable genes. While most studies comparing RNA-seq analysis pipelines have focused on benchmarking the normalisation and statistical testing of DE *(6, 21, 28, 29)*, we elected instead to focus on the joint effects sequencing noise and of analytical noise derived from alignment and count estimation methods on DE calls. A previous study evaluating the impact of alignment methods on the DE analysis, without taking into account the presence or extent of technical noise, explored these methods in combination with a single read quantification tool *(30)*. We have shown that a vast proportion of variability in the final output introduced by computational choices is attributable to the read quantification tool. This effect may be explained by the handling of multi-mapping reads and assessment of transcript variants identified in the alignment stage. However, it is encouraging to observe that a data-driven filtering of genes with inconsistent distribution of signal, reduces the effect of tool choice.

We chose to conduct our analysis on an experimentally-generated dataset in *H. sapiens* in which both experimental groups were given a highly immunogenic stimulus that would cause a significant perturbation in the host transcriptome. The high variability between individuals observed in *H. sapiens* allowed us to evaluate the robustness of the noise detection method on samples likely to exhibit considerable intra- and inter-condition inconsistency. The usefulness of this method to identify sequencing and analytical noise was further tested on a *M. musculus* dataset where inter-individual variability is minimized (the data was generated from inbred mice of the same sex). The tests on this, more robust, data emphasized the impact on the DE call introduced by noise; the noise adjustment approaches performed equally well in this scenario. The choice of two vaccines of differing immunogenicity, and therefore likely to induce very different levels of transcriptional activity, also confirms that the pipeline performs well in scenarios where DE is highly variable between experimental conditions. For both datasets, the performance of our data-driven method was compared with the *ad hoc* fixed CPM filtration procedure (applied uniformly across samples) that has been suggested for RNA-seq DE analysis *(22, 23)*. While this method is practical for bulk-excluding low abundance genes, the criteria for filtration should be sample specific; it has been demonstrated that filtering methods that make use of group conditions, such as minimum sample size of a test group, may increase the true false positive rate in the downstream in the analysis *(31)*, this effect becomes particularly visible with the increase in complexity of the experimental design of the dataset. While the noise adjustment pipeline performed similarly on unfiltered and CPM-filtered data, we observed that the noise adjustment of unfiltered data results in a higher number of uniquely significantly DEGs that have low average expression values compared with those uniquely identified by noise adjustment of CPM filtered data. However, both methods demonstrated an improvement on the CPM filtration method.

The analysis is provided as a standalone pipeline, accessible via a wrapper script that invokes all the components, in the correct order and with a seamless linking of parameter transfer; the speed of the processing is achieved via a *divide-et-impera* split of the initial dataset that allows the execution of this pipeline without an extensive memory or disk usage. The individual scripts that make up this pipeline are accessible and could be replace by synonymous components e.g. that make use of alternative alignment tools. One limitation of the current implementation of our novel noise detection pipeline is its use of the slow PatMaN algorithm *(32)* chosen for its high mapping accuracy but at the cost of prolonging the alignment time. However this step of the pipeline can be adapted for other deterministic alignment tools to reduce the trade-off between accuracy and time. Additionally, the implementation of the noise thresholds in the DE analysis is thus far only adapted to the *limma/voom* method. This is due to difficulties that are encountered with adjusting the functions for methods that employ scaling-based normalisation methods with negative binomial distributions *(33, 34)*.

## Conclusion

The effect of the sequencing and analytical noise may change the list of DE genes and impact the biological interpretation of experiments. By quantifying and excluding sequencing noise, and choosing appropriate tools for alignment and quantification, we show on a H sapiens and a M Musculus example that the output of different pipelines of calling DE can converge.

## Acknowledgements

We acknowledge the suggestions and numerous discussions with our colleagues from the bioinformatics section at the Oxford Vaccine Group, Department of Paediatrics, University of Oxford, UK.

## Supporting information

**S1 File. Supporting file.** Supplementary file.

**S1 Data. Supplementary Fig. 1**

**S2 Data. Supplementary Fig. 13**

**S1 Table. Supplementary Table 1.**

**S2 Table. Supplementary Table 2.**

