## Supplementary material for "Effects of technical noise on bulk RNA-seq differential gene expression inference": S1 File. Supporting file. Supplementary file.

**Supplementary Fig. 1.** (Attached as a separate PDF) **MA plots comparing the expression levels of genes called by the different genome alignment/read assembly tool combinations.** We compare the log ratio (M) and mean average (A) values between each pair of combinations for a single sample (XXX). The boxplots of the differences between expression values of genes for a given abundance are represented in black. The central red lune indicates a 0 log-ratio, i.e. identical values in the compared samples. The two blue lines indicate a log-ration of + or - 1. This analysis illustrates the variability specific to the analytical noise and the deviation from what should be a standard non-DE example.

**Supplementary Fig. 2.** Heatmaps showing the absolute and relative variation in expression levels of the top 100 most variable genes for different choices of genome alignment/read assembly tools for **A)** sample 1 and **B)** sample 2. The variability is measured using the standard deviation between the 15 measurements. The expression levels are represented on a $\log_2$ scale. The clustering of the method combinations is represented by the dendrograms on the x-axis. The $\log_2$ expression values and corresponding normalised count values associated with each colour of the heat gradient are depicted in the legend in the upper left corner. The grouping indicates that the alignment method drives the differences observed between the tool combinations. These observed similarities are consistent between samples.

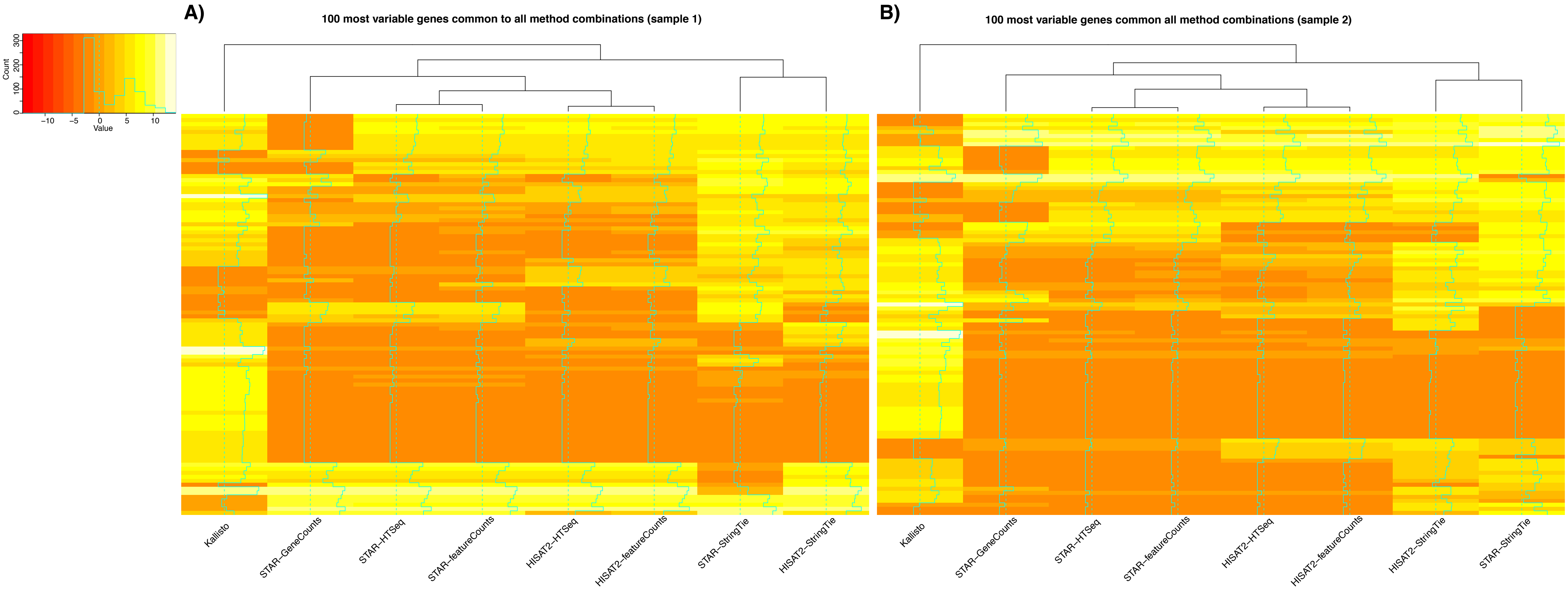


**Supplementary Fig. 2.**

**Supplementary Fig. 3.** Read count density plots of gene expression levels for each combination of genome alignment/read assembly tools derived from the unfiltered count matrices prior to TMM normalisation. The median is represented by the vertical blue line. For all samples, and all combinations of tools, the distributions of abundances are similar.

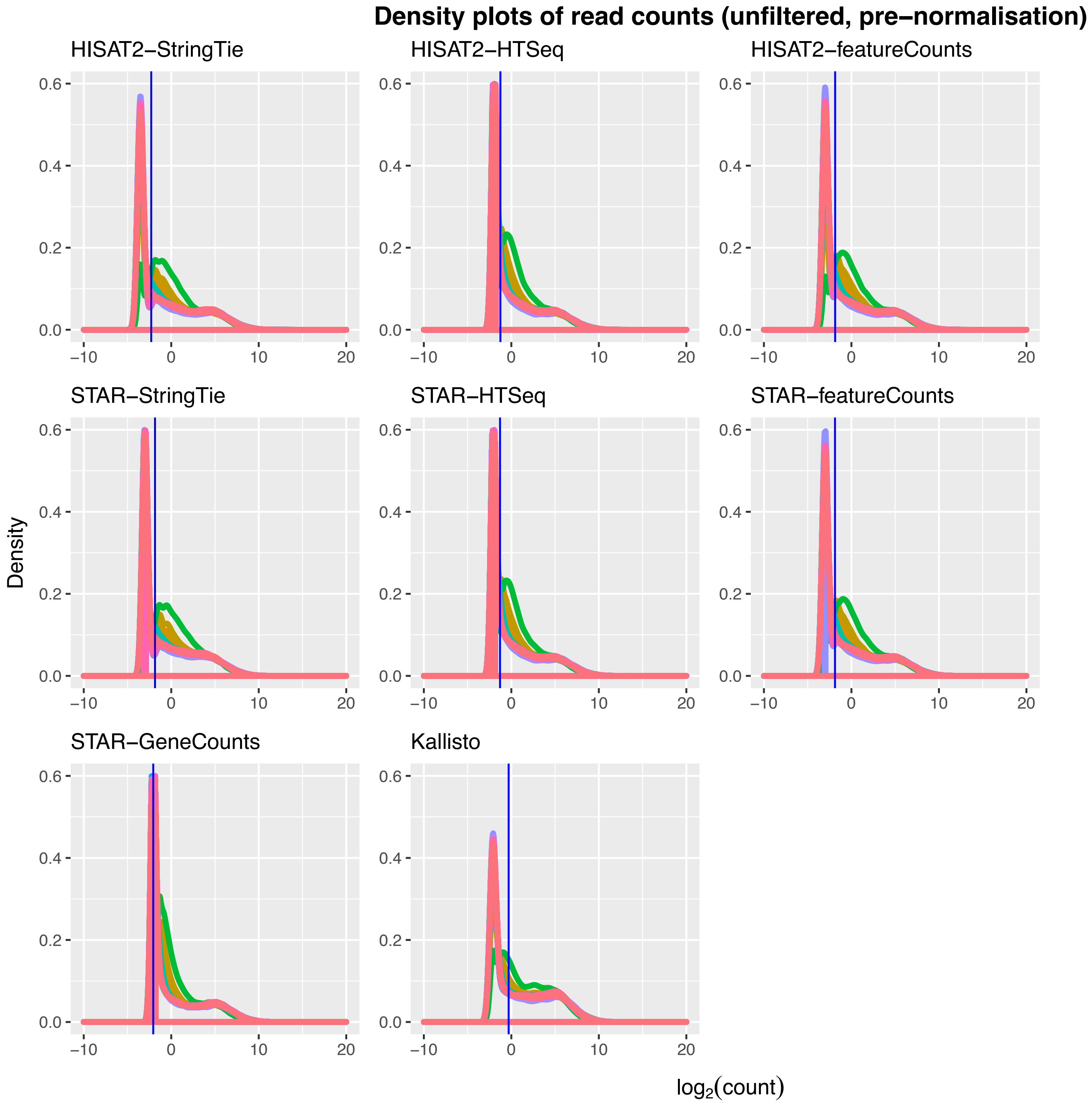


**Supplementary Fig. 3.**

**Supplementary Fig. 4.** Read count density plots of gene expression levels for each combination of genome alignment/read assembly tools following CPM filtration prior to TMM normalisation. The median is represented by the vertical blue line. For all samples, and all combinations of tools, the distributions of abundances are similar.

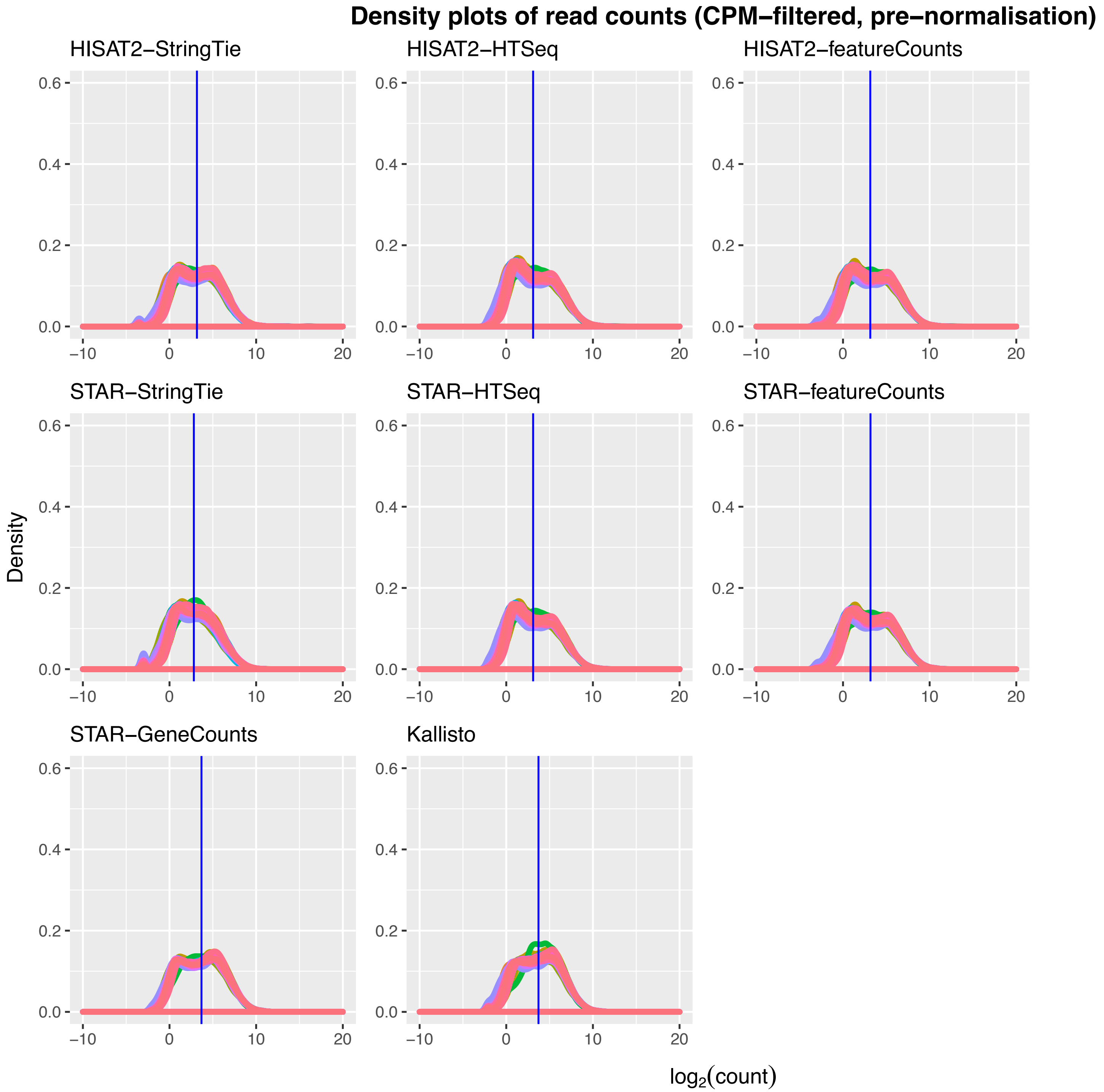


Supplementary Fig. 4.

**Supplementary Fig. 5.** Read count density plots of gene expression levels for each combination of genome alignment/read assembly tools from unfiltered count matrices following TMM normalisation. The median is represented by the vertical blue line.

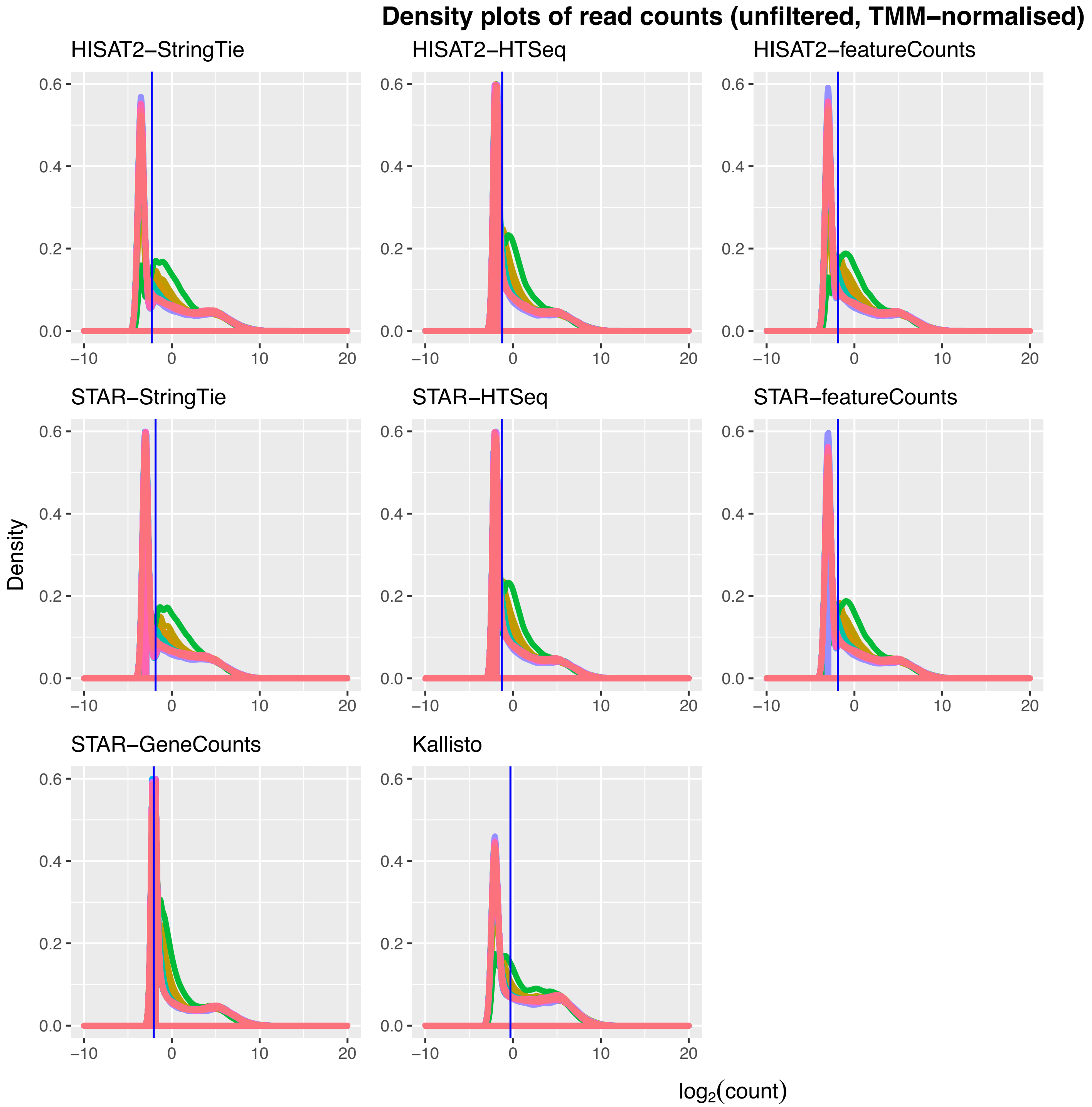


**Supplementary Fig. 5.**

**Supplementary Fig. 6.** Read count density plots of gene expression levels for each combination of genome alignment/read assembly tools following CPM filtration and TMM normalisation. The median is represented by the vertical blue line.

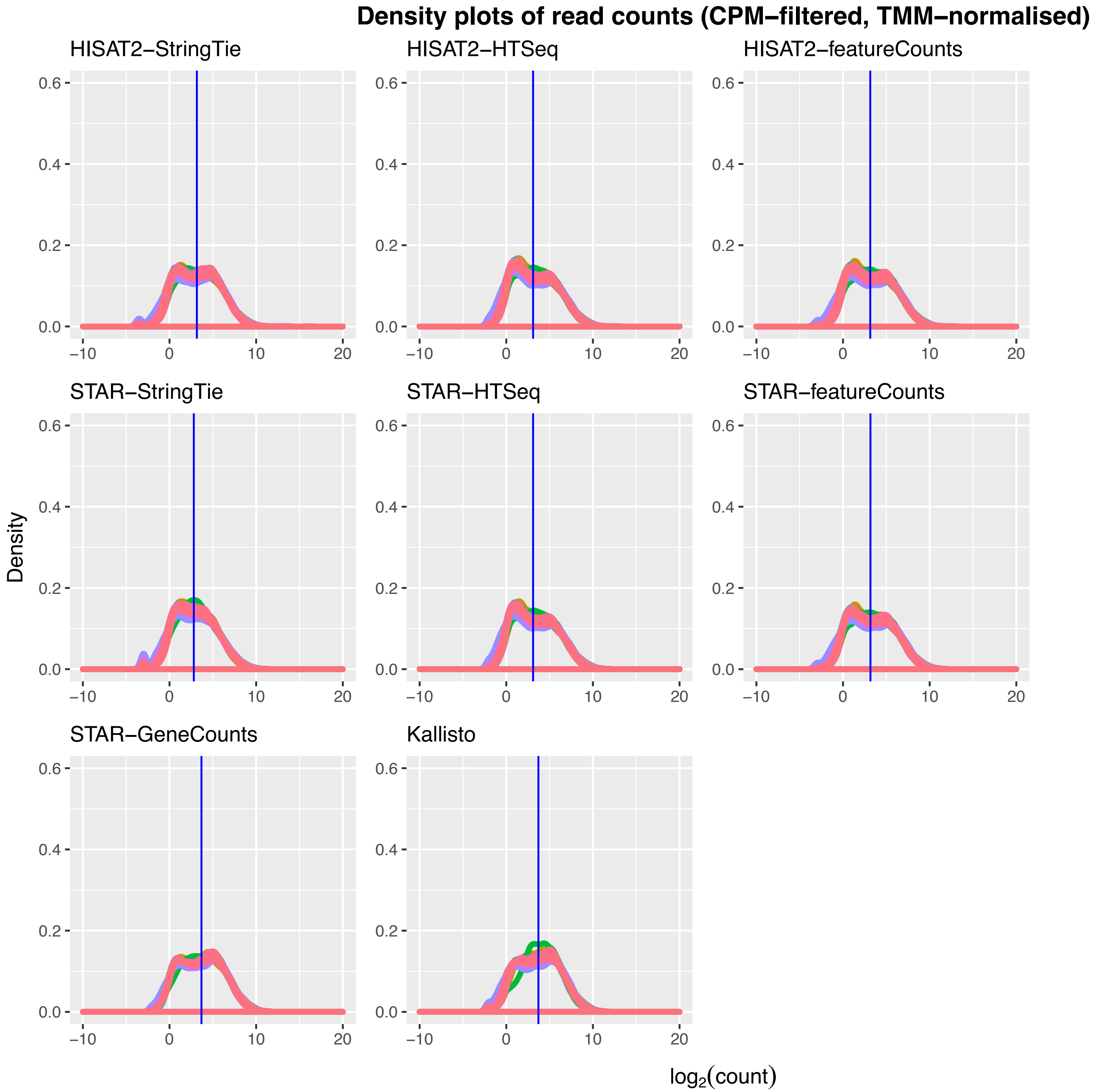


**Supplementary Fig. 6.**

**Supplementary Fig. 7.** Read count density plots of gene expression levels for each combination of genome alignment/read assembly tools, unfiltered, following TMM normalisation and *voom* transformation. The median is represented by the vertical blue line.

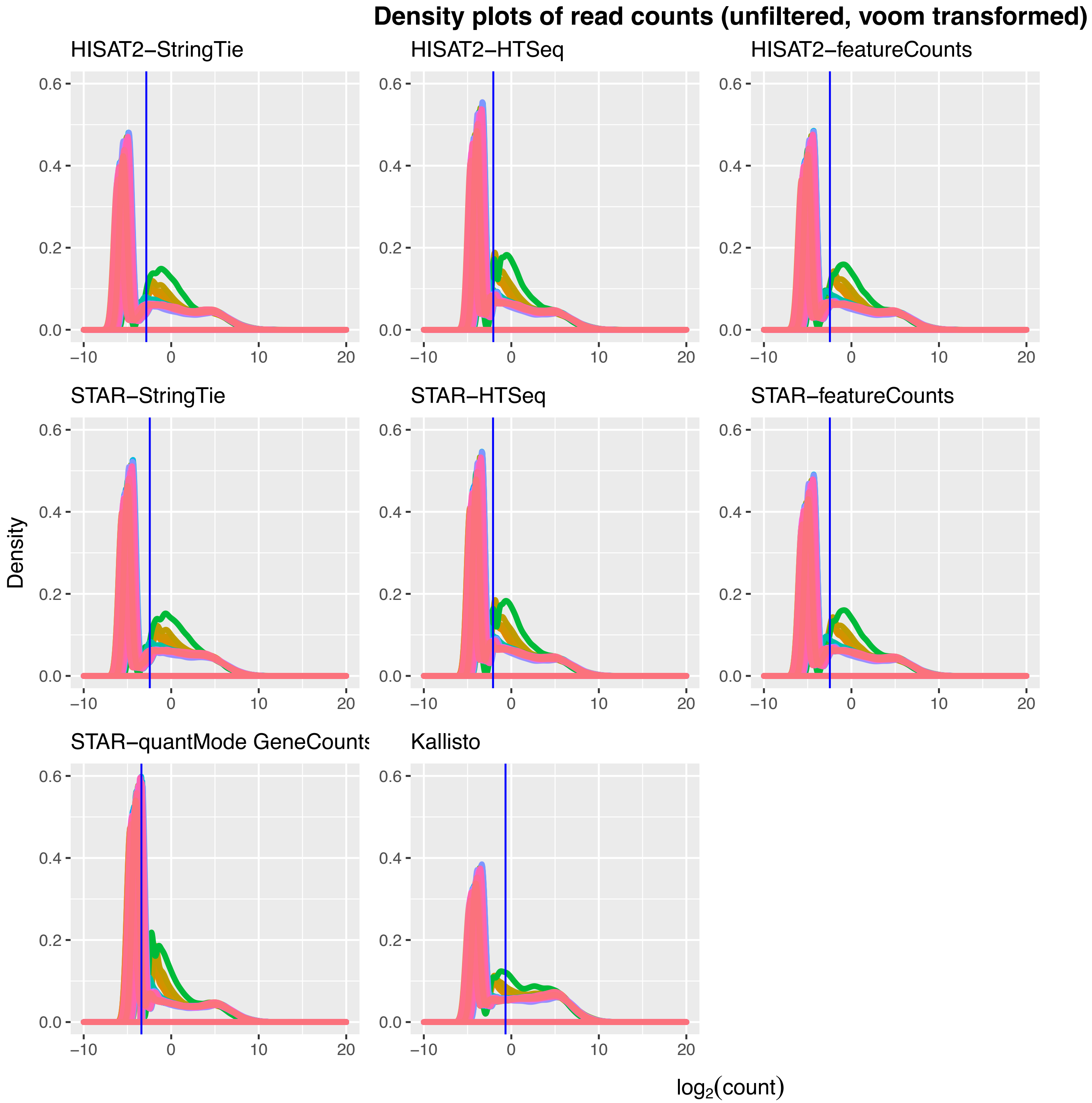


**Supplementary Fig. 7.**

**Supplementary Fig. 8.** Read count density plots of gene expression levels for each combination of genome alignment/read assembly tools, CPM-filtered, following TMM normalisation and *voom* transformation. The median is represented by the vertical blue line.

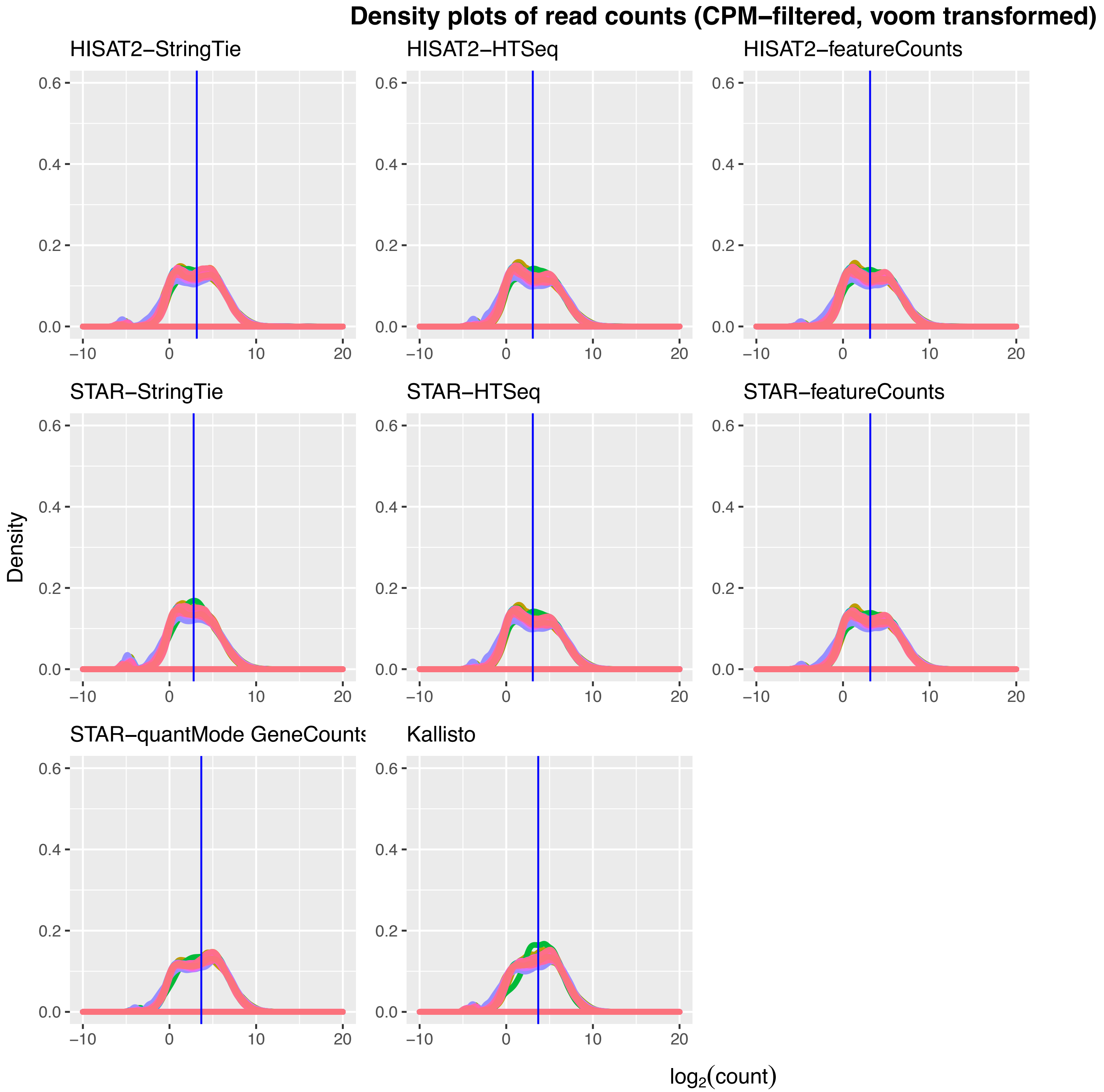


**Supplementary Fig. 8.**

**Supplementary Fig. 9.** Examples of expression profiles for **A.** a high abundance (RPUSD1-201) and **B.** a low abundance (SLC7A9-201) genes, generated from the noise detection pipeline. The expression profiles are built as a point sum across the length of a transcript of the abundances of all reads incident to a particular position.

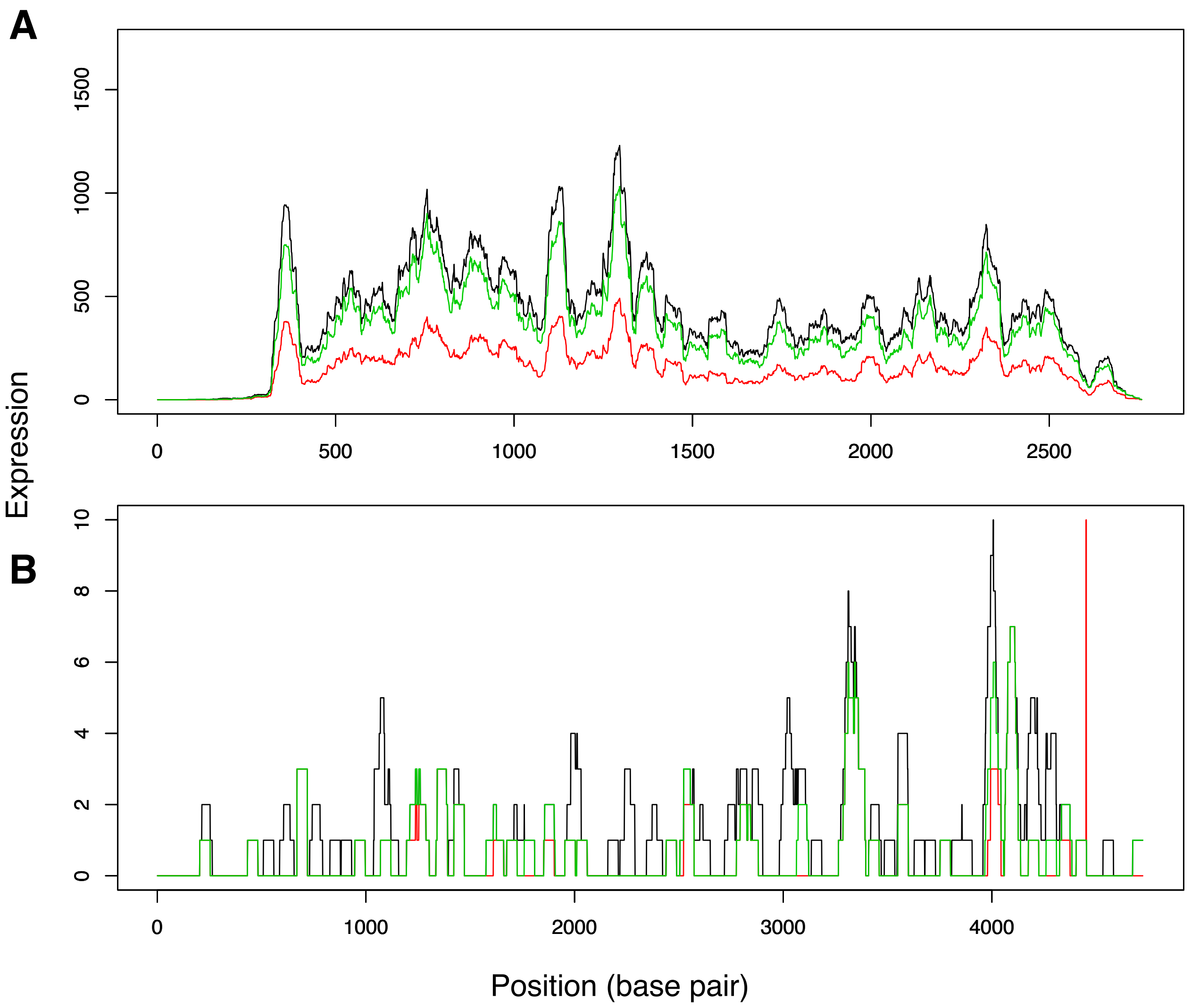


**Supplementary Fig. 9.**

**Supplementary Fig. 10.** Read count density plots of genes with expression levels less or equal to the noise threshold for each sample and each combination of genome alignment/read assembly tools. The median is represented by the vertical blue line.

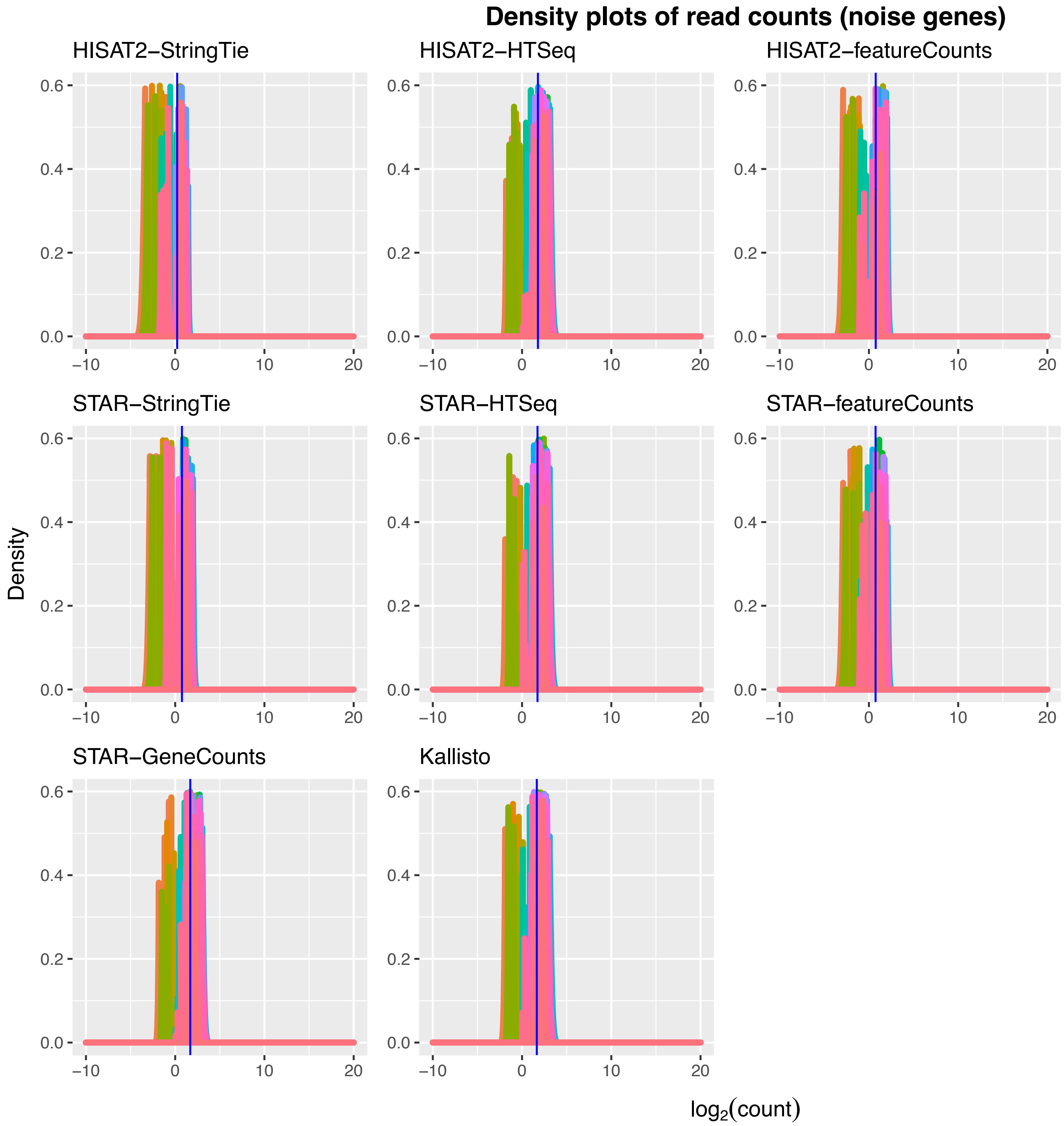


Supplementary Fig. 10.

**Supplementary Fig. 11.** Read count density plots of gene expression levels for each combination of genome alignment/read assembly tools, unfiltered, following TMM normalisation, *voom* transformation and noise adjustment. The median is represented by the vertical blue line.

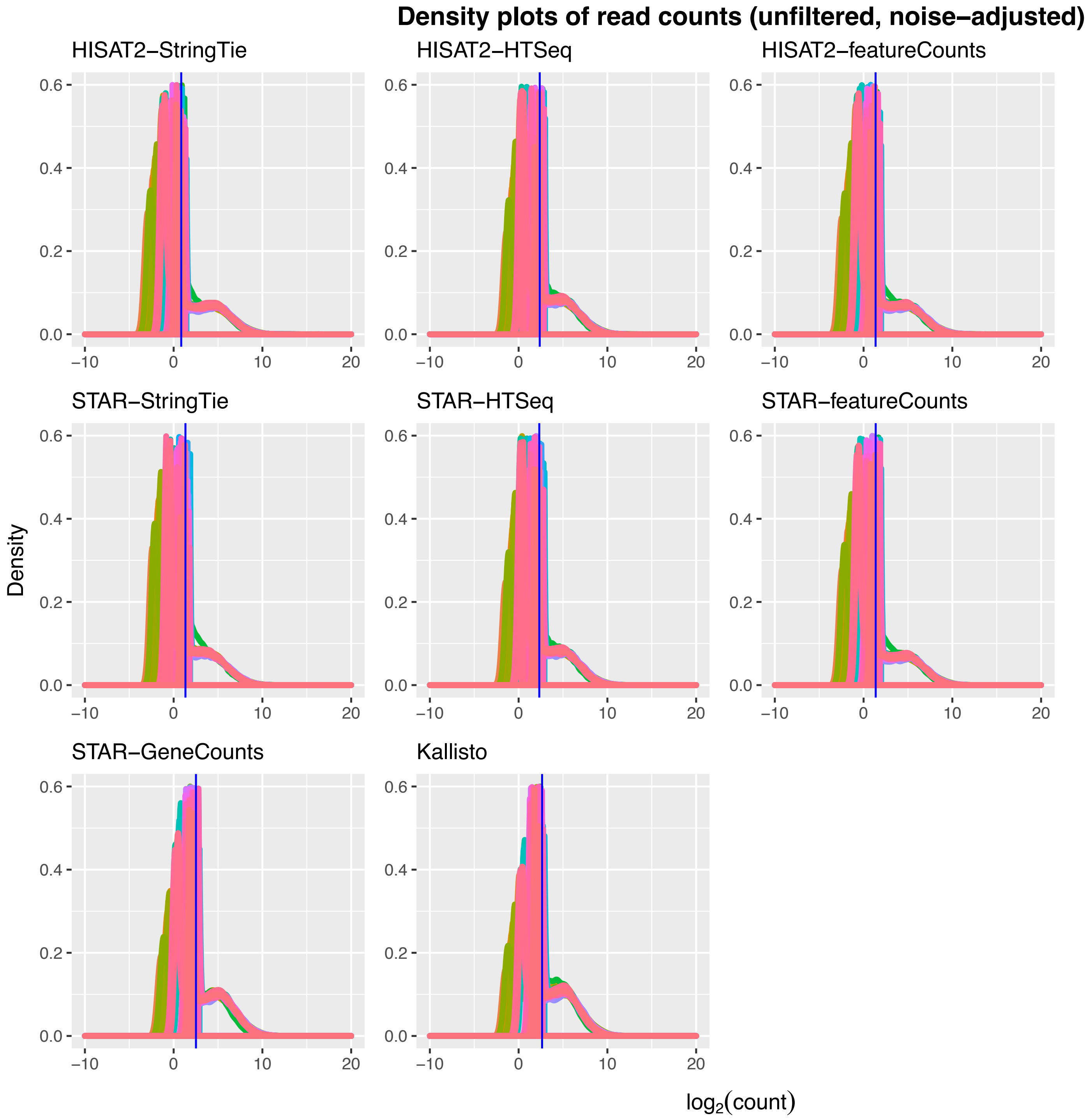


**Supplementary Fig. 11.**

**Supplementary Fig. 12.** Read count density plots of gene expression levels for each combination of genome alignment/read assembly tools, CPM-filtered, following TMM normalisation, *voom* transformation and noise adjustment. The median is represented by the vertical blue line.

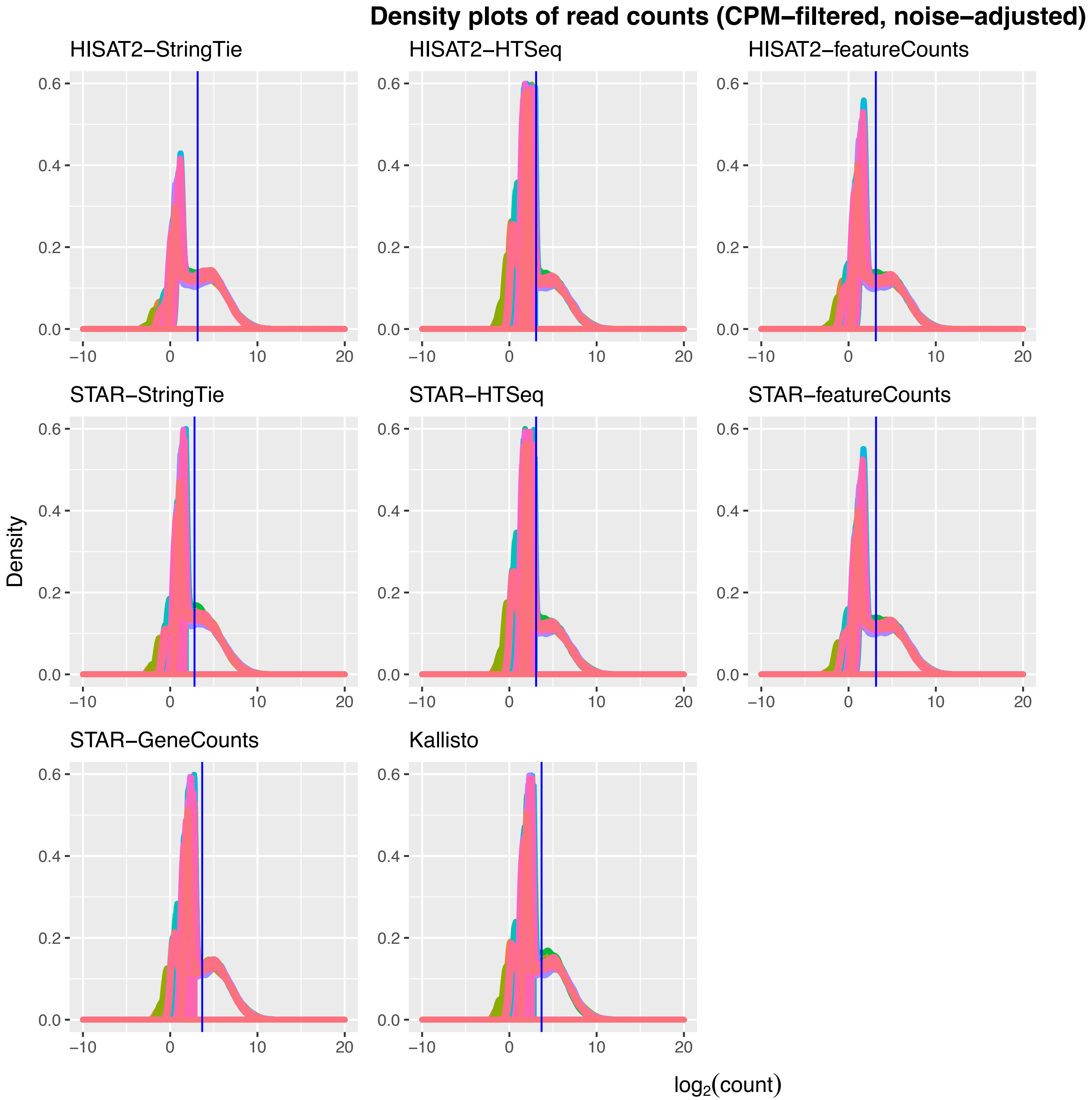


**Supplementary Fig. 12.**

**Supplementary Fig. 13A–H.** **(Attached as separate PDFs)** MA plots of day 1 $\log_2$ fold change (LFC) versus day 0 and mean expression values for control (left panels) and test (right panels) groups, with (bottom panels) and without (top panels) noise thresholds applied for all genome alignment/read assembly tool combinations. The blue lines correspond to an LFC of 0 and the red lines correspond to an LFC of +/- 1. Genes with an absolute LFC of >=1 and an FDR of <0.01 are highlighted in red. The consistent distribution of abundances across the entire range observed for the noise-adjusted MA plots suggests a converging effect of the noise adjustment on a variety of analytical backgrounds.

**Supplementary Fig. 14.** Read count density plots of gene expression levels for each combination of genome alignment/read assembly tools derived from the *M. musculus* dataset at each stage of the pipeline. **A.** Unfiltered, TMM-normalised. **B.** Unfiltered, TMM-normalised, *voom*-transformed. **C.** CPM-filtered, TMM-normalised, *voom*-transformed. **D.** Genes identified in the noise range. **E.** Unfiltered, TMM-normalised, *voom*-transformed, noise removed. **F.** CPM-filtered, TMM-normalised, *voom*-transformed, noise removed.

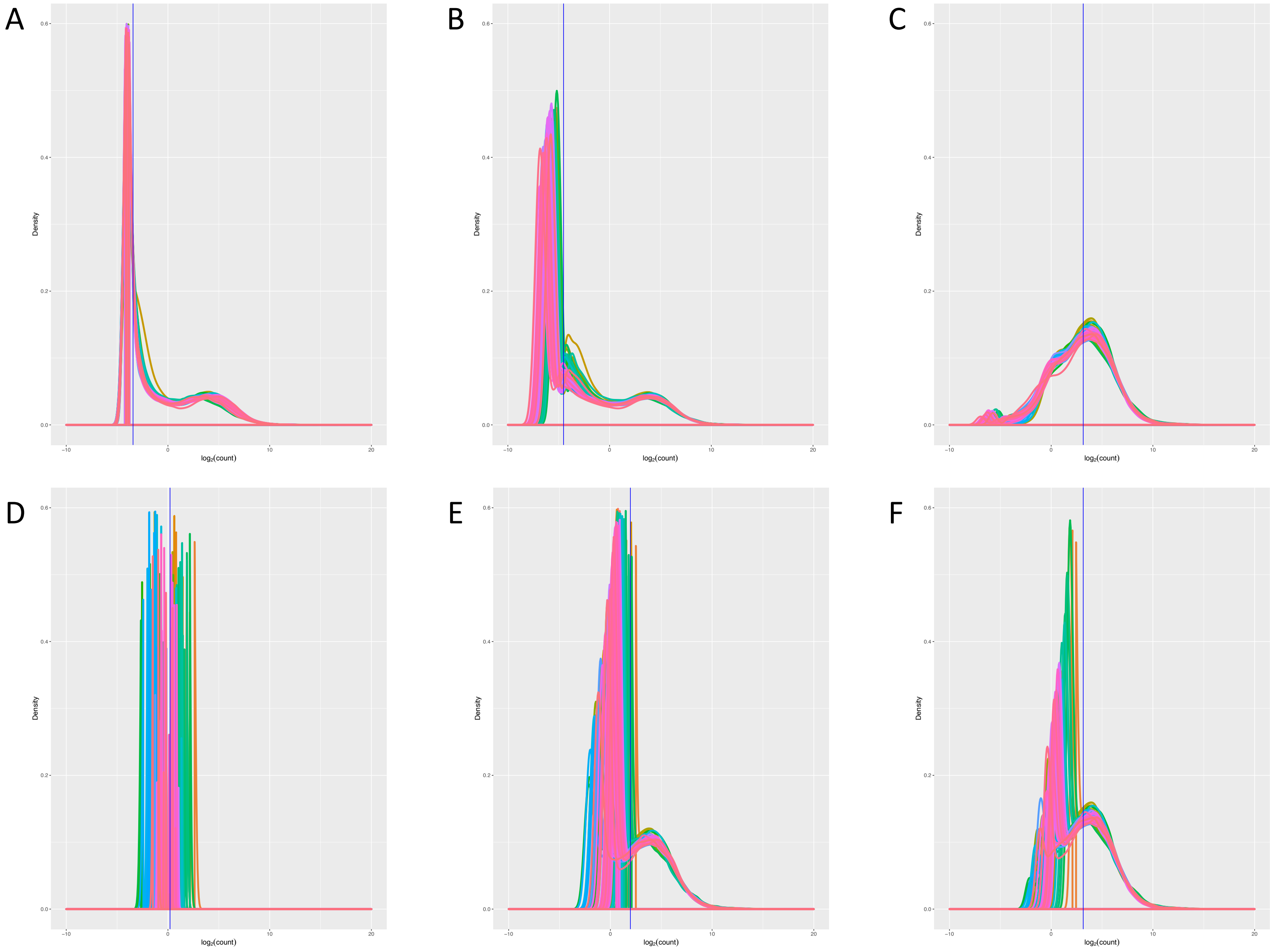


Supplementary Fig. 14.

**Supplementary Fig. 15.** MA plots of day 1 $log_2$ fold change (LFC) versus day 0 and mean expression values for vaccine one (left panels) and vaccine two (right panels) groups, with (bottom panels) and without (top panels) noise thresholds applied for all genome alignment/read assembly tool combinations. The blue lines correspond to an LFC of 0 and the red lines correspond to an LFC of +/- 1. Genes with an absolute LFC of >=1 and an FDR of <0.01 are highlighted in red. The consistent distribution of abundances across the entire range observed for the noise-adjusted MA plots suggests a converging effect of the noise adjustment on a variety of analytical backgrounds.

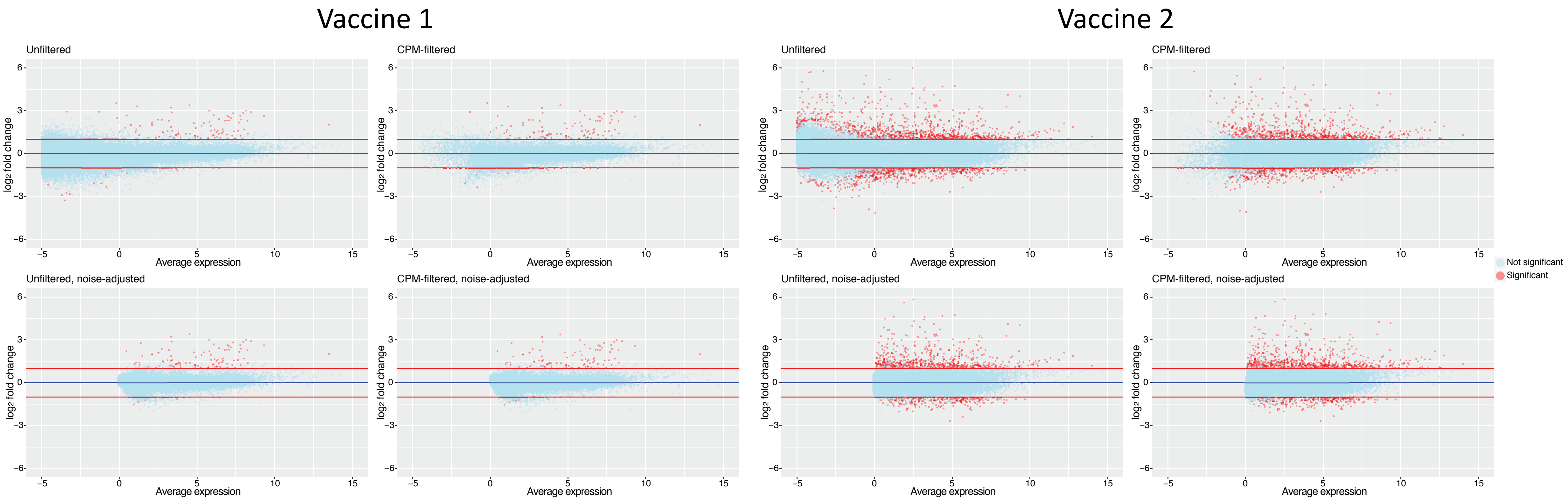


**Supplementary Fig. 15.**

**Supplementary Fig. 16.** Density histograms of the average expression, relative to baseline, of genes uniquely called differentially expressed (false-discovery rate-adjusted $p$-value $< 0.01$ and absolute log fold change $>= 1$) associated with each tool combination, experimental group, and filtration method.

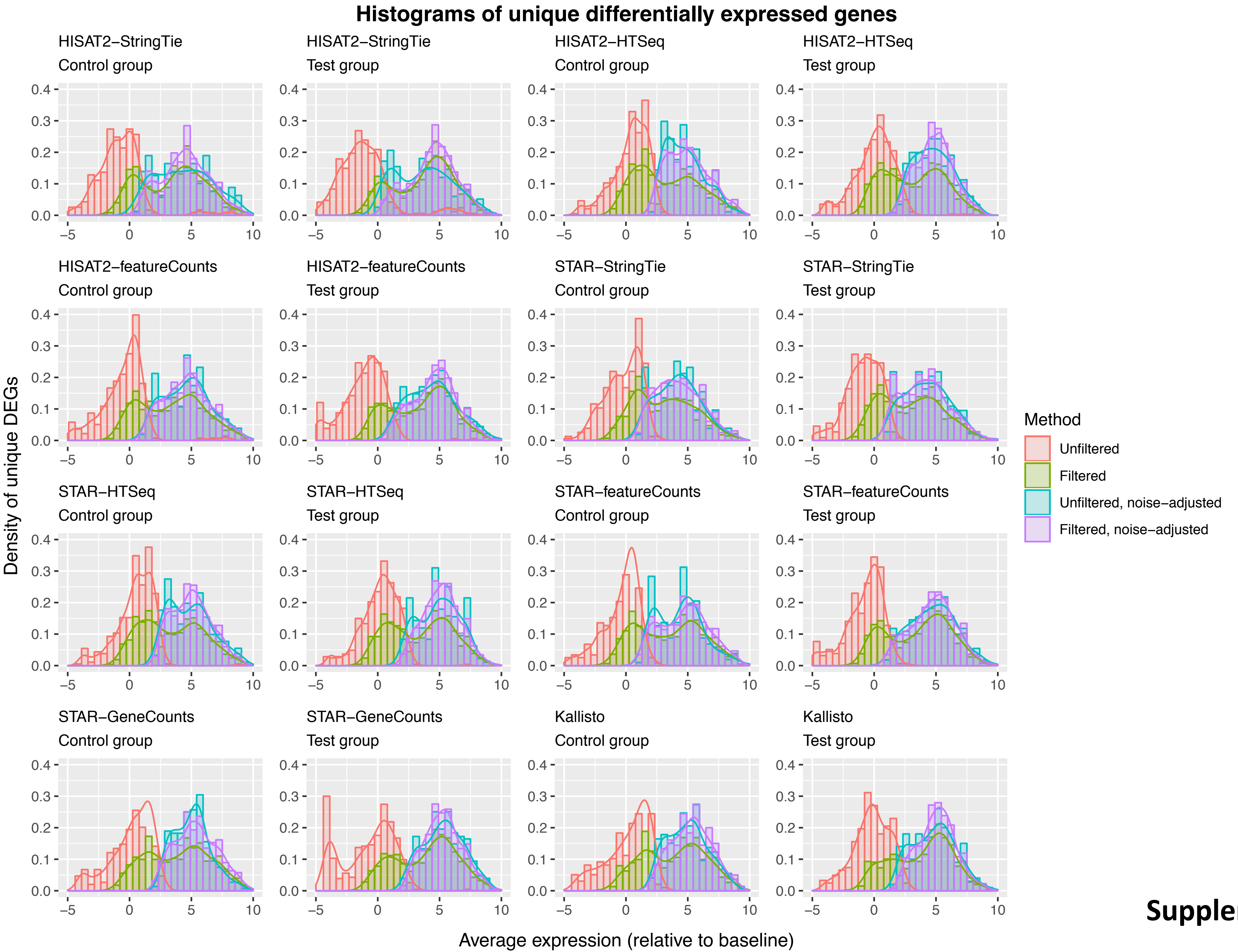


**Supplementary Fig. 16.**
