## Supplementary material for "Effects of technical noise on bulk RNA-seq differential gene expression inference": S1 Data. Supplementary Fig. 1

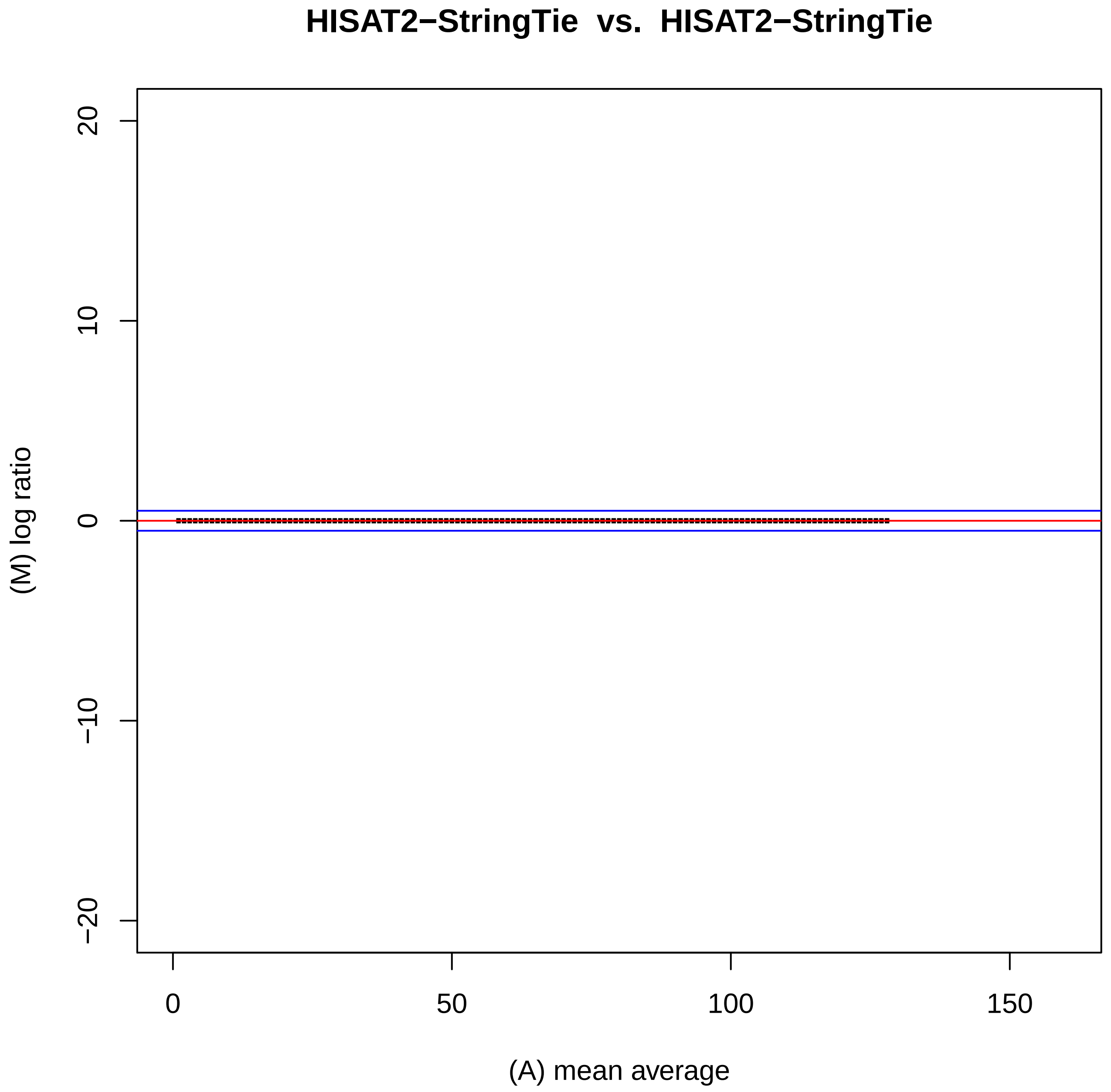
HISAT2−StringTie vs. HISAT2−StringTie
20
10
0
−10
−20
(M) log ratio
0
50
100
150
(A) mean average

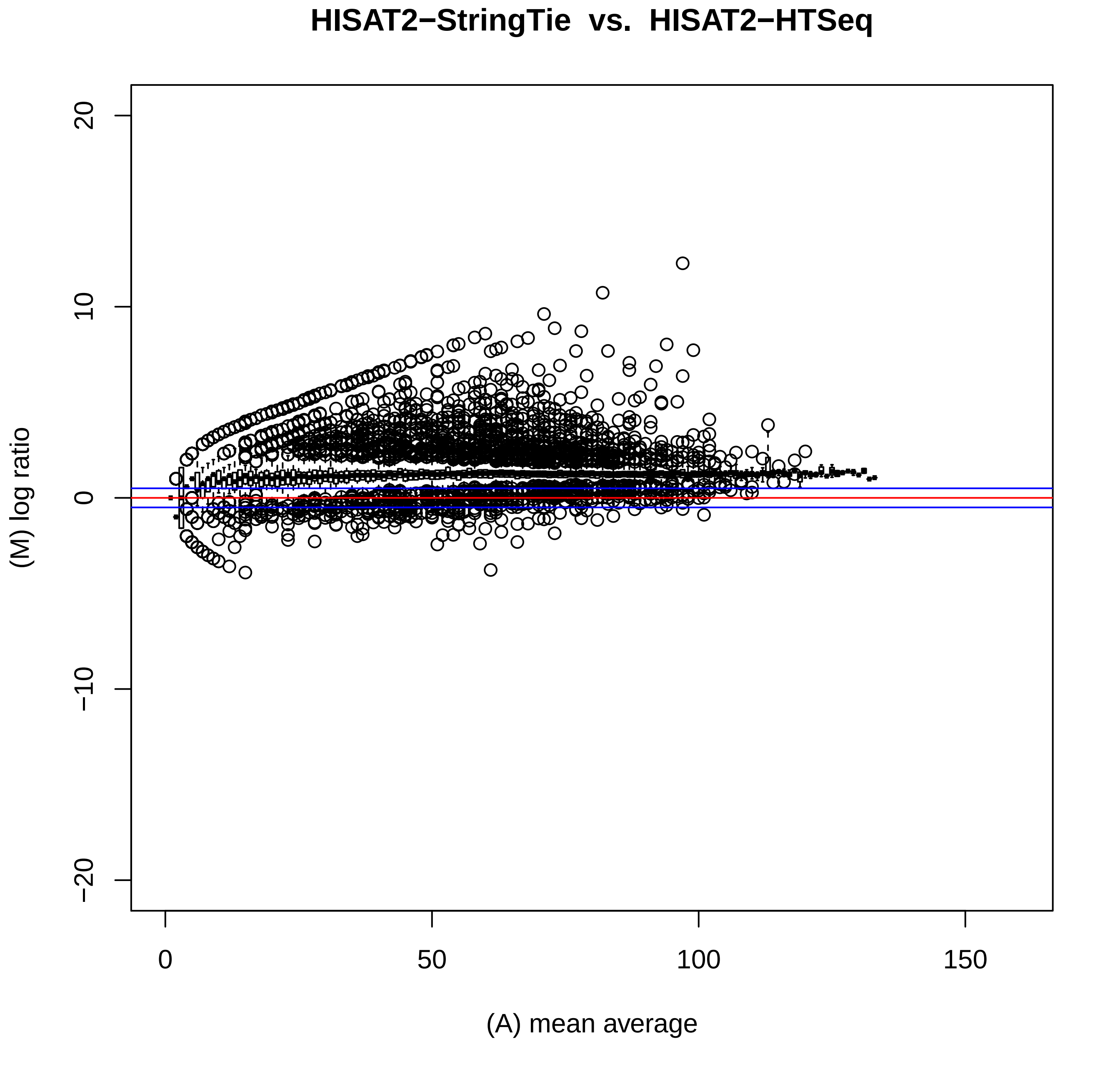
HISAT2−StringTie vs. HISAT2−HTSeq
(M) log ratio
(A) mean average
20
10
0
−10
−20
0
50
100
150

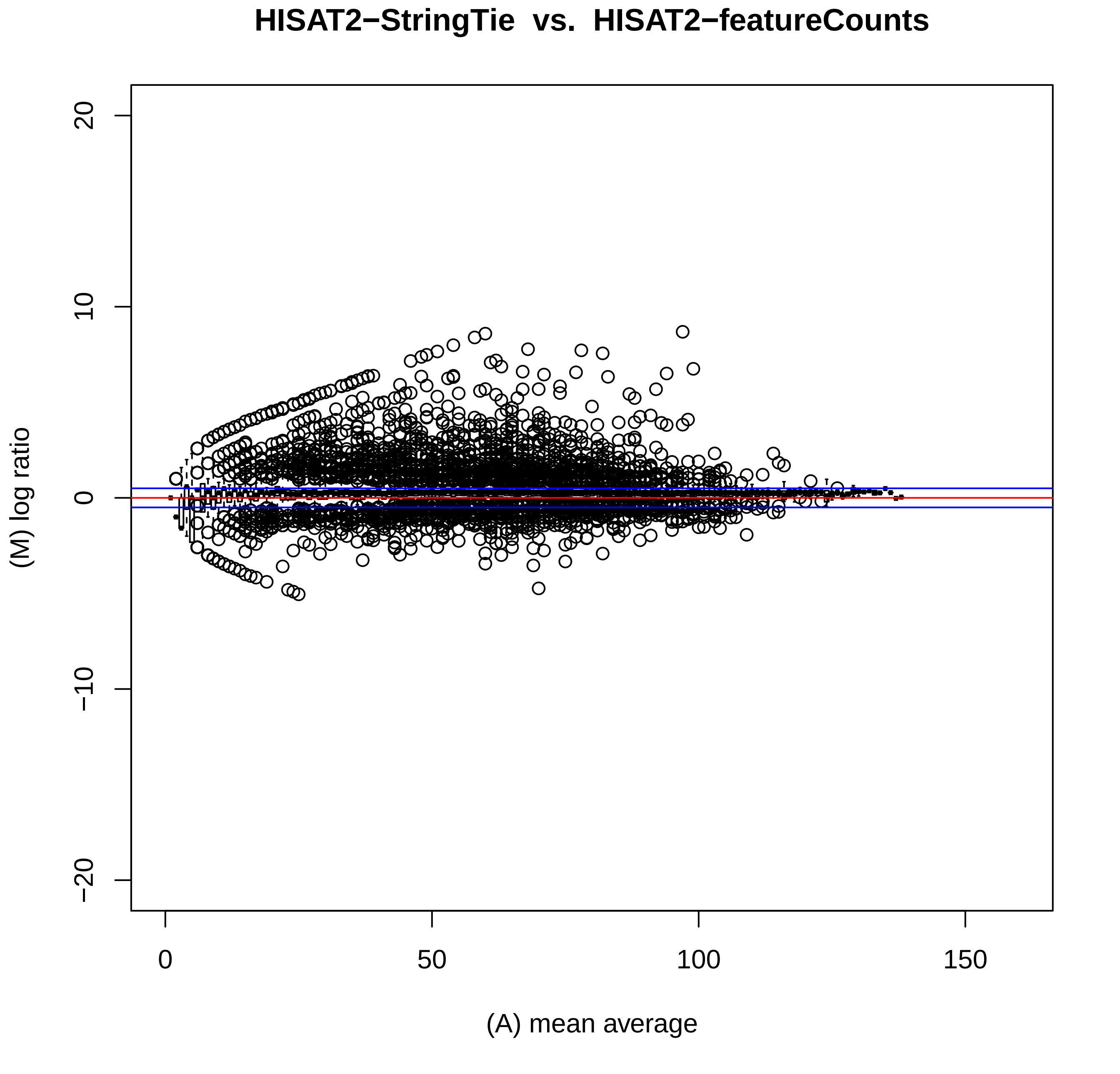
HISAT2−StringTie vs. HISAT2−featureCounts
(M) log ratio
(A) mean average
20
10
0
−10
−20
0
50
100
150

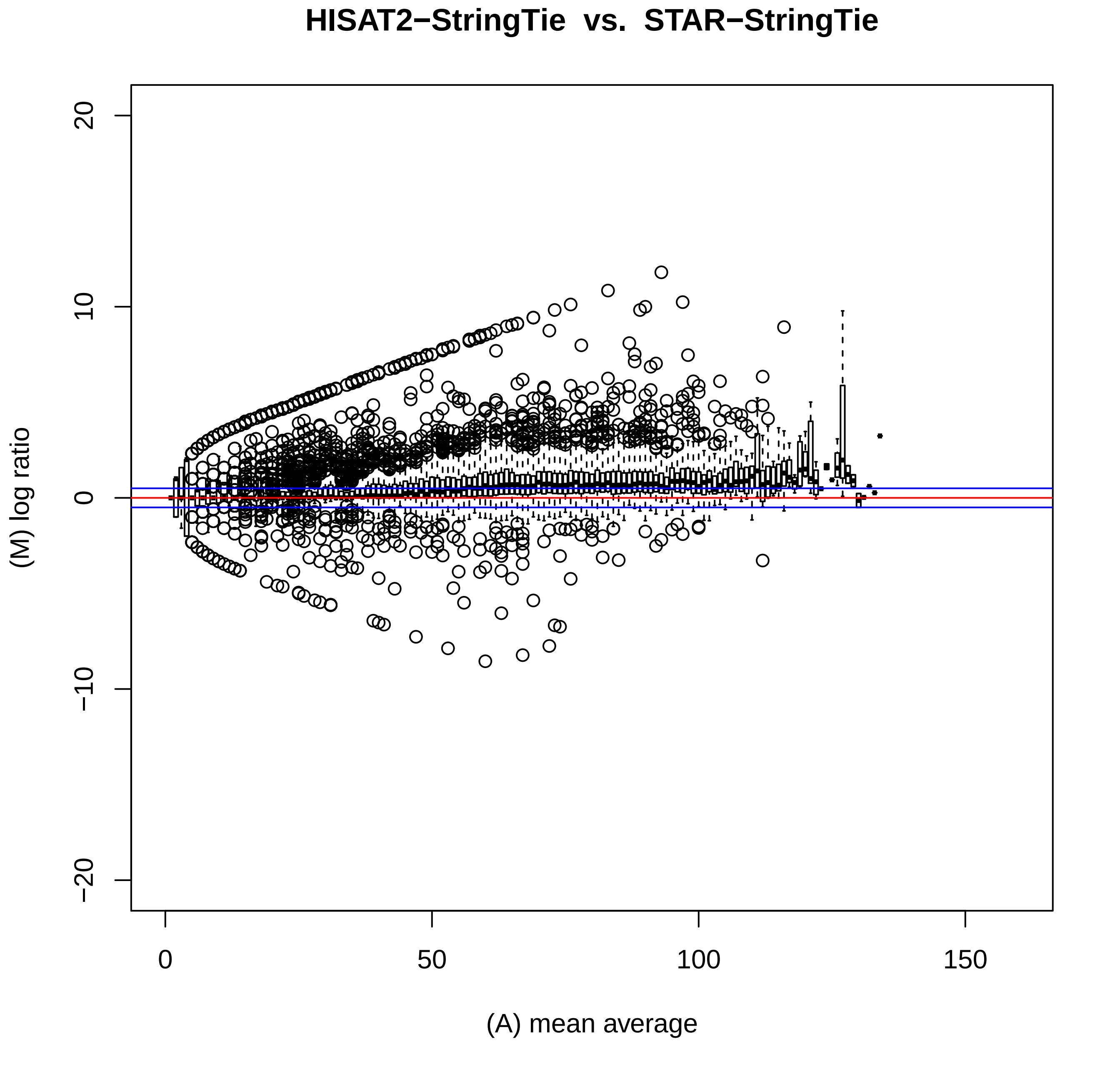

HISAT2−StringTie vs. STAR−StringTie
(M) log ratio
(A) mean average
20
10
0
−10
−20
0
50
100
150

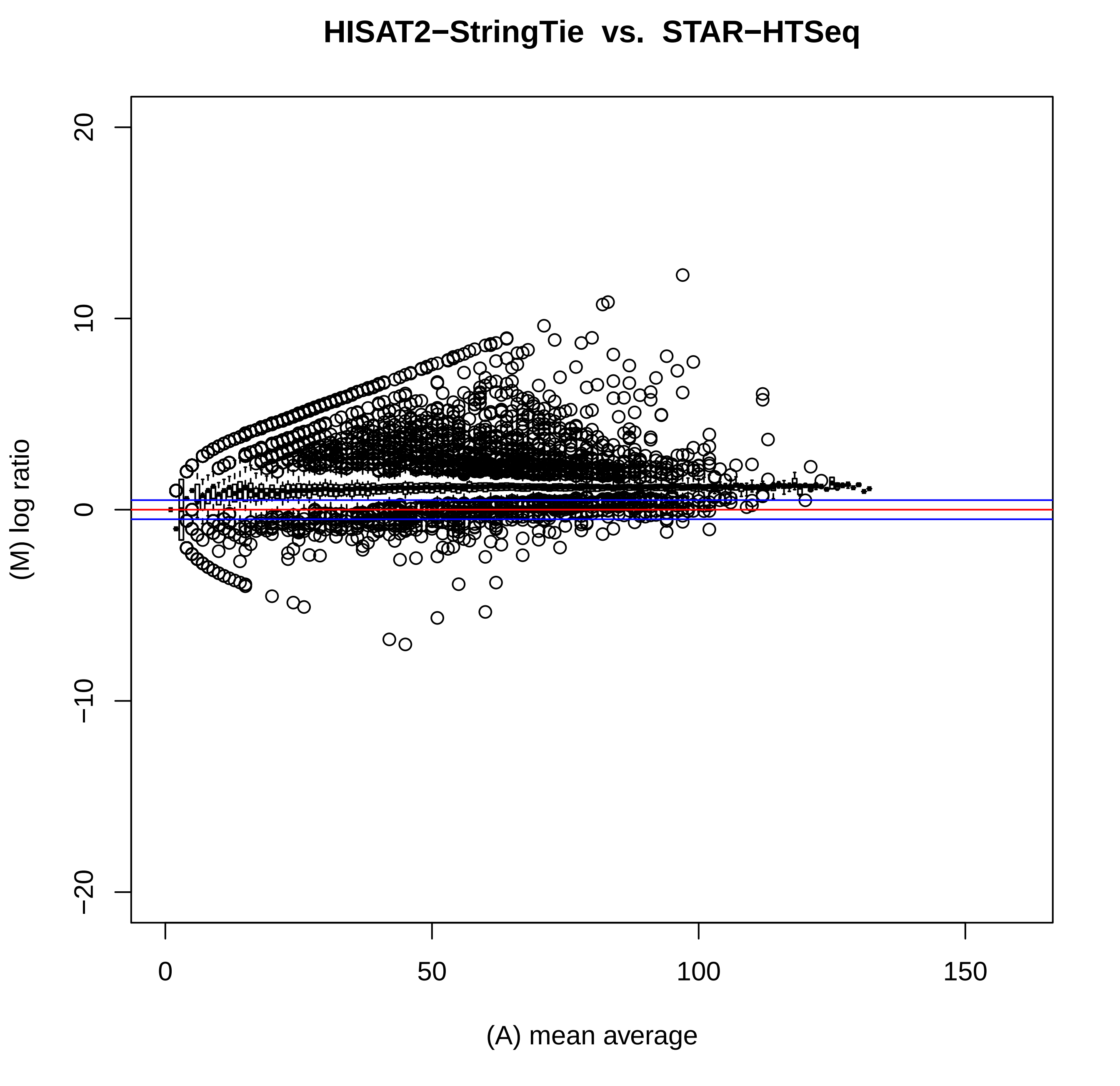
HISAT2−StringTie vs. STAR−HTSeq
(M) log ratio
(A) mean average
20
10
0
−10
−20
0
50
100
150

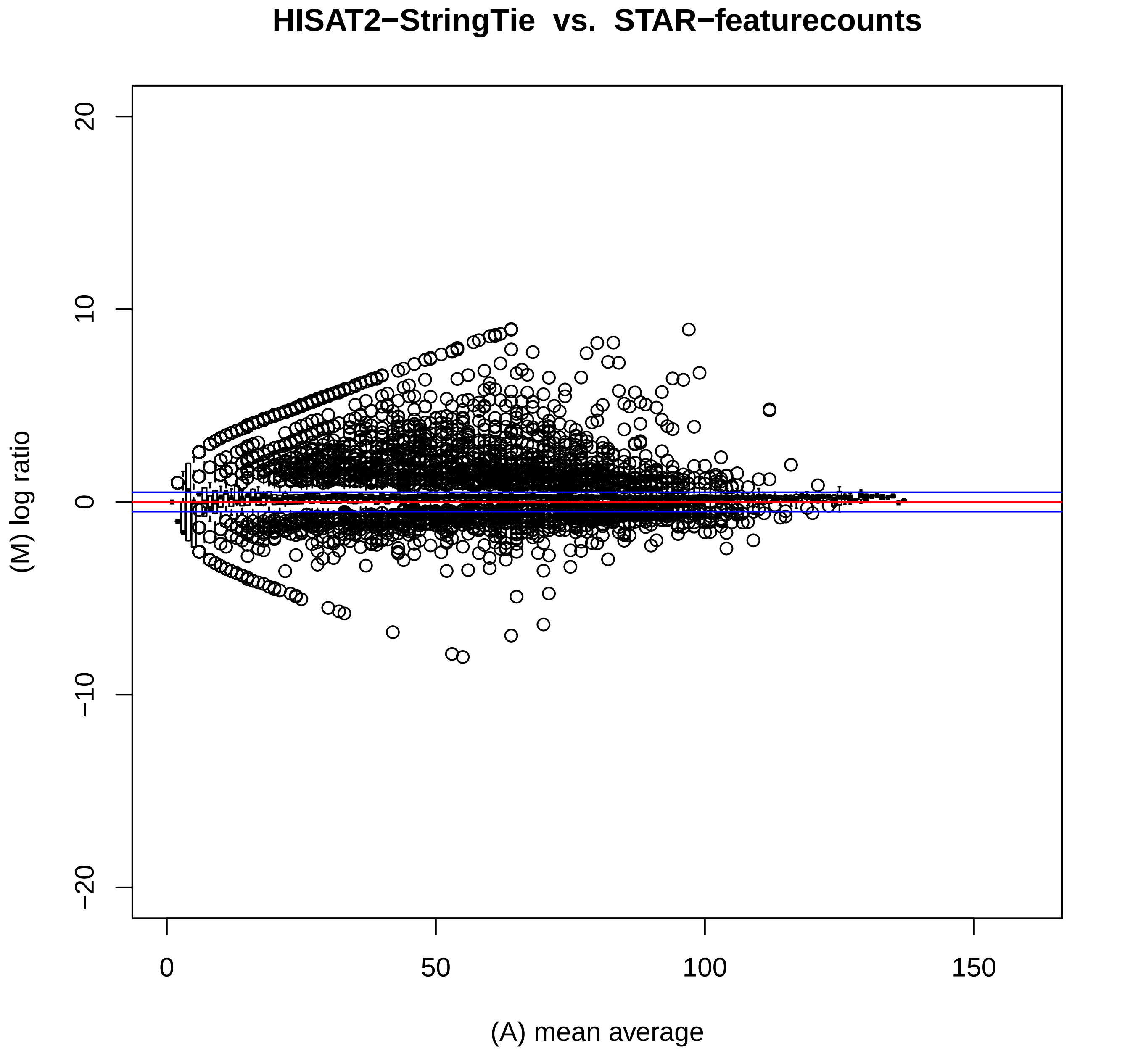

HISAT2−StringTie vs. STAR−featurecounts
(M) log ratio
20
10
0
−10
−20
(A) mean average
0
50
100
150

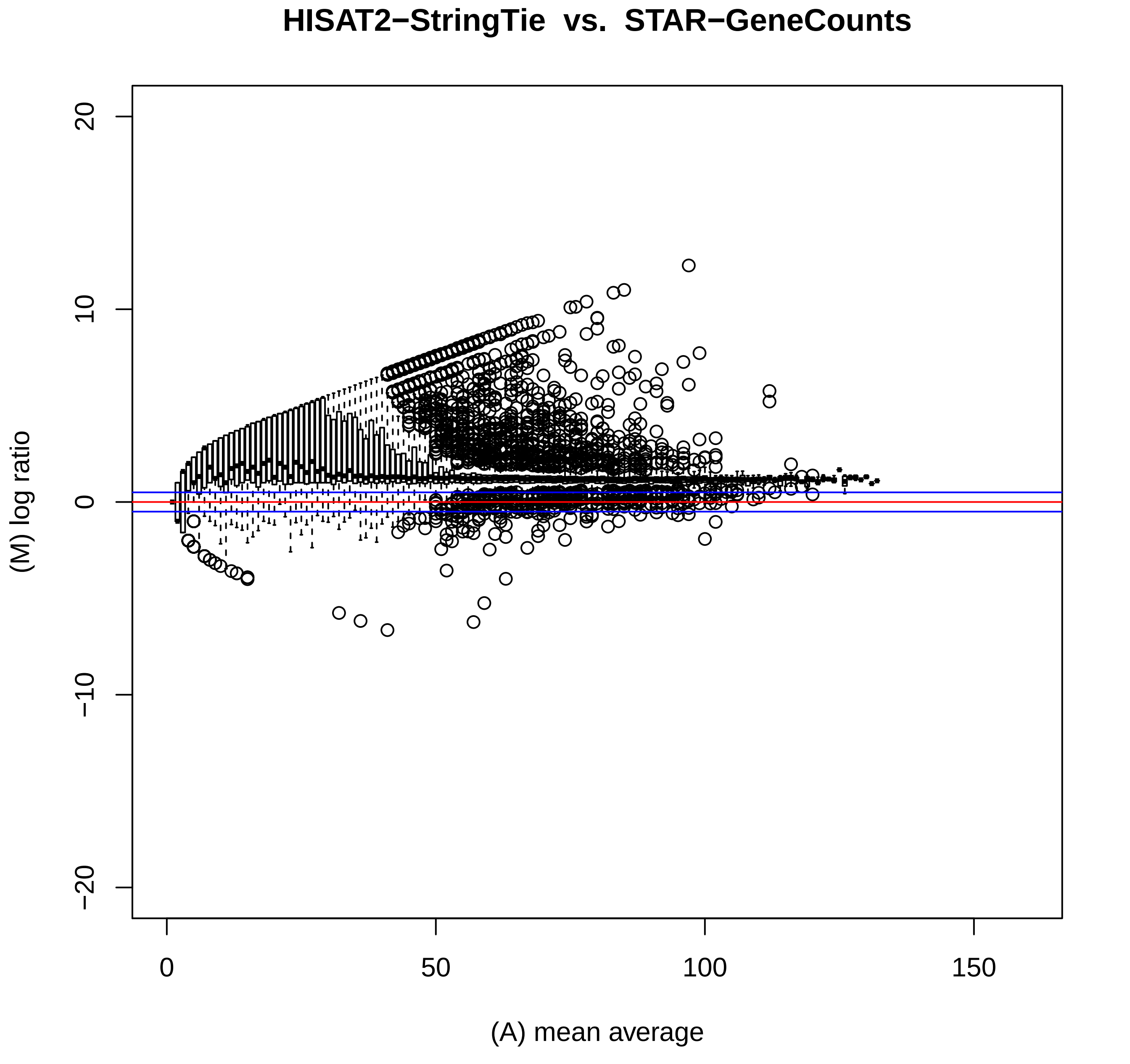

HISAT2−StringTie vs. STAR−GeneCounts
(M) log ratio
20
10
0
−10
−20
(A) mean average
0
50
100
150

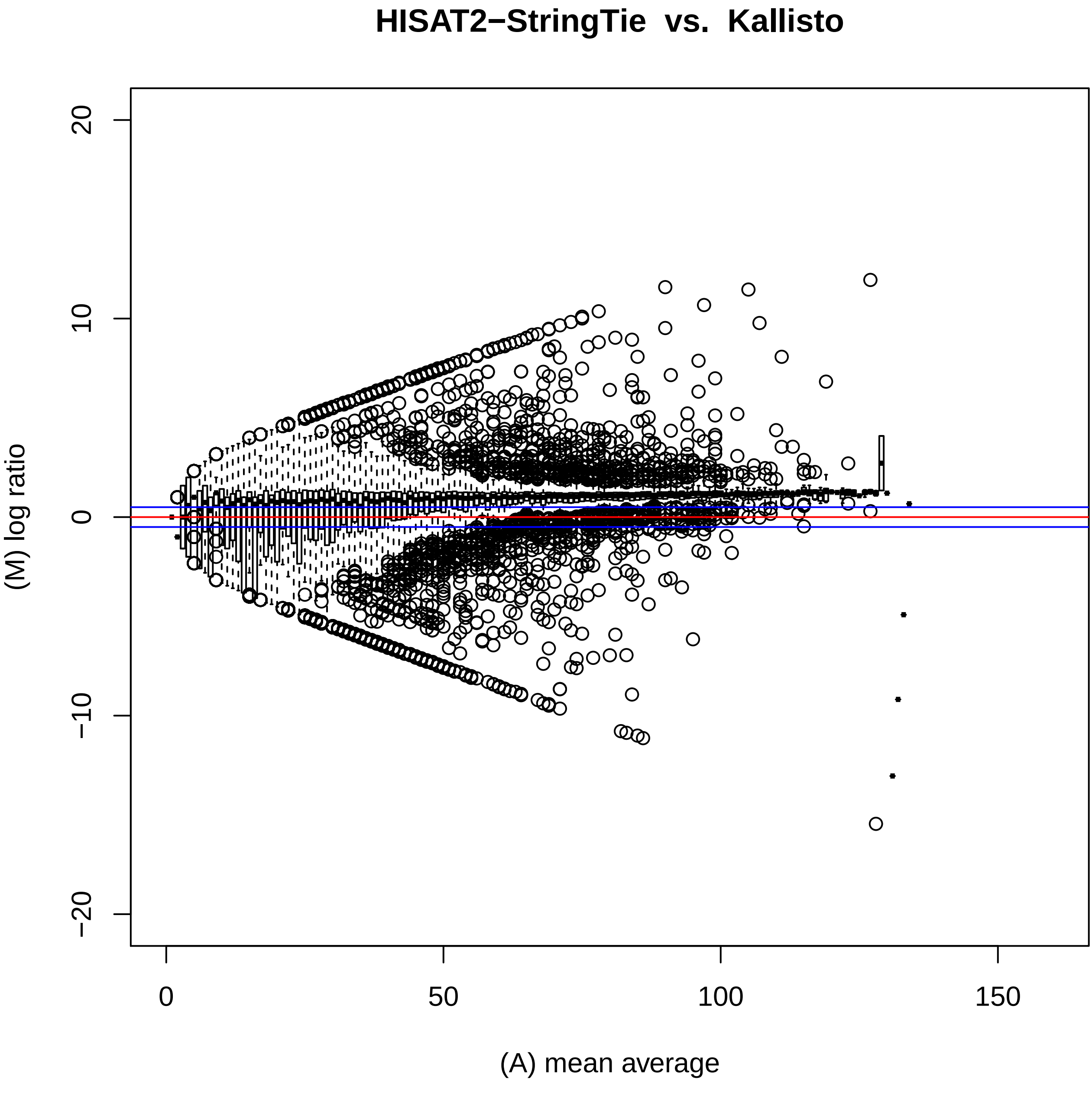
HISAT2−StringTie vs. Kallisto
(M) log ratio
(A) mean average
20
10
0
−10
−20
0
50
100
150

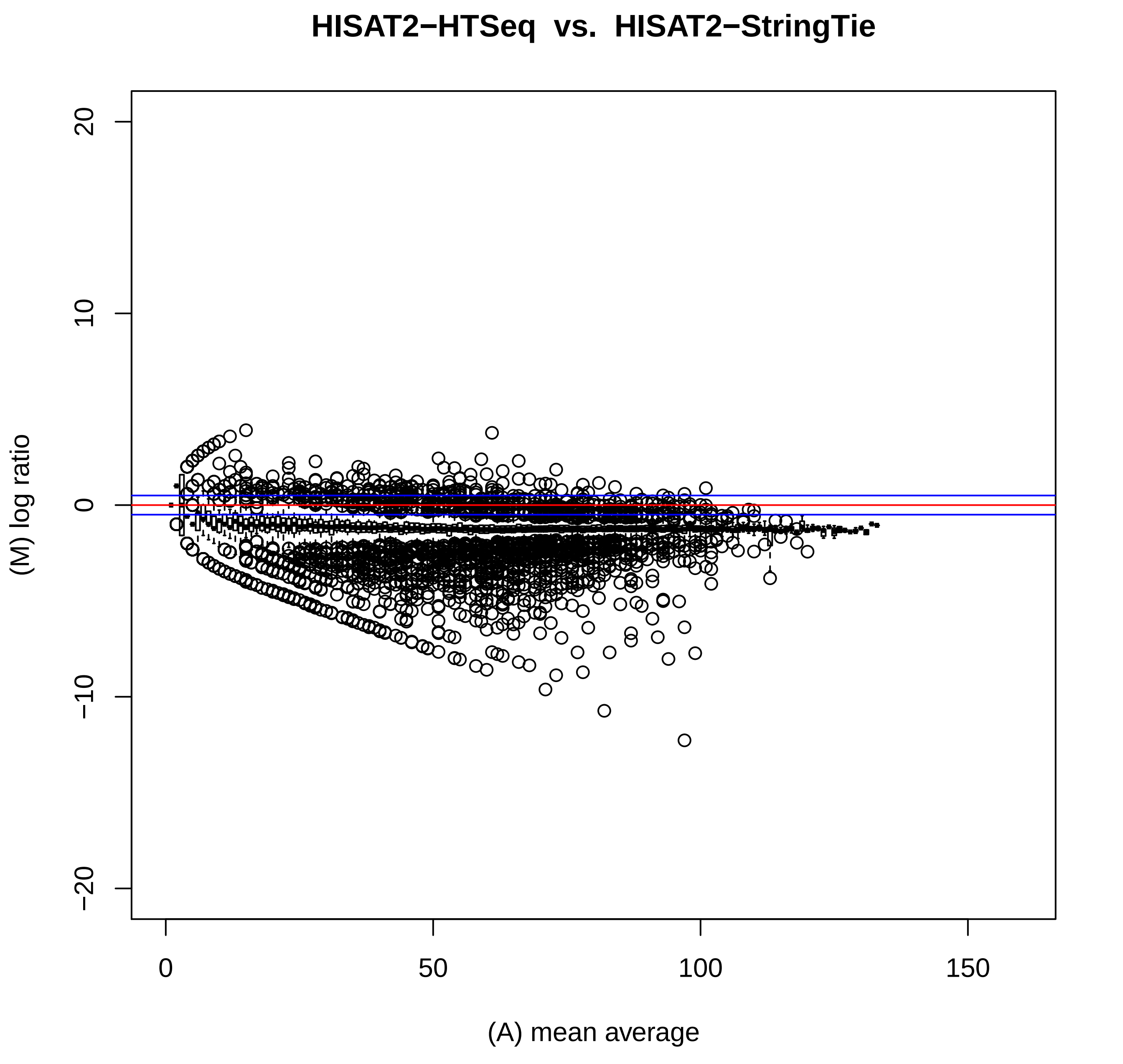
HISAT2−HTSeq vs. HISAT2−StringTie
(M) log ratio
20
10
0
−10
−20
(A) mean average
0
50
100
150

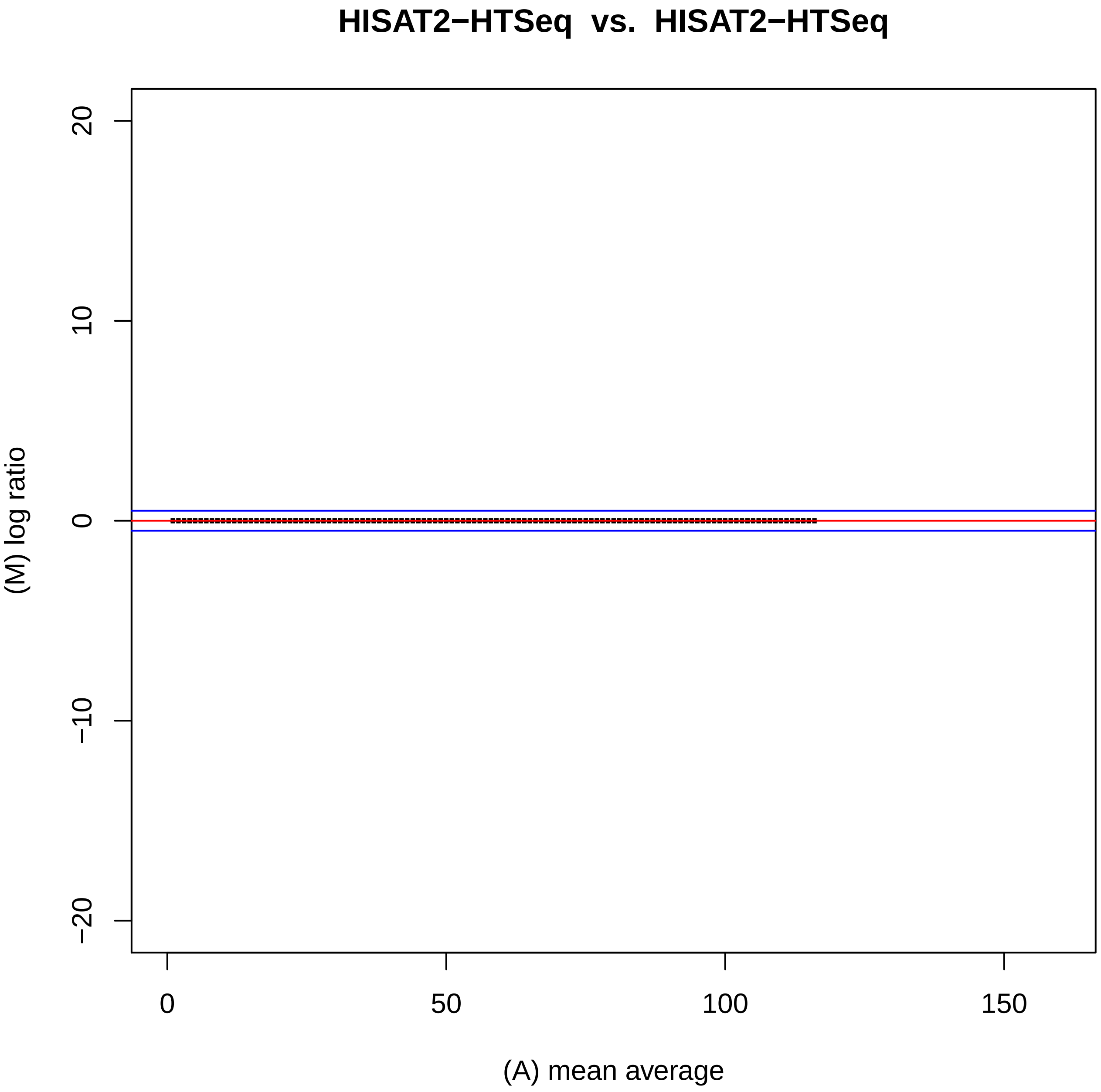

HISAT2−HTSeq vs. HISAT2−HTSeq
20
10
0
−10
−20
(M) log ratio
0
50
100
150
(A) mean average

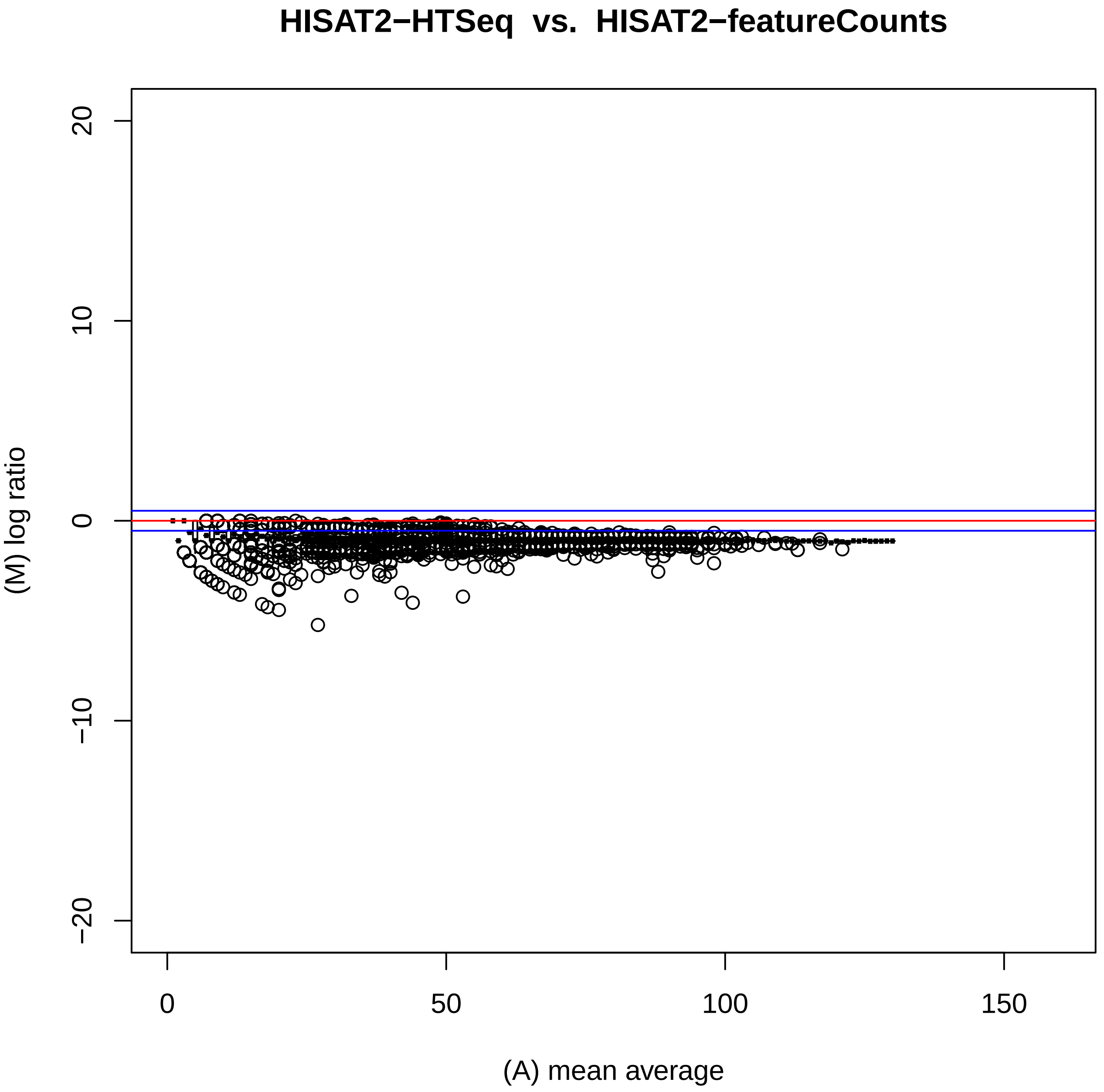
HISAT2−HTSeq vs. HISAT2−featureCounts
(M) log ratio
(A) mean average
20
10
0
−10
−20
0
50
100
150

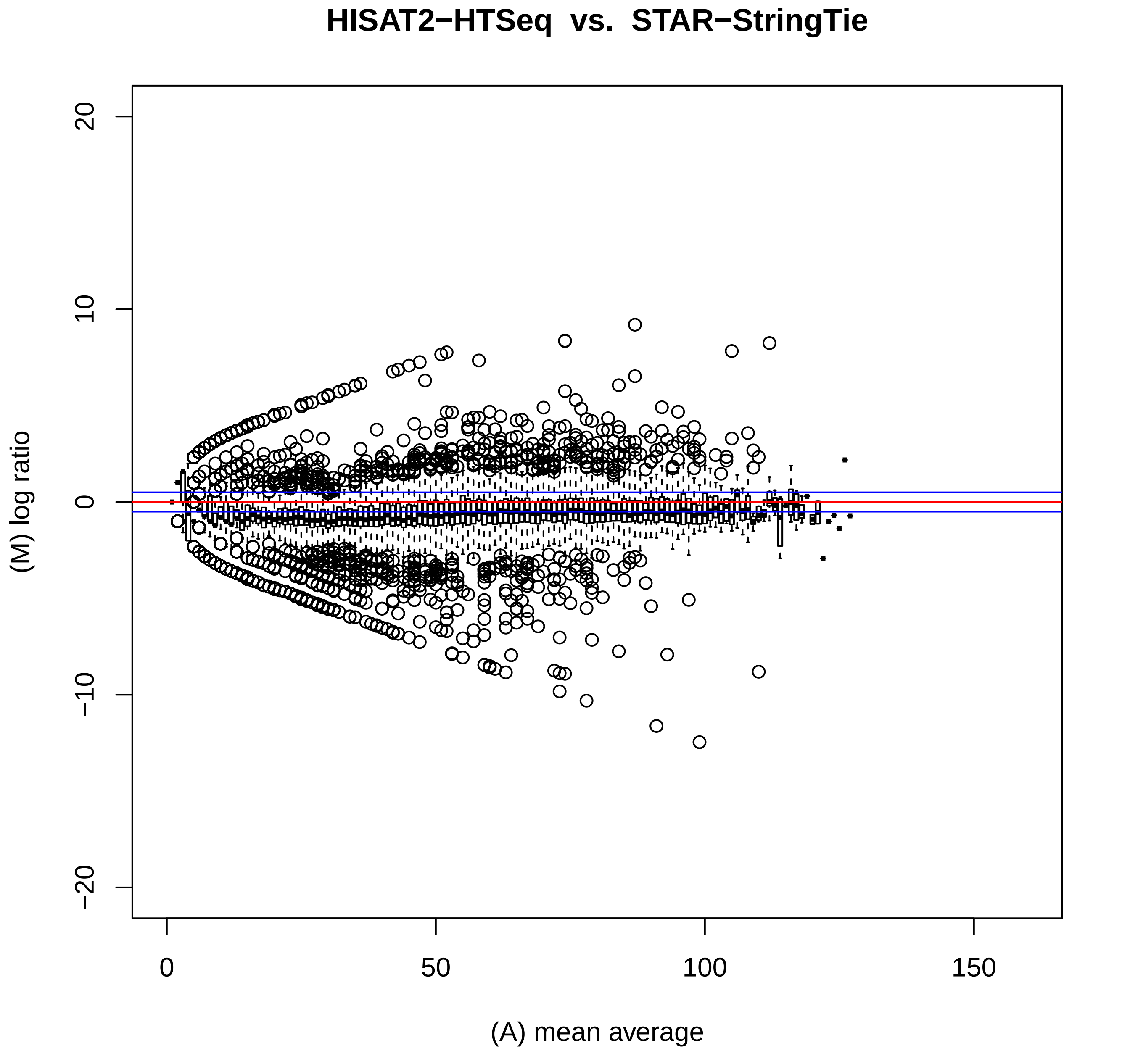

HISAT2−HTSeq vs. STAR−StringTie
(M) log ratio
(A) mean average
20
10
0
−10
−20
0
50
100
150

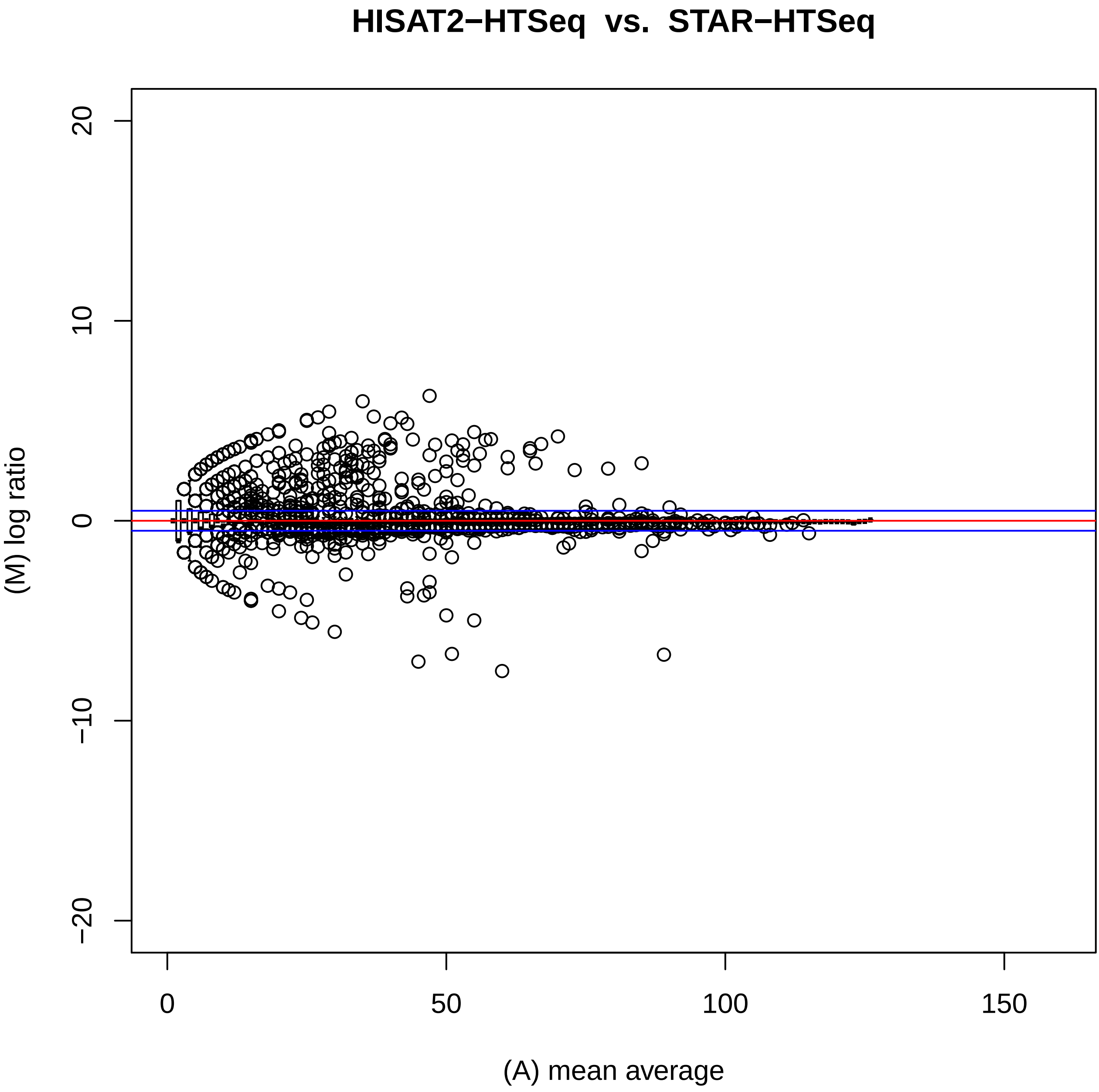
HISAT2−HTSeq vs. STAR−HTSeq
(M) log ratio
(A) mean average
20
10
0
−10
−20
0
50
100
150

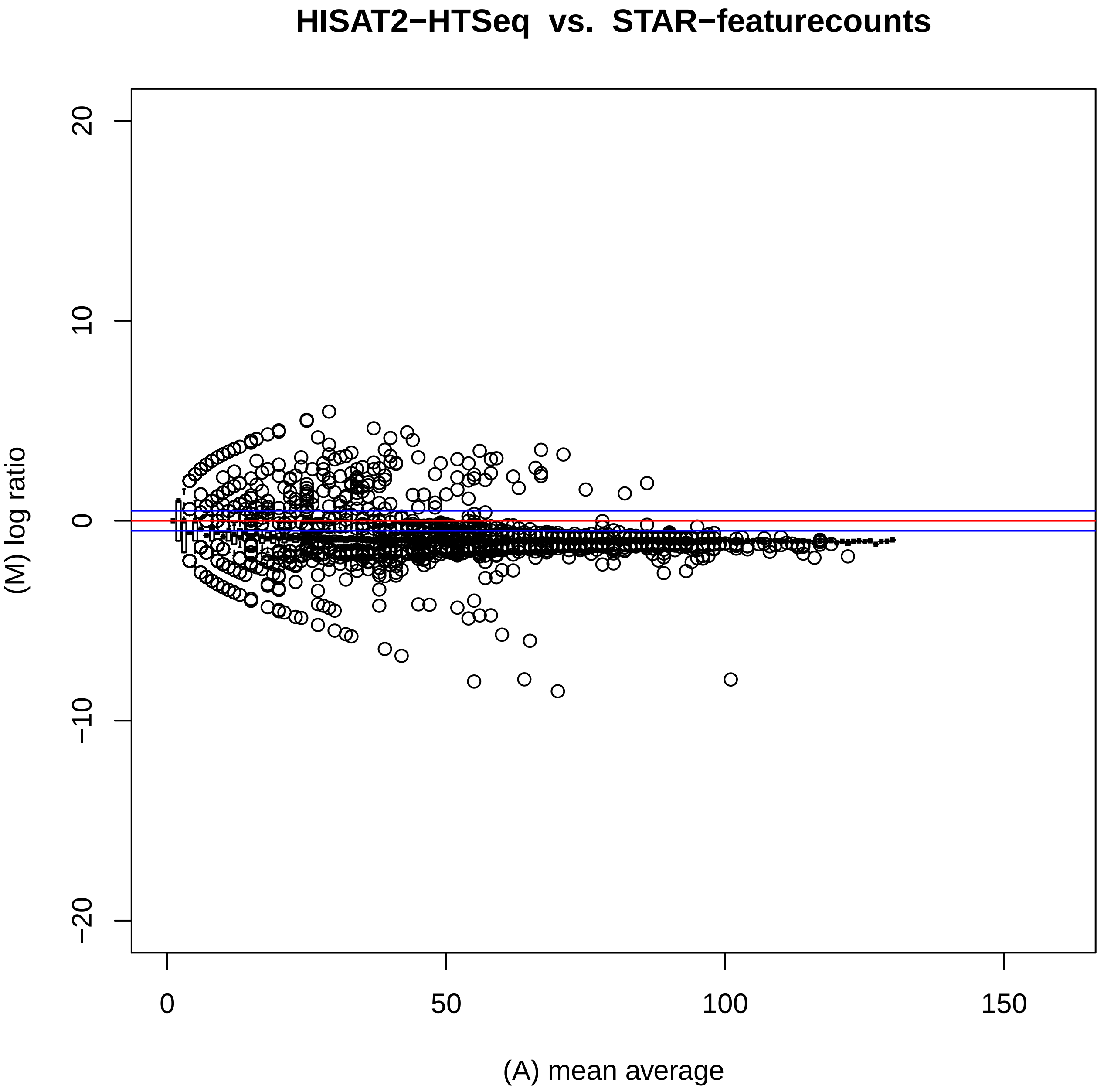
HISAT2−HTSeq vs. STAR−featurecounts
(M) log ratio
(A) mean average
20
10
0
−10
−20
0
50
100
150

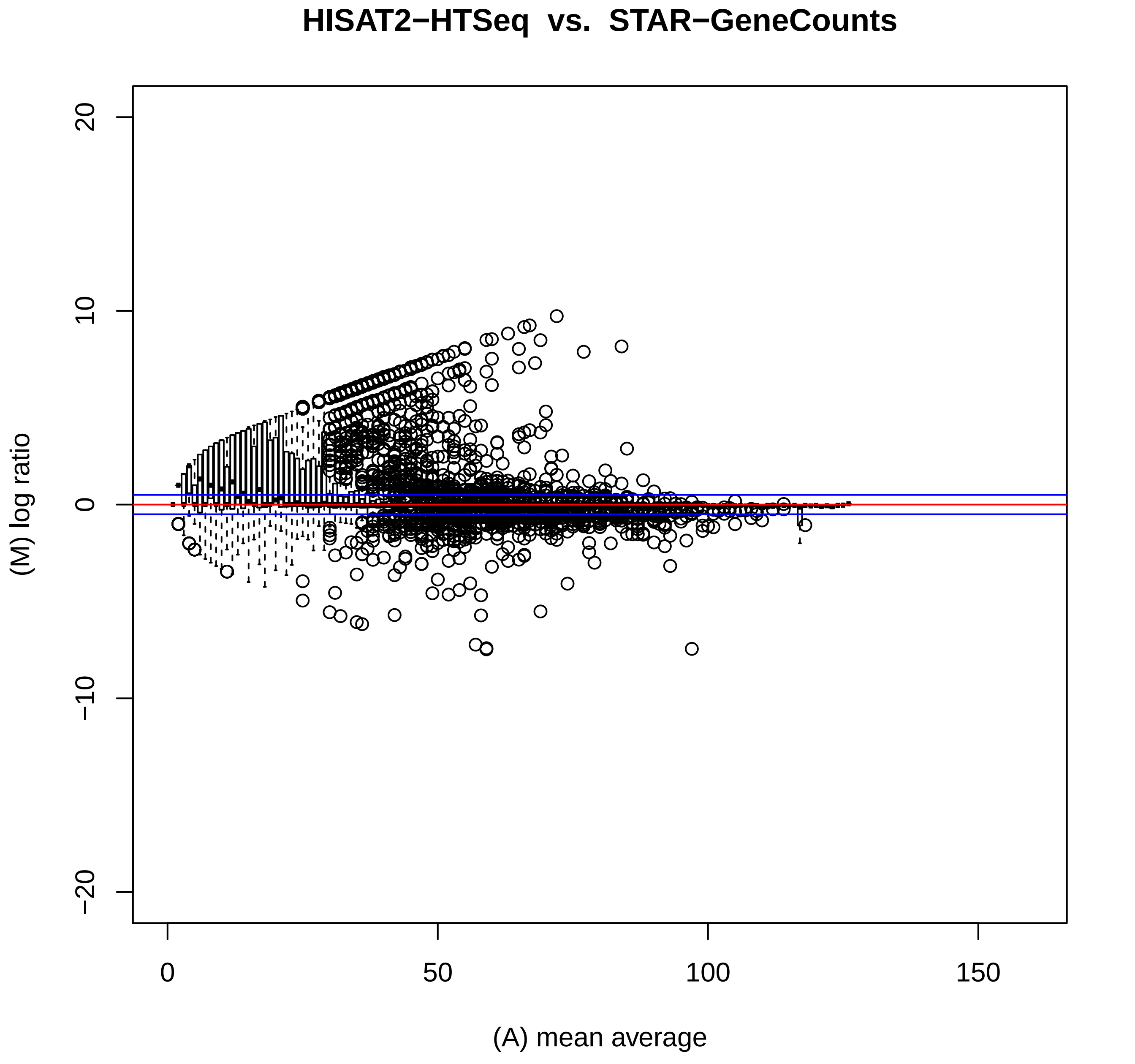

HISAT2−HTSeq vs. STAR−GeneCounts
(M) log ratio
20
10
0
−10
−20
0
50
100
150
(A) mean average

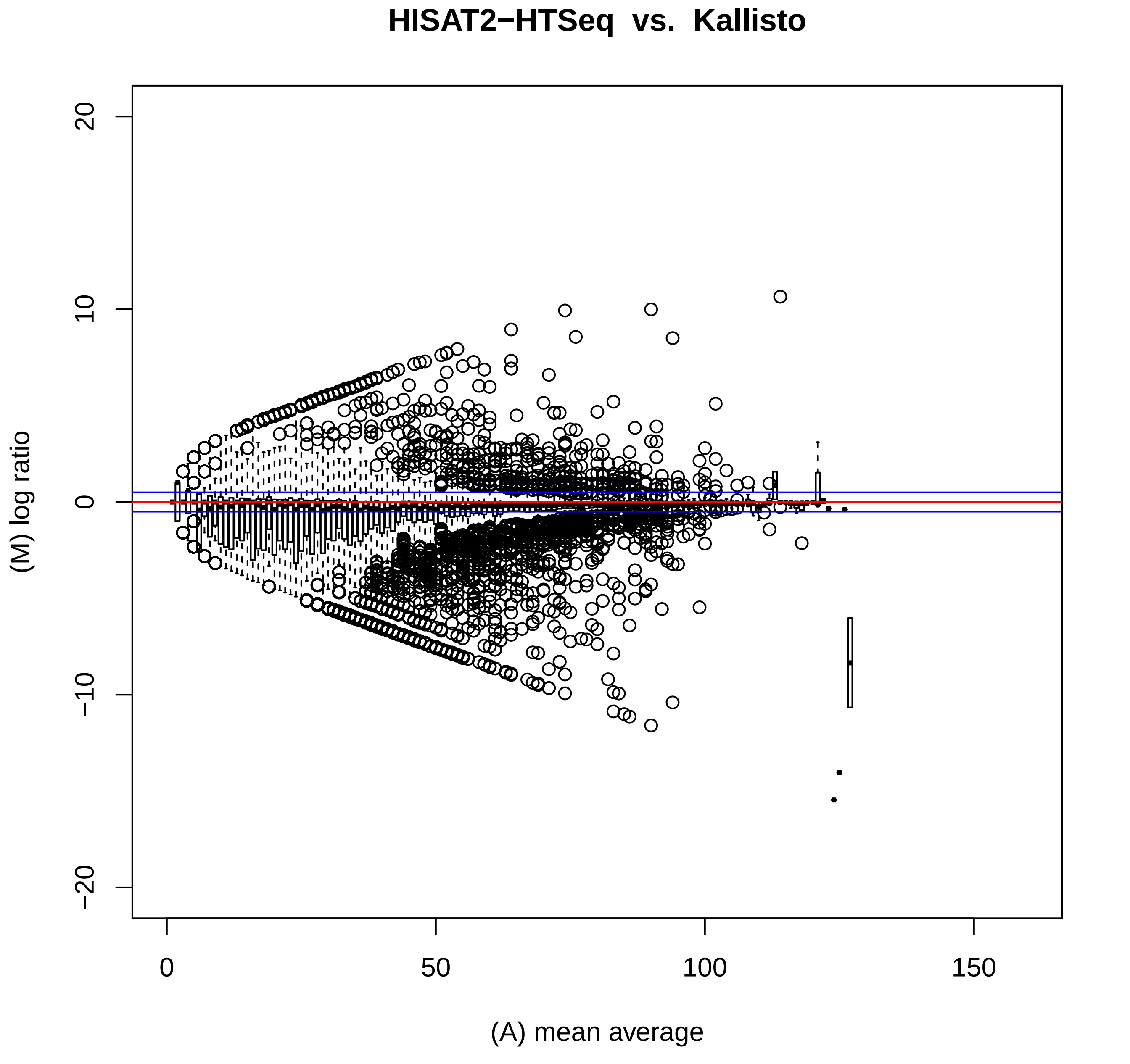
HISAT2−HTSeq vs. Kallisto
(M) log ratio
20
10
0
−10
−20
0
50
100
150
(A) mean average

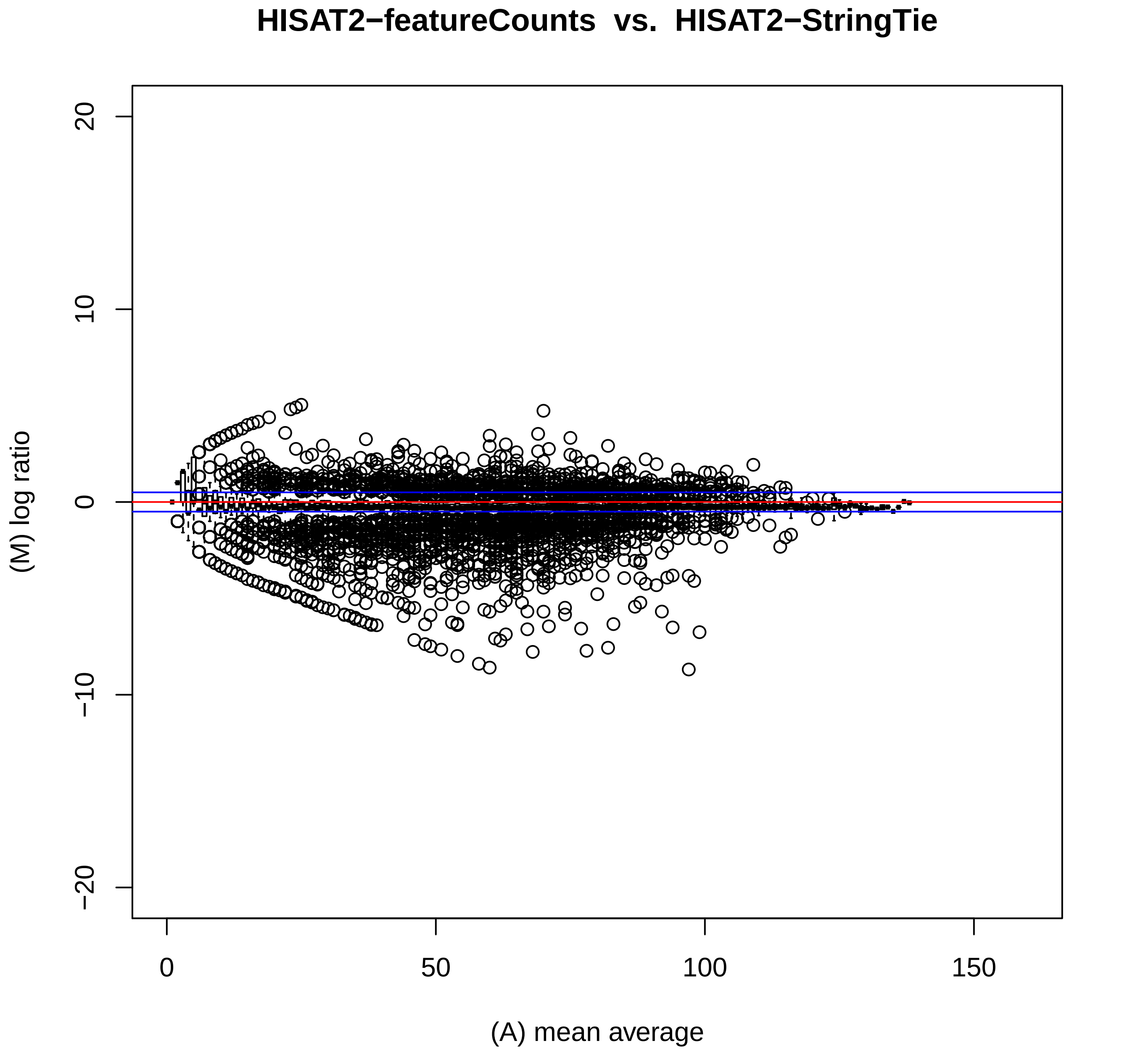

HISAT2−featureCounts vs. HISAT2−StringTie
(M) log ratio
(A) mean average
20
10
0
−10
−20
0
50
100
150

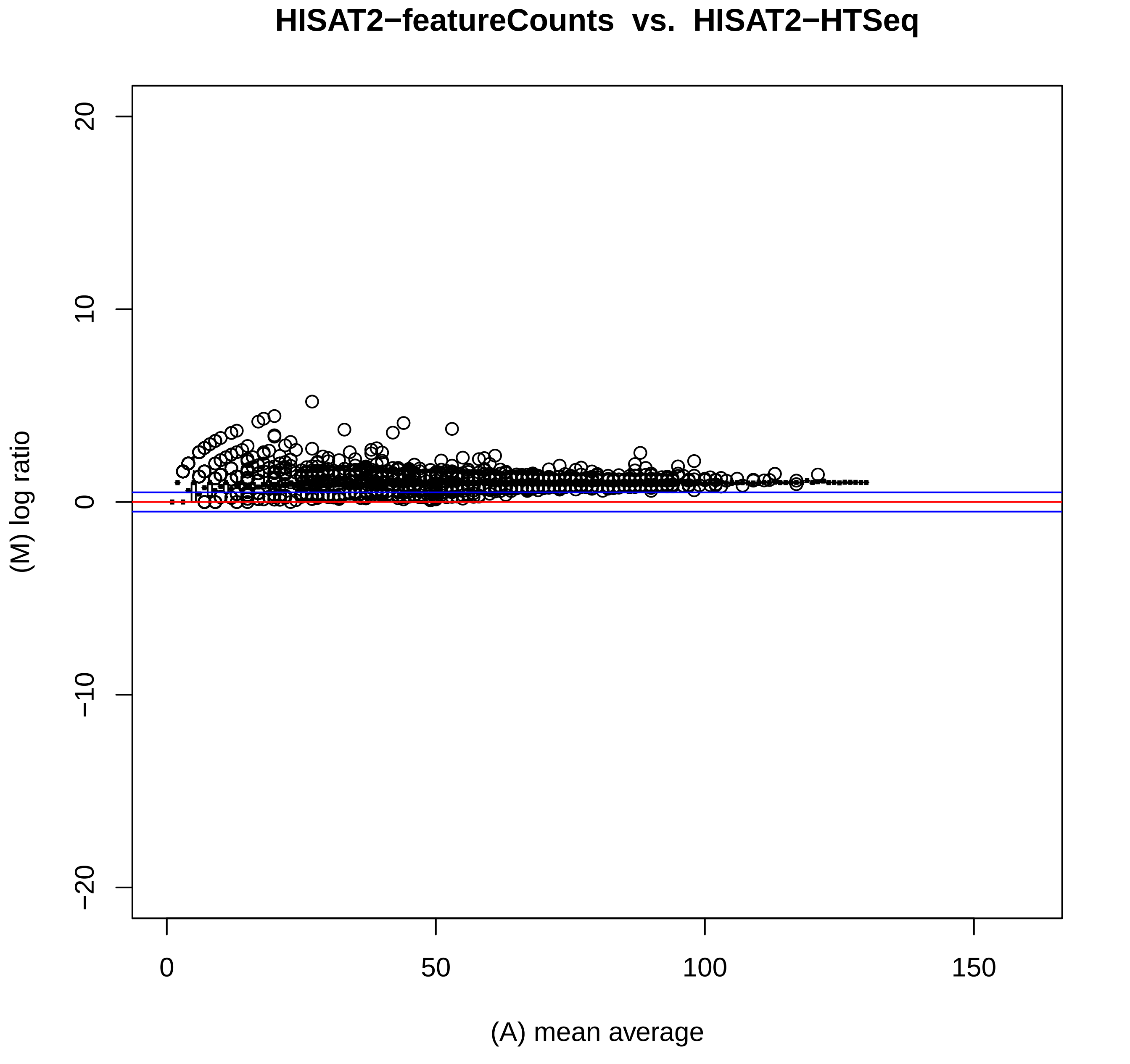
HISAT2−featureCounts vs. HISAT2−HTSeq
(M) log ratio
(A) mean average
20
10
0
−10
−20
0
50
100
150

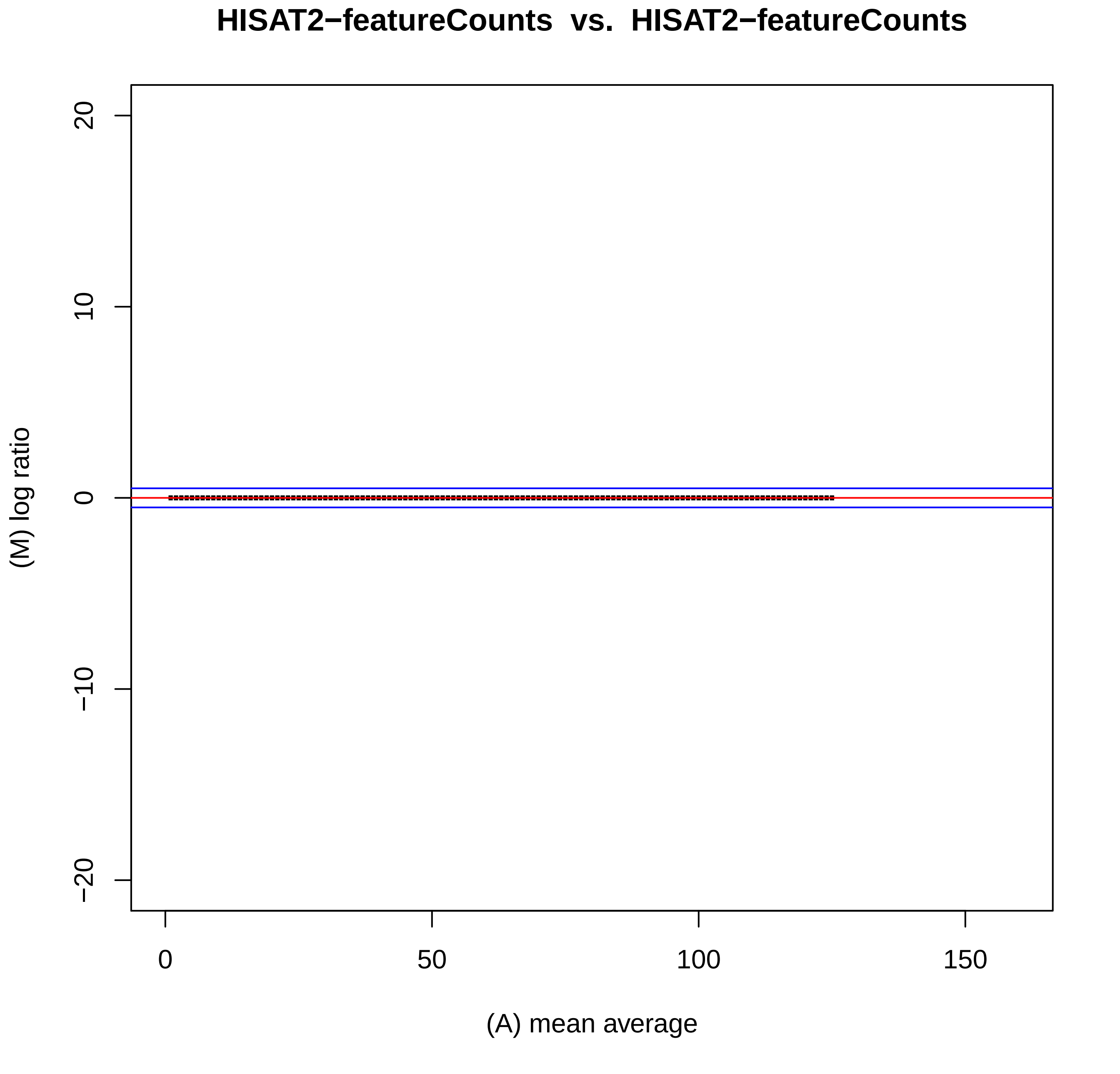

HISAT2−featureCounts vs. HISAT2−featureCounts
20
10
0
−10
−20
(M) log ratio
0
50
100
150
(A) mean average

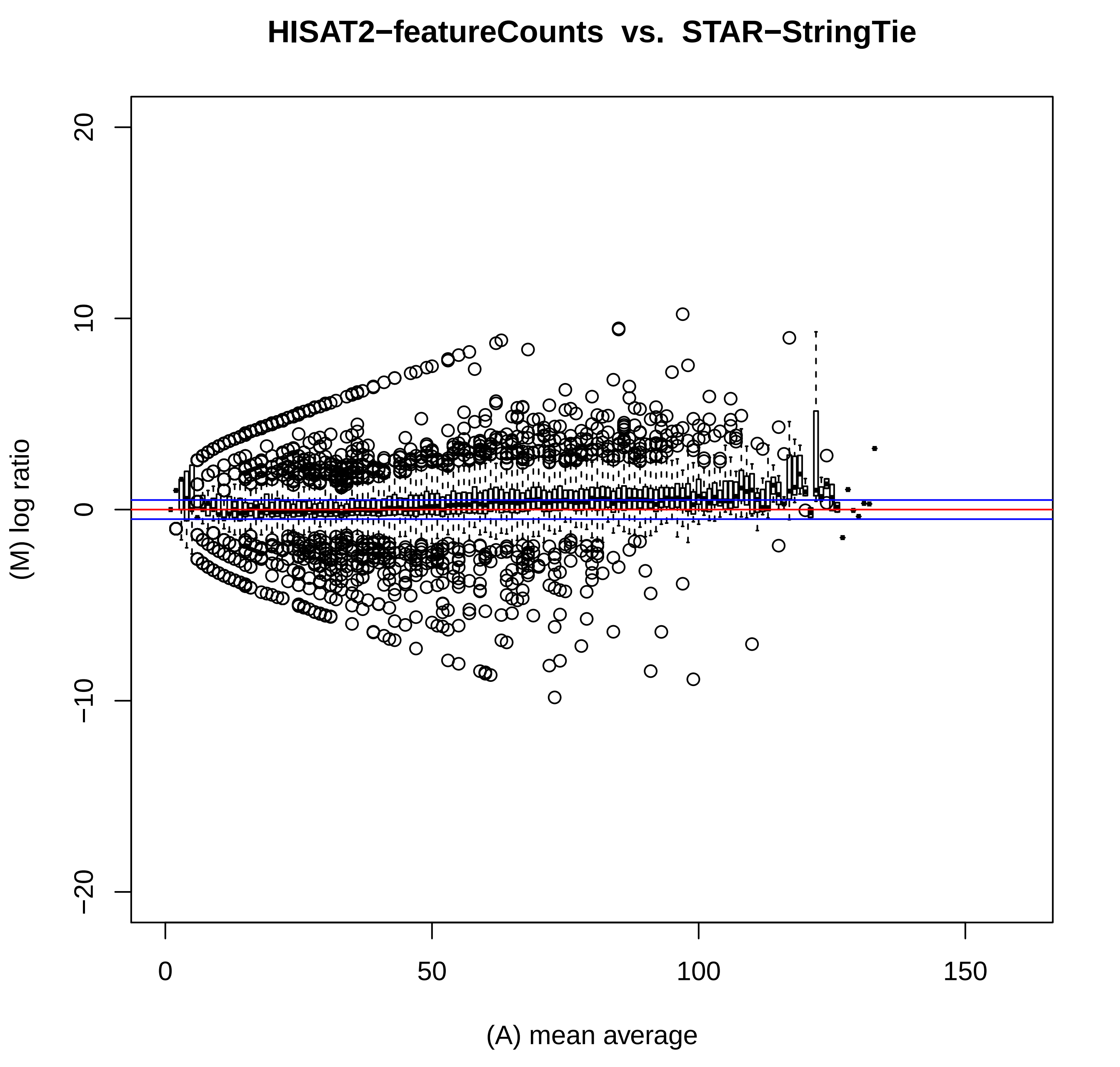
HISAT2−featureCounts vs. STAR−StringTie
(M) log ratio
(A) mean average
20
10
0
−10
−20
0
50
100
150

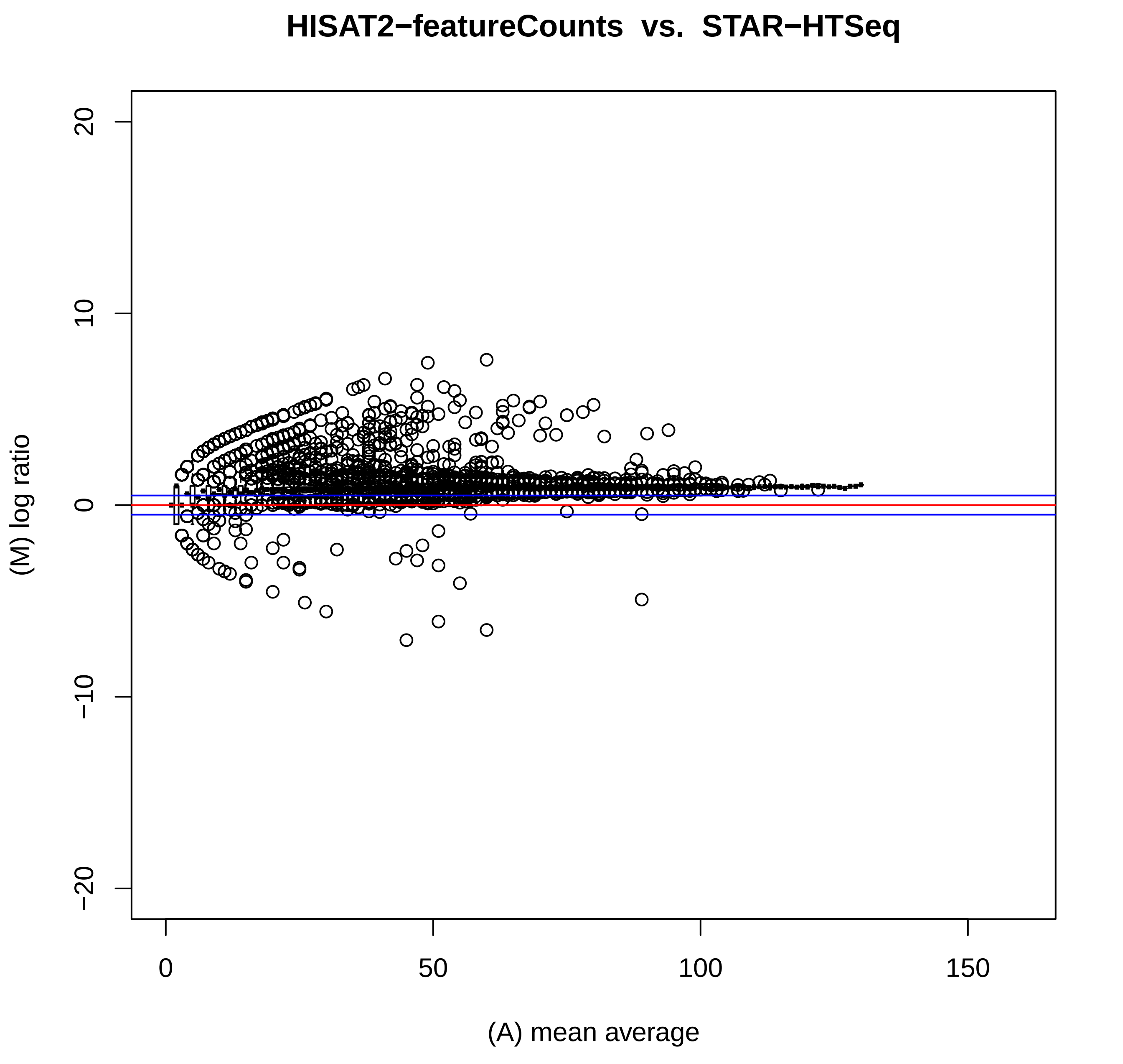
HISAT2−featureCounts vs. STAR−HTSeq
(M) log ratio
(A) mean average
20
10
0
−10
−20
0
50
100
150

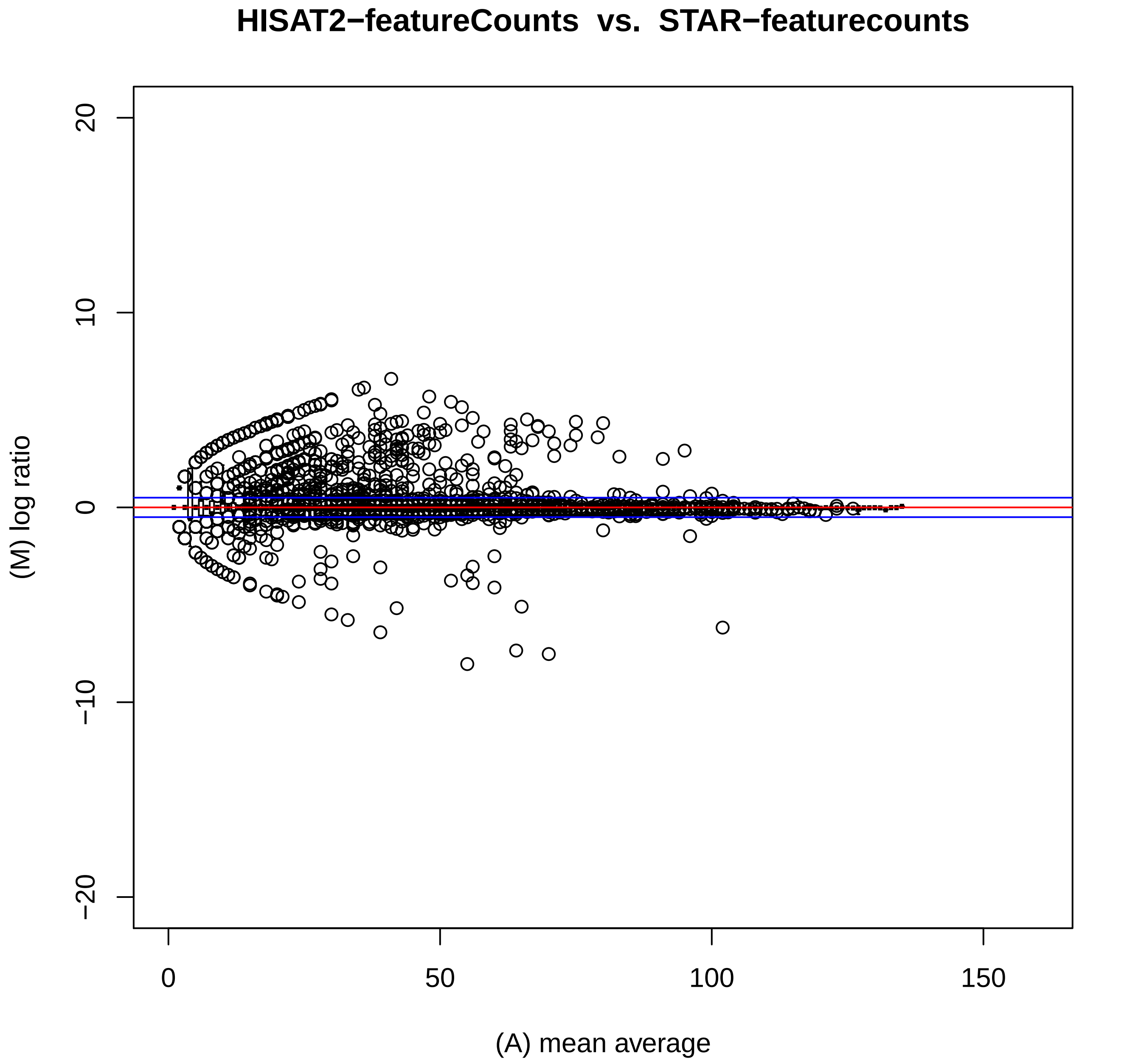
HISAT2−featureCounts vs. STAR−featurecounts
(M) log ratio
(A) mean average
20
10
0
−10
−20
0
50
100
150

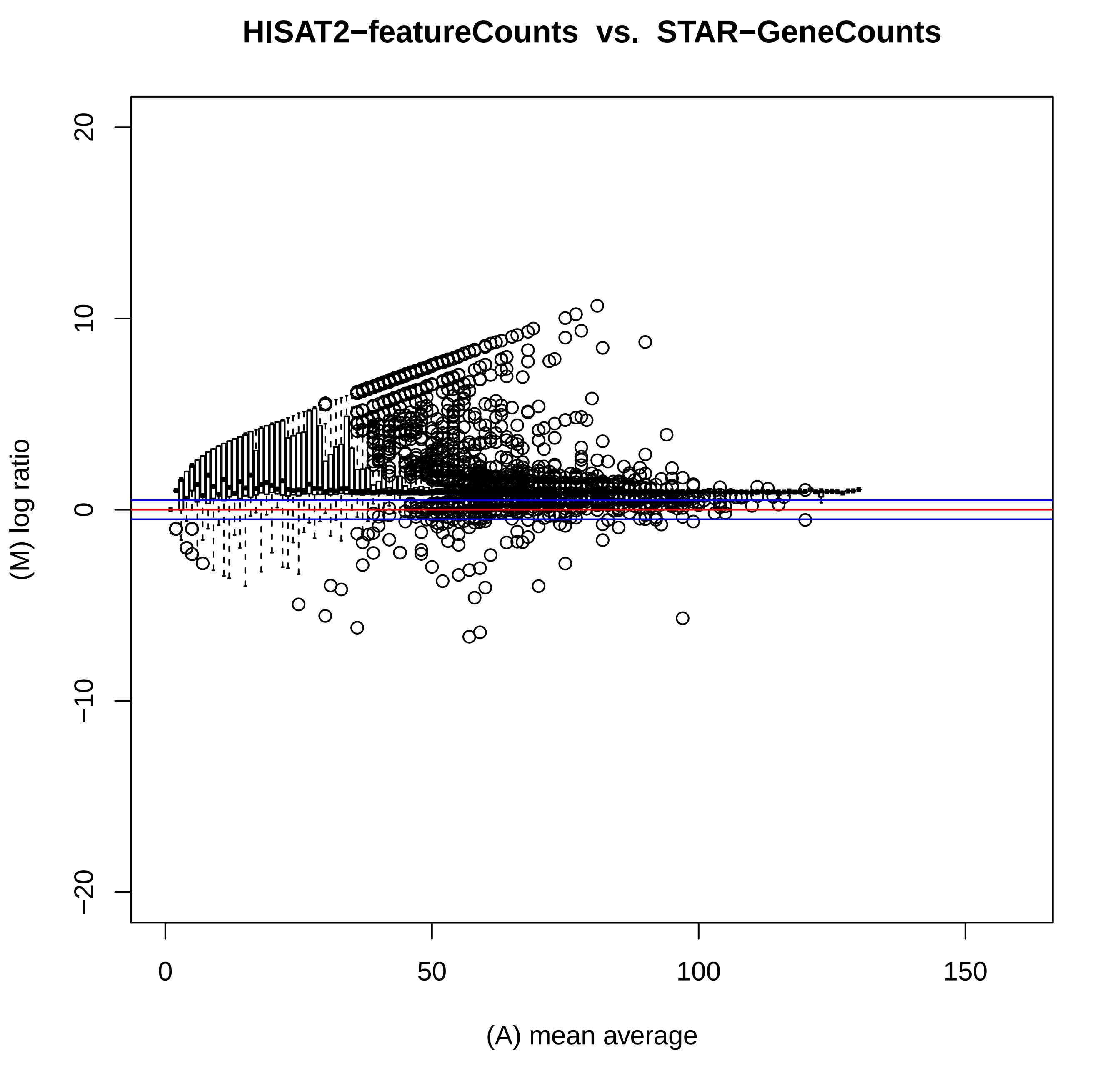

HISAT2−featureCounts vs. STAR−GeneCounts
(M) log ratio
(A) mean average
20
10
0
−10
−20
0
50
100
150

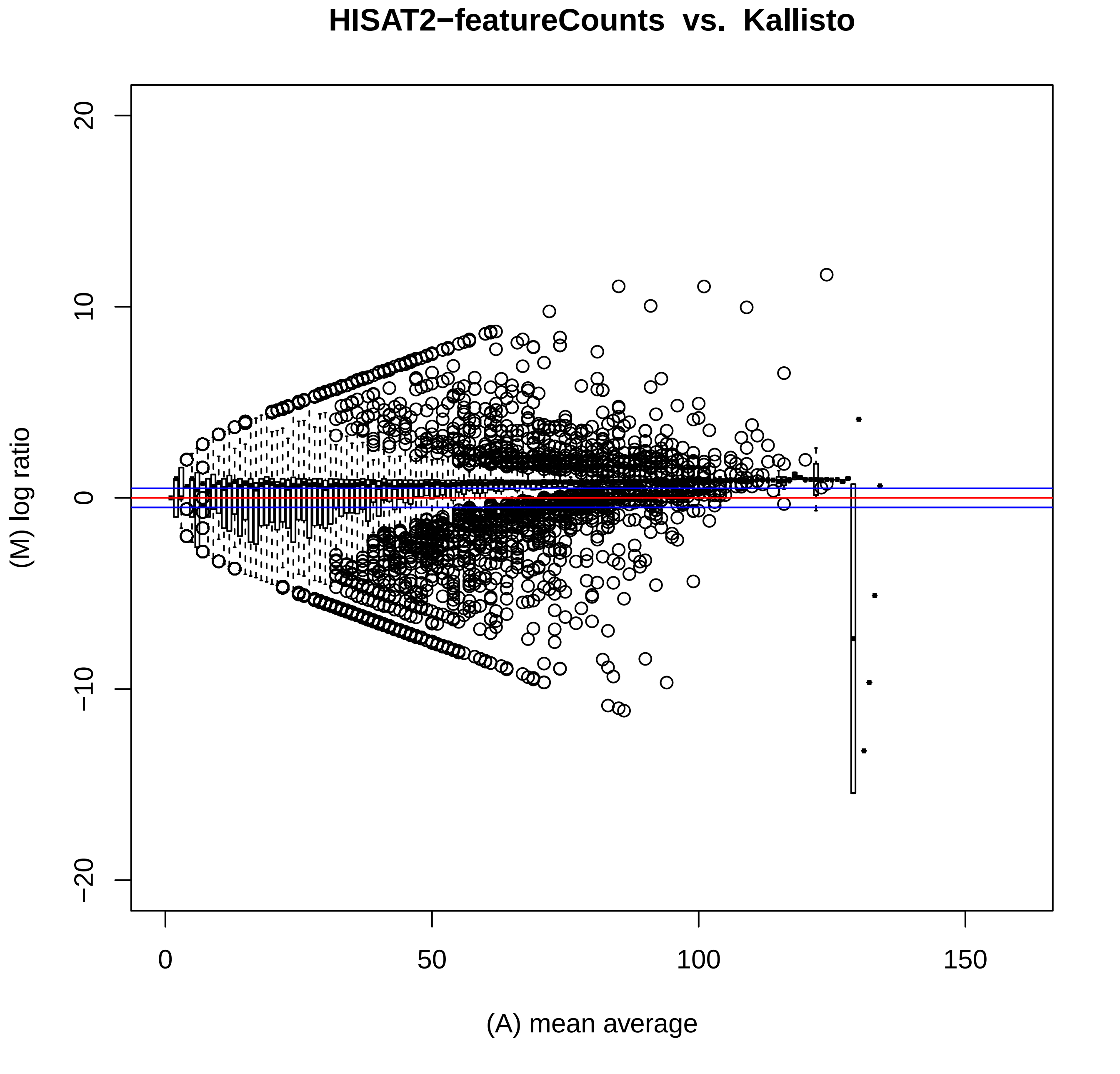
HISAT2−featureCounts vs. Kallisto
(M) log ratio
(A) mean average
20
10
0
−10
−20
0
50
100
150

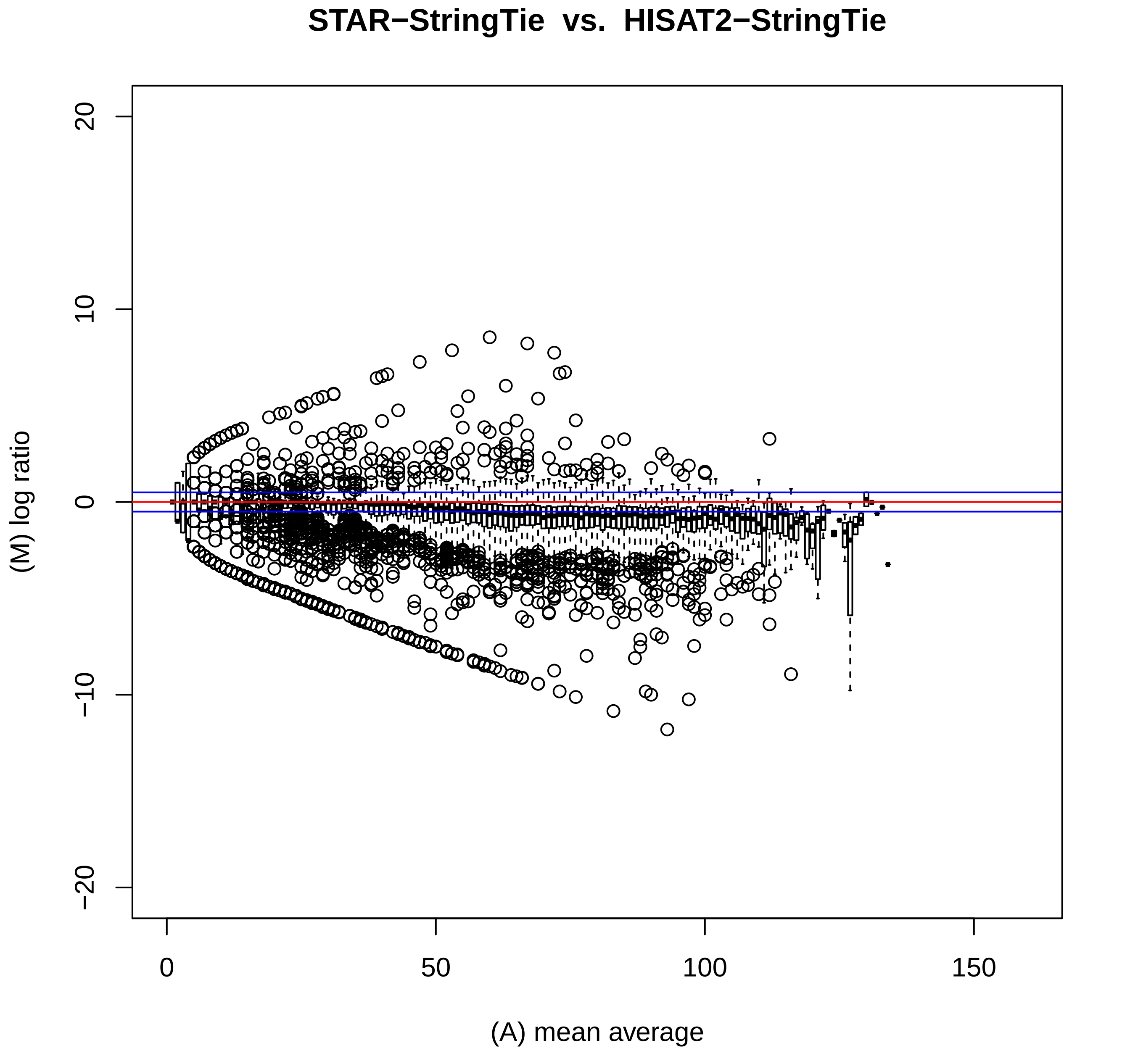

STAR−StringTie vs. HISAT2−StringTie
20
10
0
−10
−20
(M) log ratio
0
50
100
150
(A) mean average

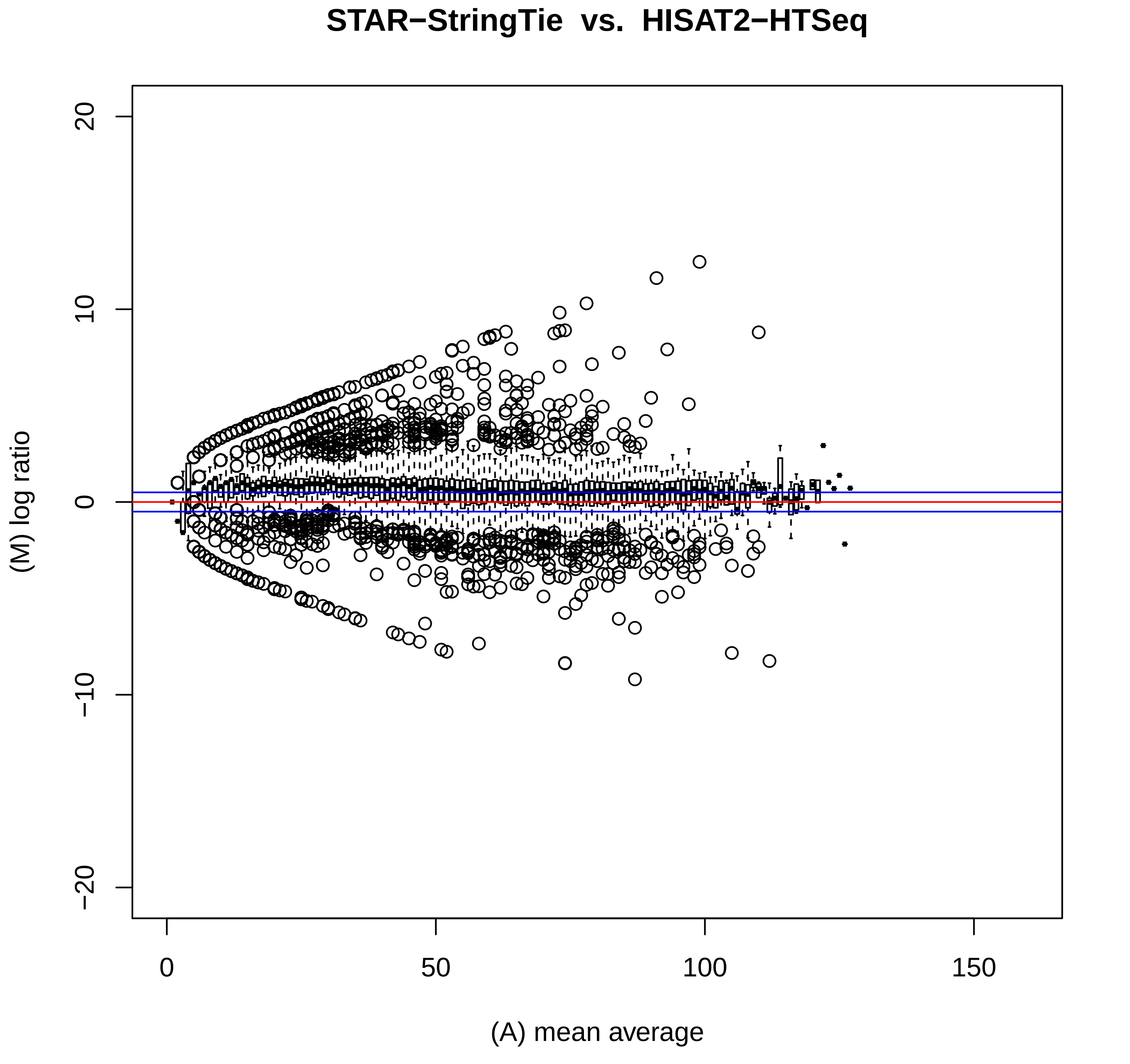
STAR−StringTie vs. HISAT2−HTSeq
(M) log ratio
(A) mean average
20
10
0
−10
−20
0
50
100
150

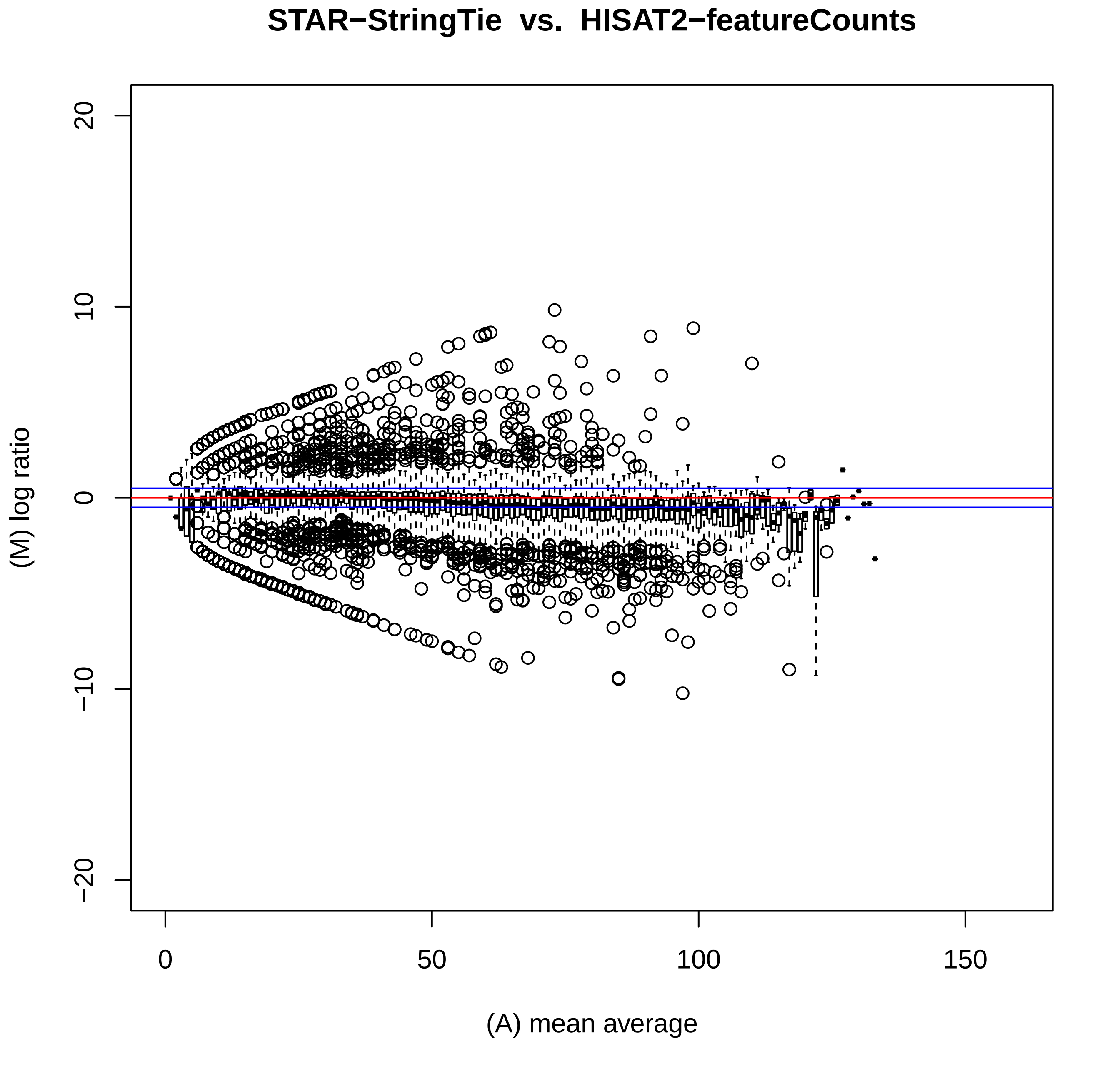
STAR−StringTie vs. HISAT2−featureCounts
(M) log ratio
(A) mean average
20
10
0
−10
−20
0
50
100
150

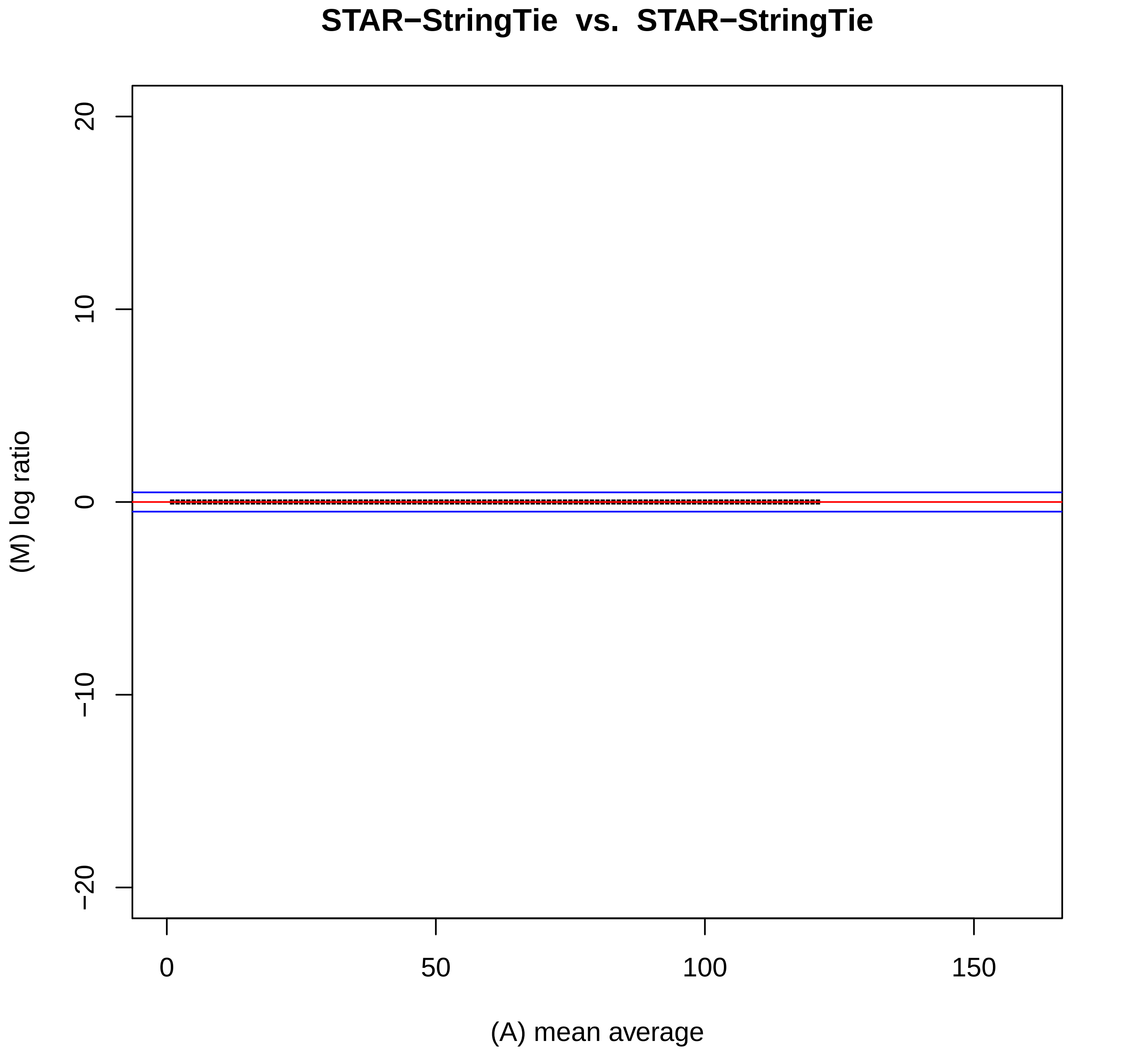
STAR−StringTie vs. STAR−StringTie
(M) log ratio
(A) mean average
20
10
0
−10
−20
0
50
100
150

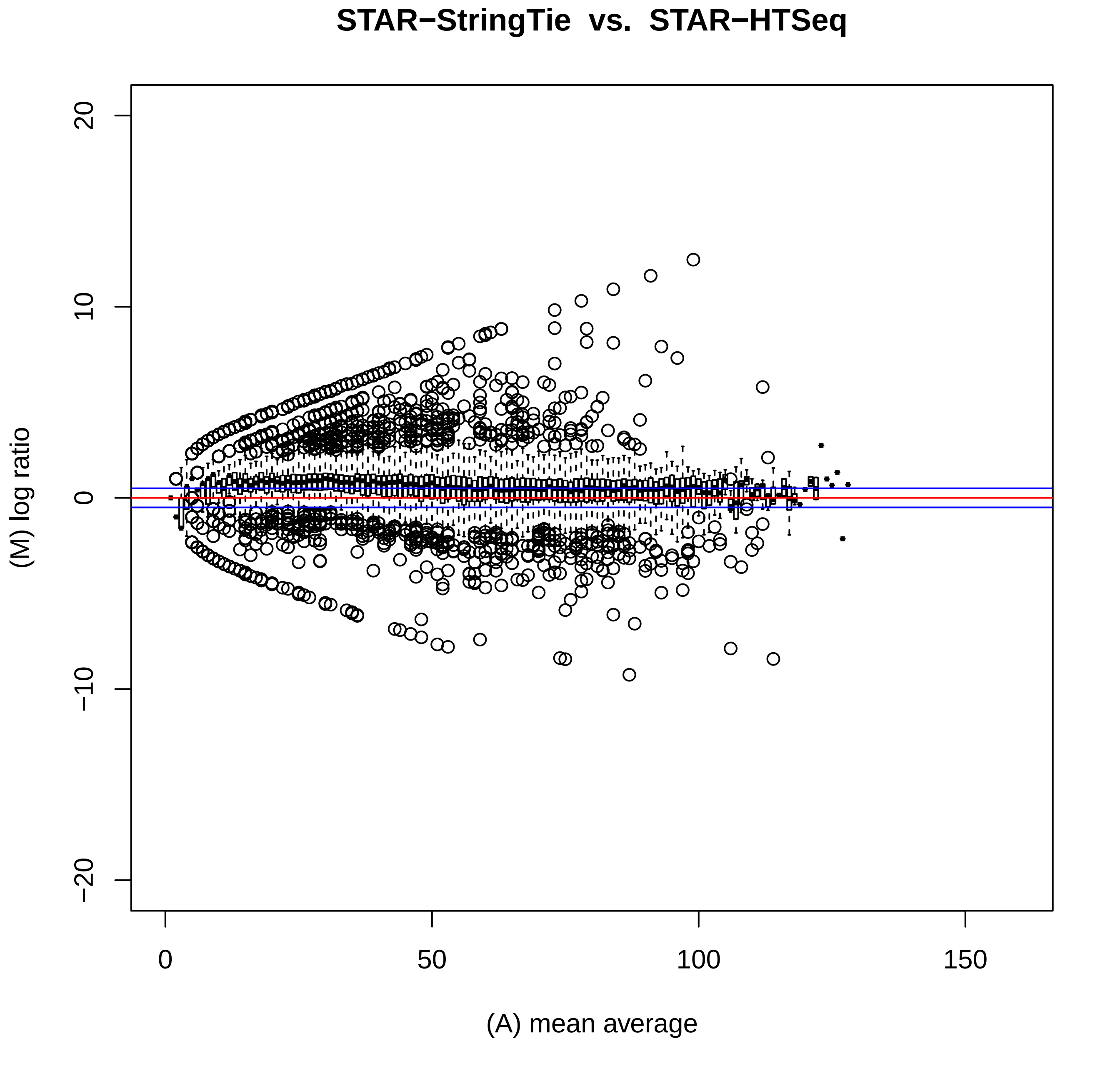
STAR−StringTie vs. STAR−HTSeq
(M) log ratio
(A) mean average
20
10
0
−10
−20
0
50
100
150

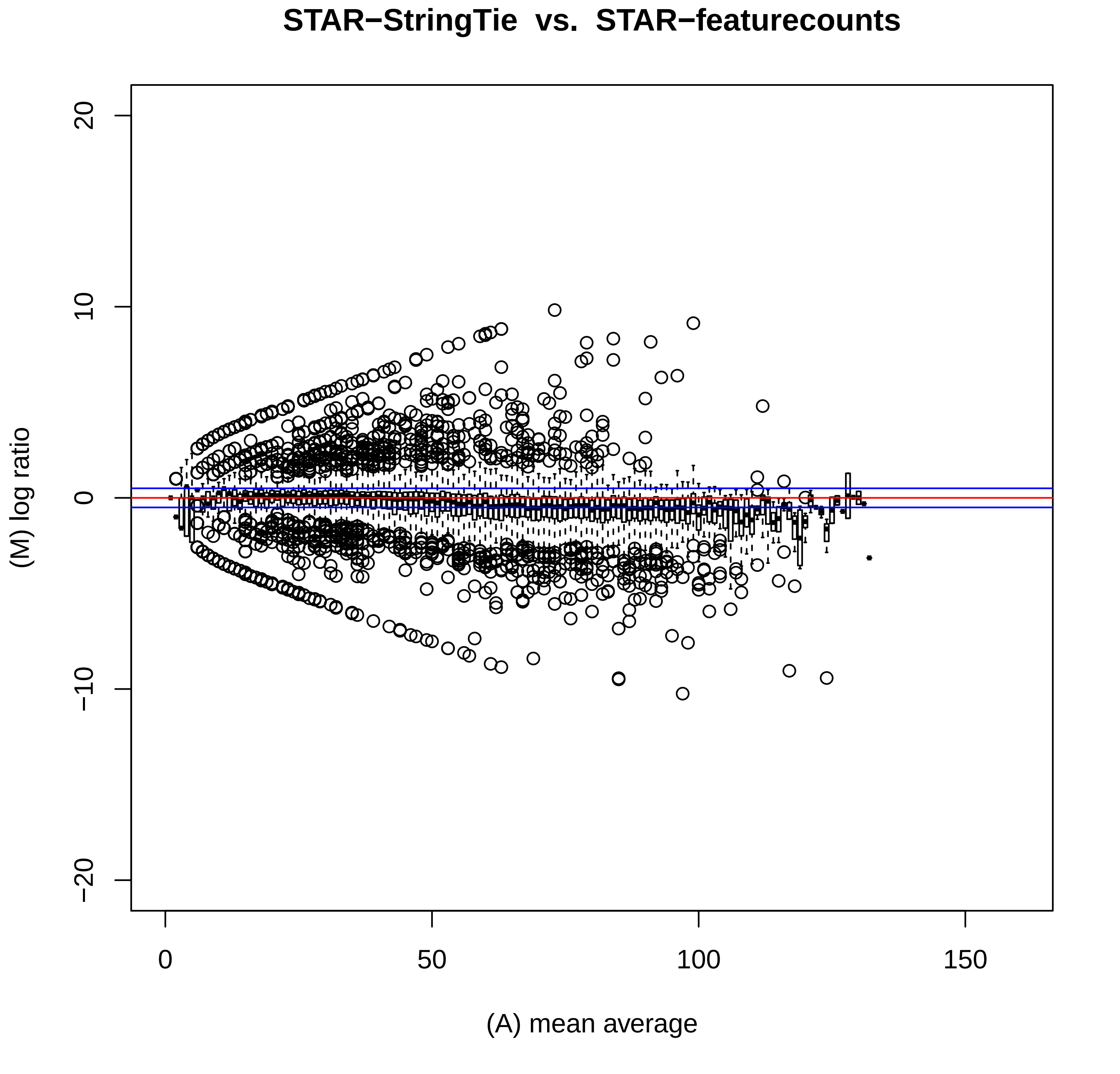

STAR−StringTie vs. STAR−featurecounts
20
10
0
−10
−20
(M) log ratio
0
50
100
150
(A) mean average

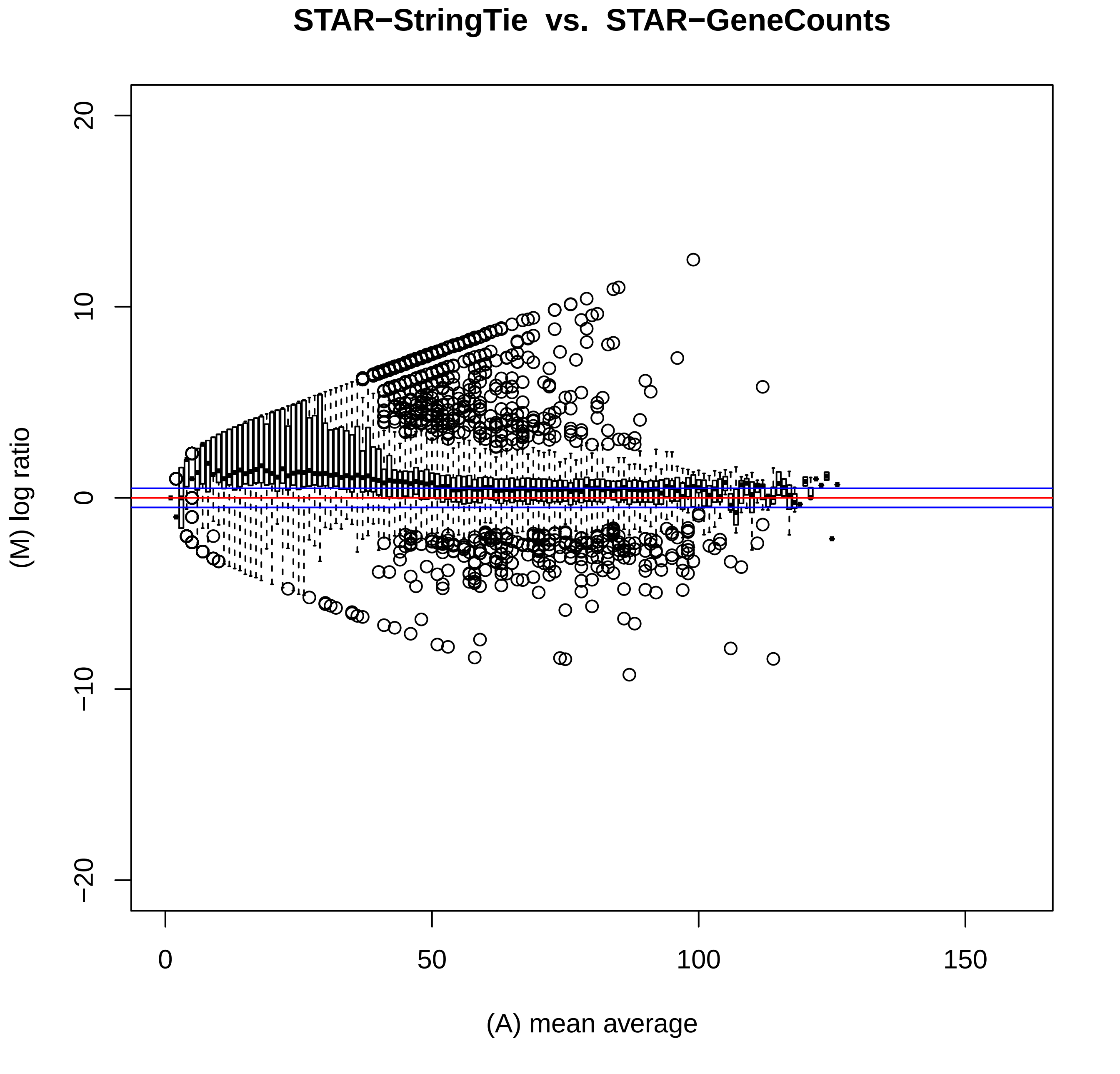
STAR−StringTie vs. STAR−GeneCounts
(M) log ratio
(A) mean average
20
10
0
−10
−20
0
50
100
150

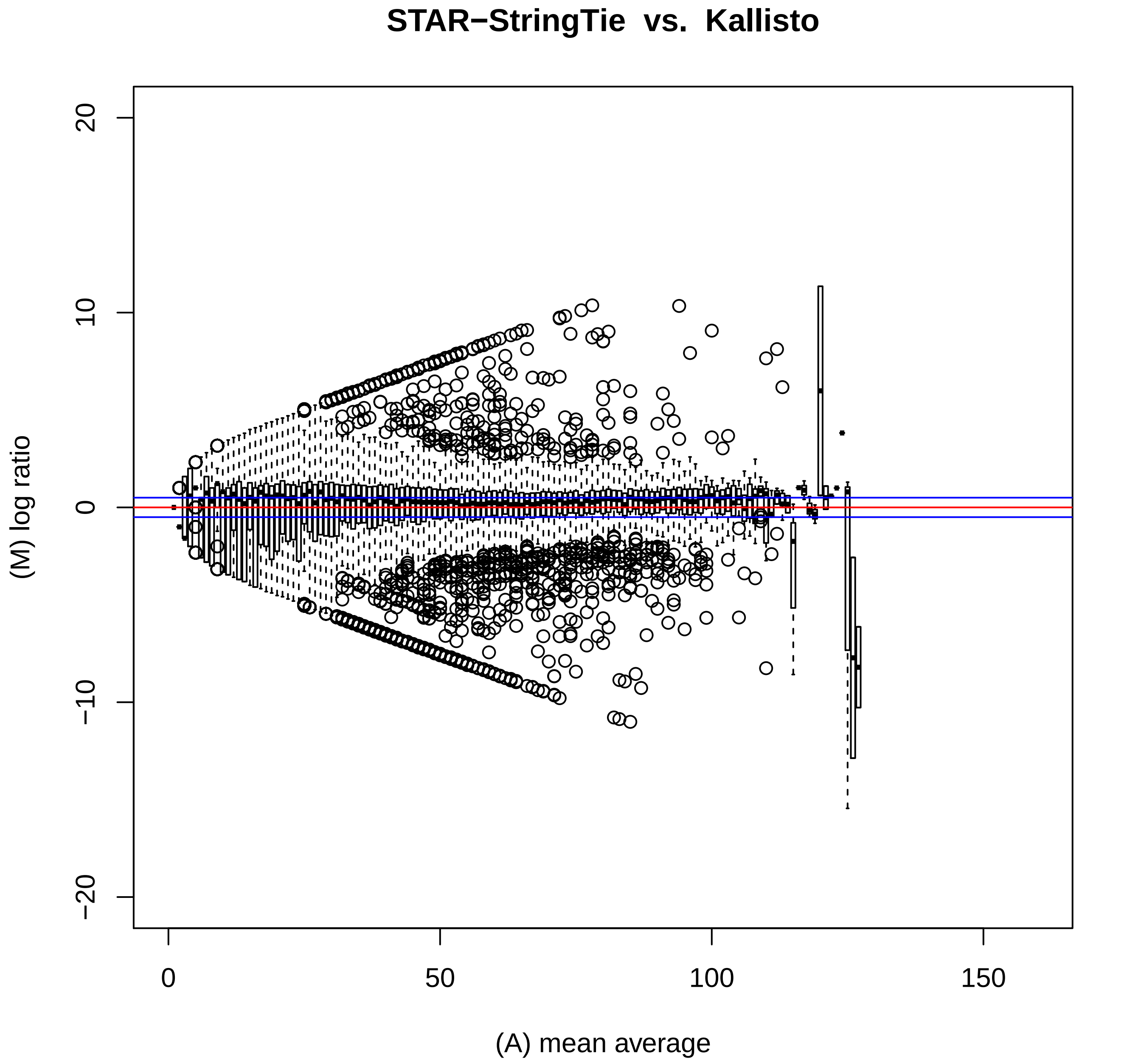

STAR−StringTie vs. Kallisto
(M) log ratio
(A) mean average
20
10
0
−10
−20
0
50
100
150

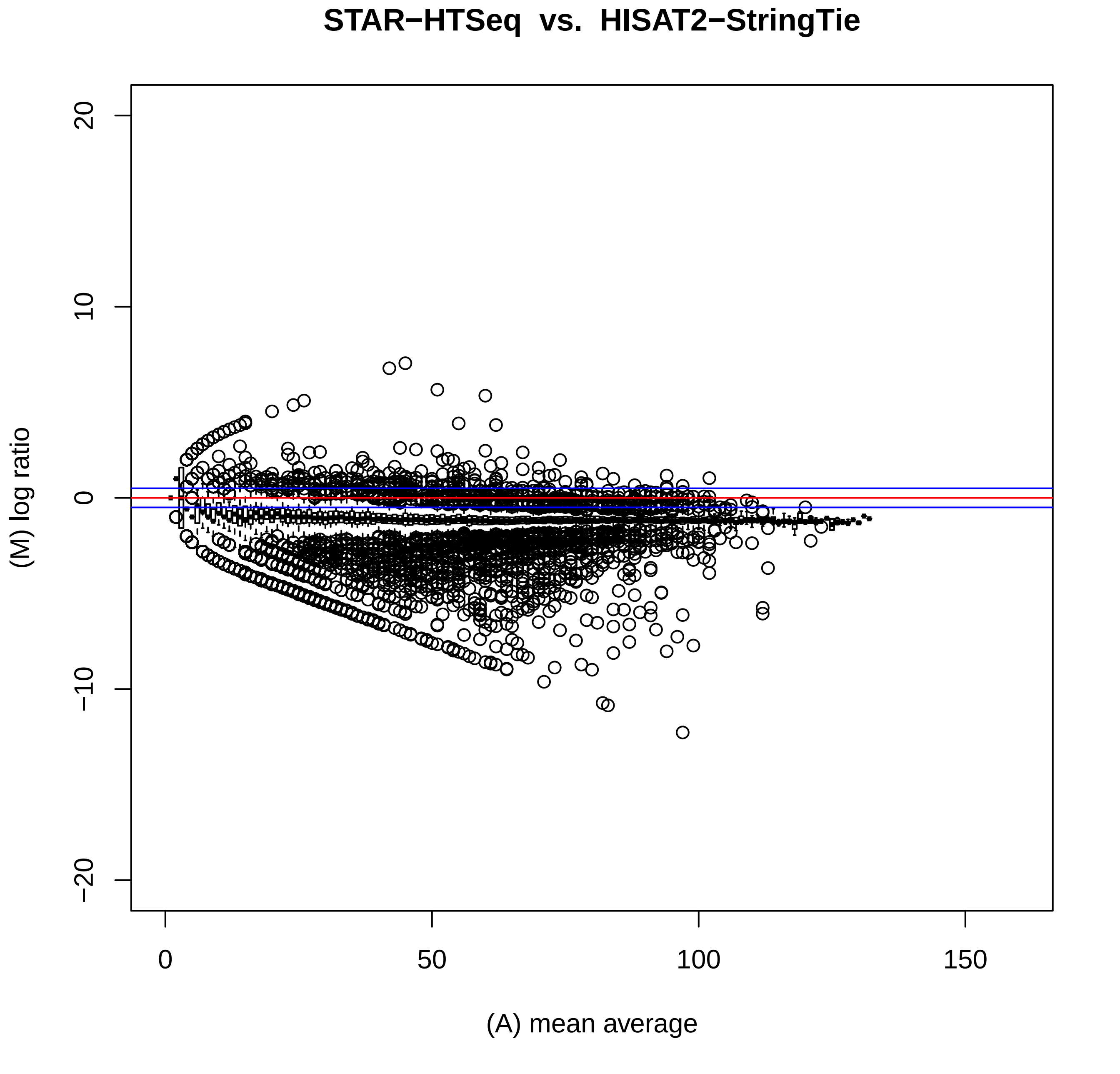
STAR−HTSeq vs. HISAT2−StringTie
20
10
0
−10
−20
(M) log ratio
0
50
100
150
(A) mean average

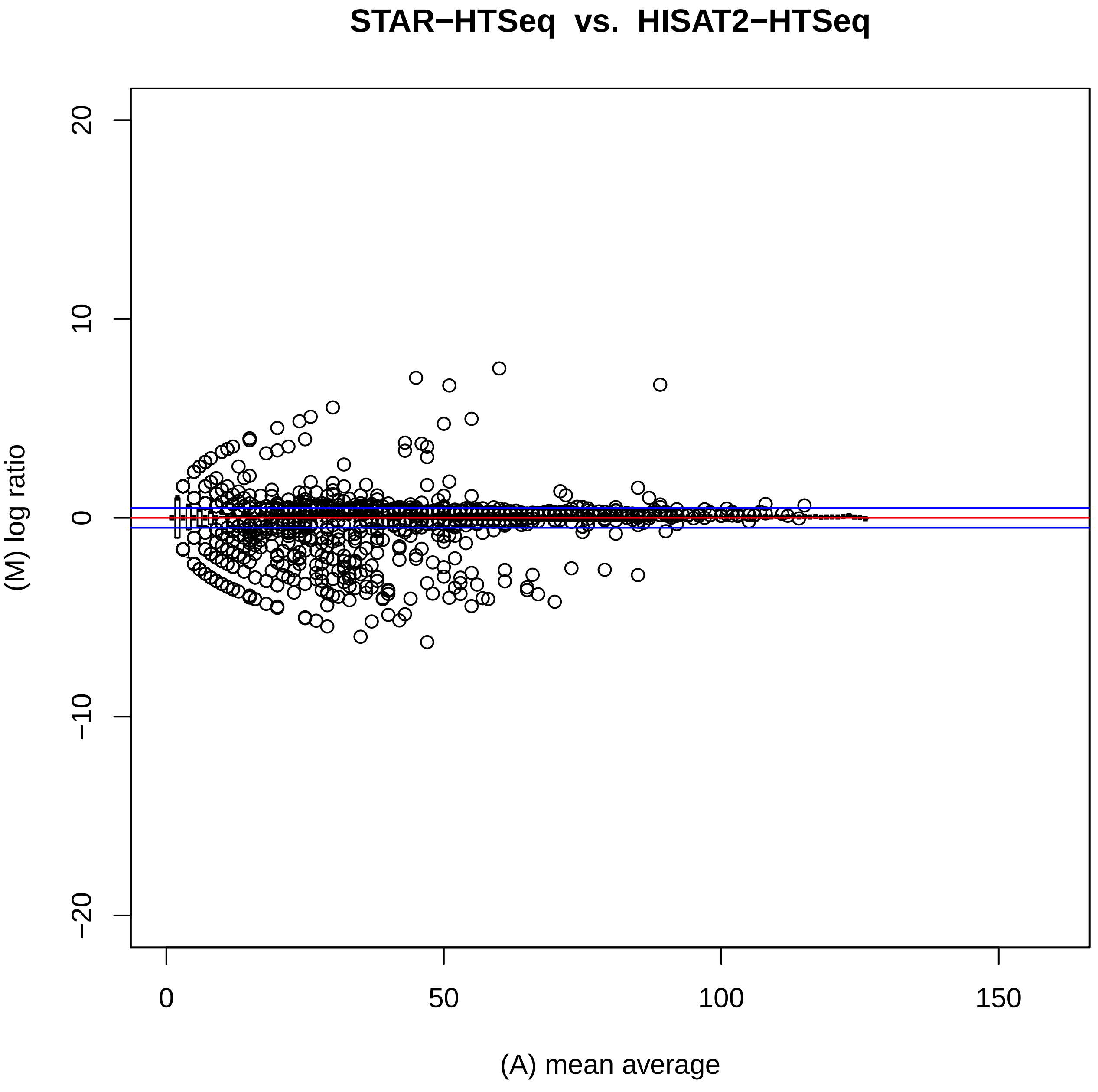
STAR−HTSeq vs. HISAT2−HTSeq
(M) log ratio
(A) mean average
20
10
0
−10
−20
0
50
100
150

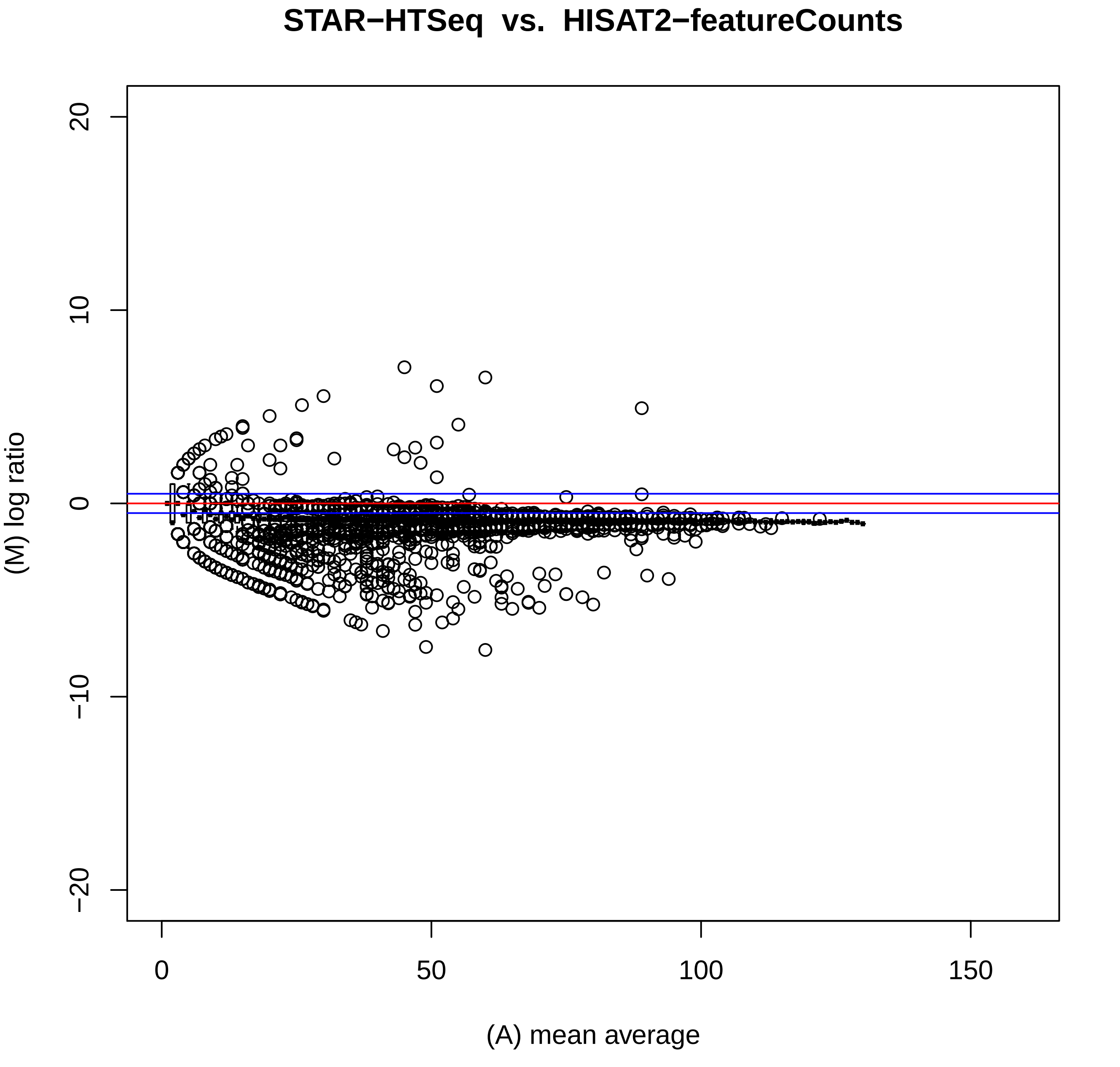
STAR−HTSeq vs. HISAT2−featureCounts
(M) log ratio
(A) mean average
20
10
0
−10
−20
0
50
100
150

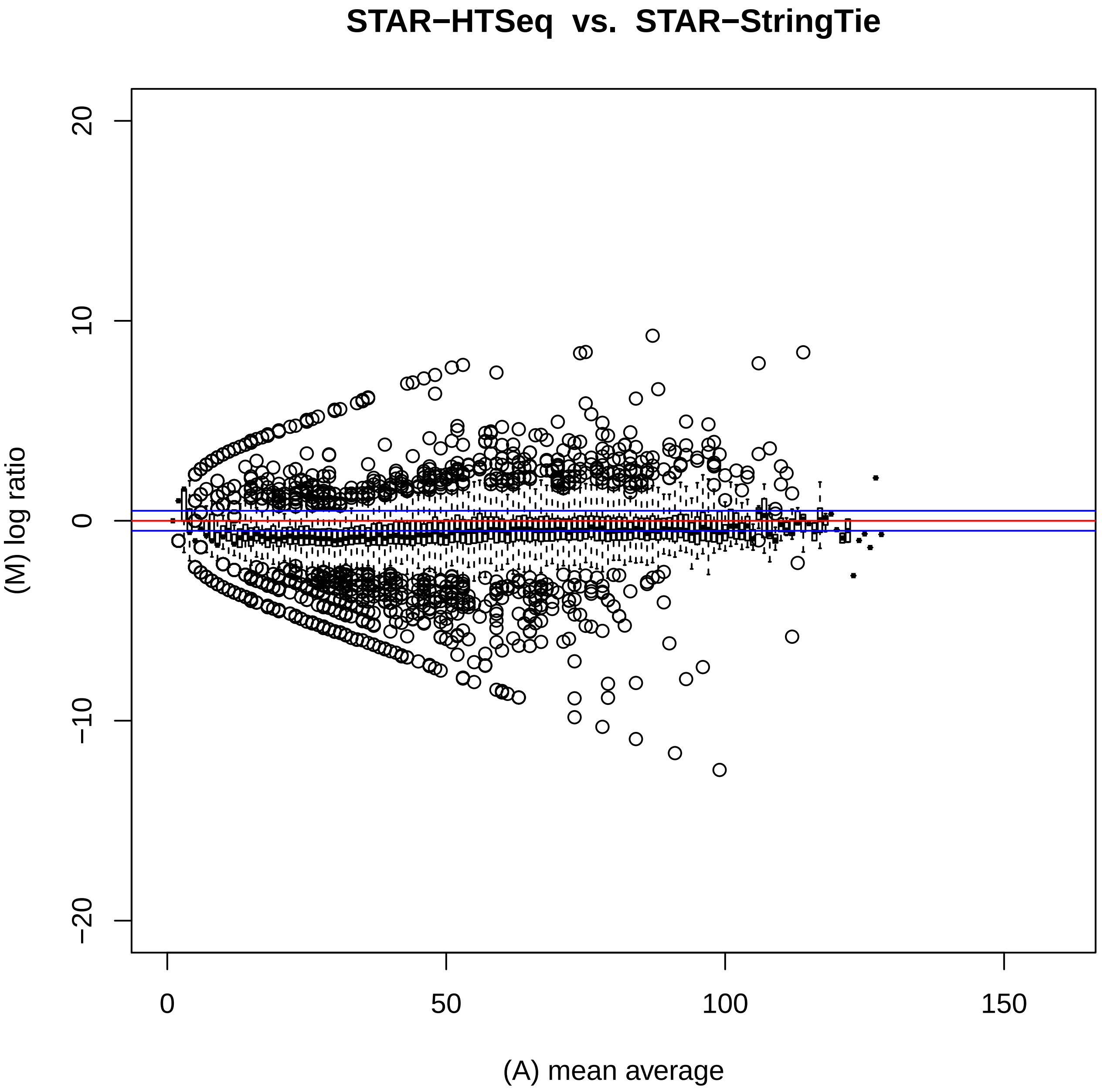
STAR−HTSeq vs. STAR−StringTie
(M) log ratio
(A) mean average
20
10
0
−10
−20
0
50
100
150

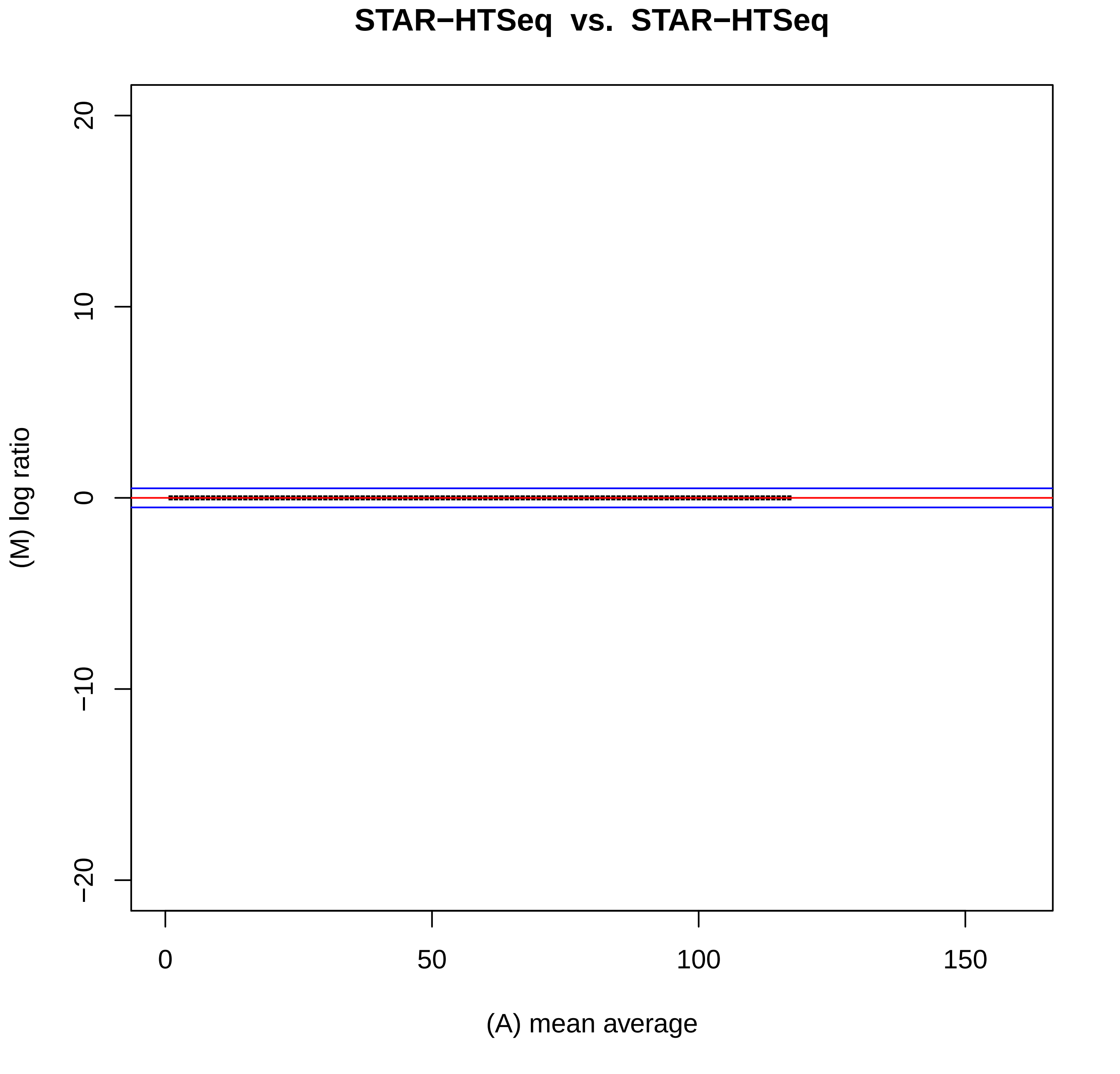
STAR−HTSeq vs. STAR−HTSeq
(M) log ratio
20
10
0
−10
−20
0
50
100
150
(A) mean average

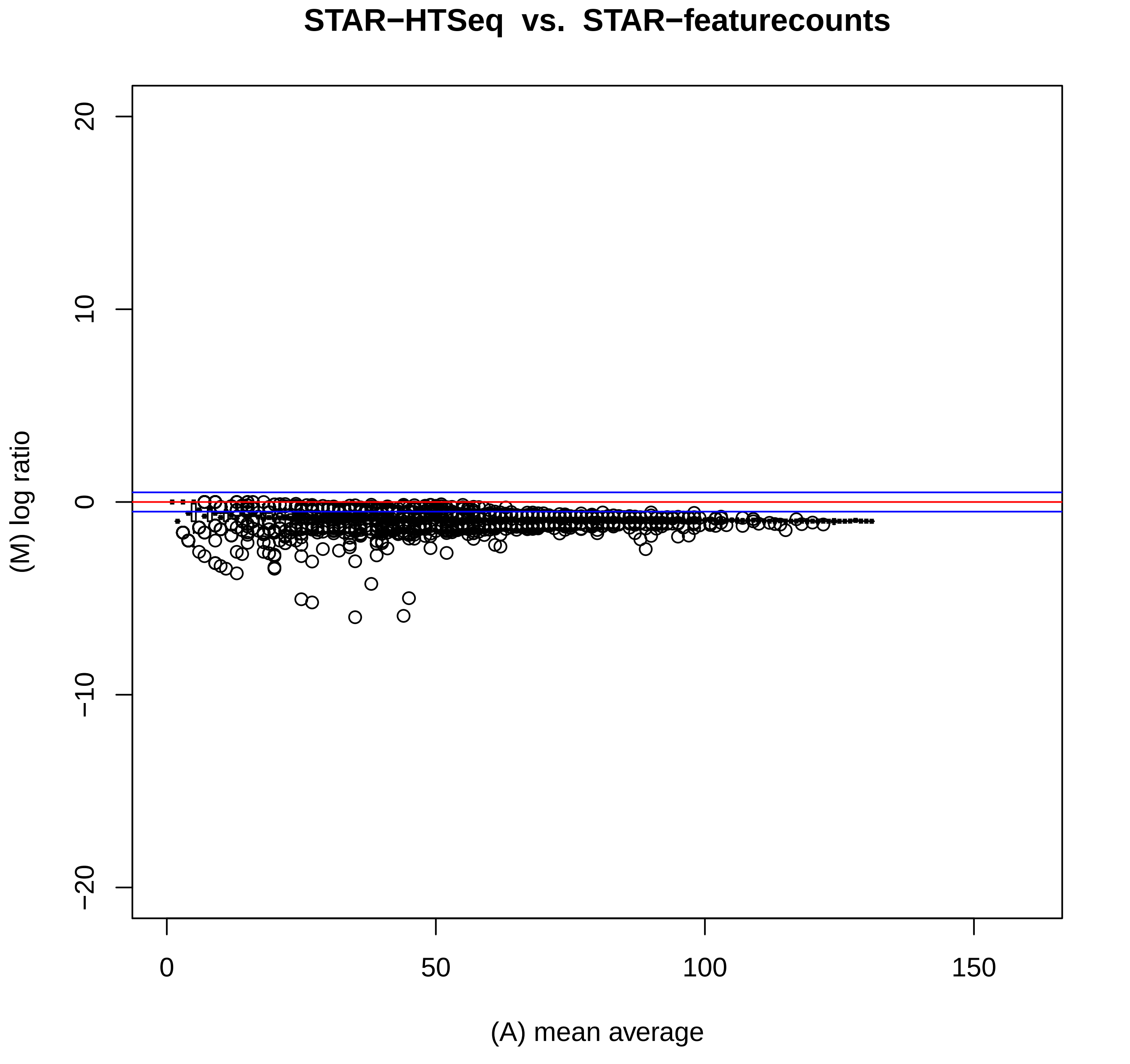

STAR−HTSeq vs. STAR−featurecounts
(M) log ratio
(A) mean average
20
10
0
−10
−20
0
50
100
150

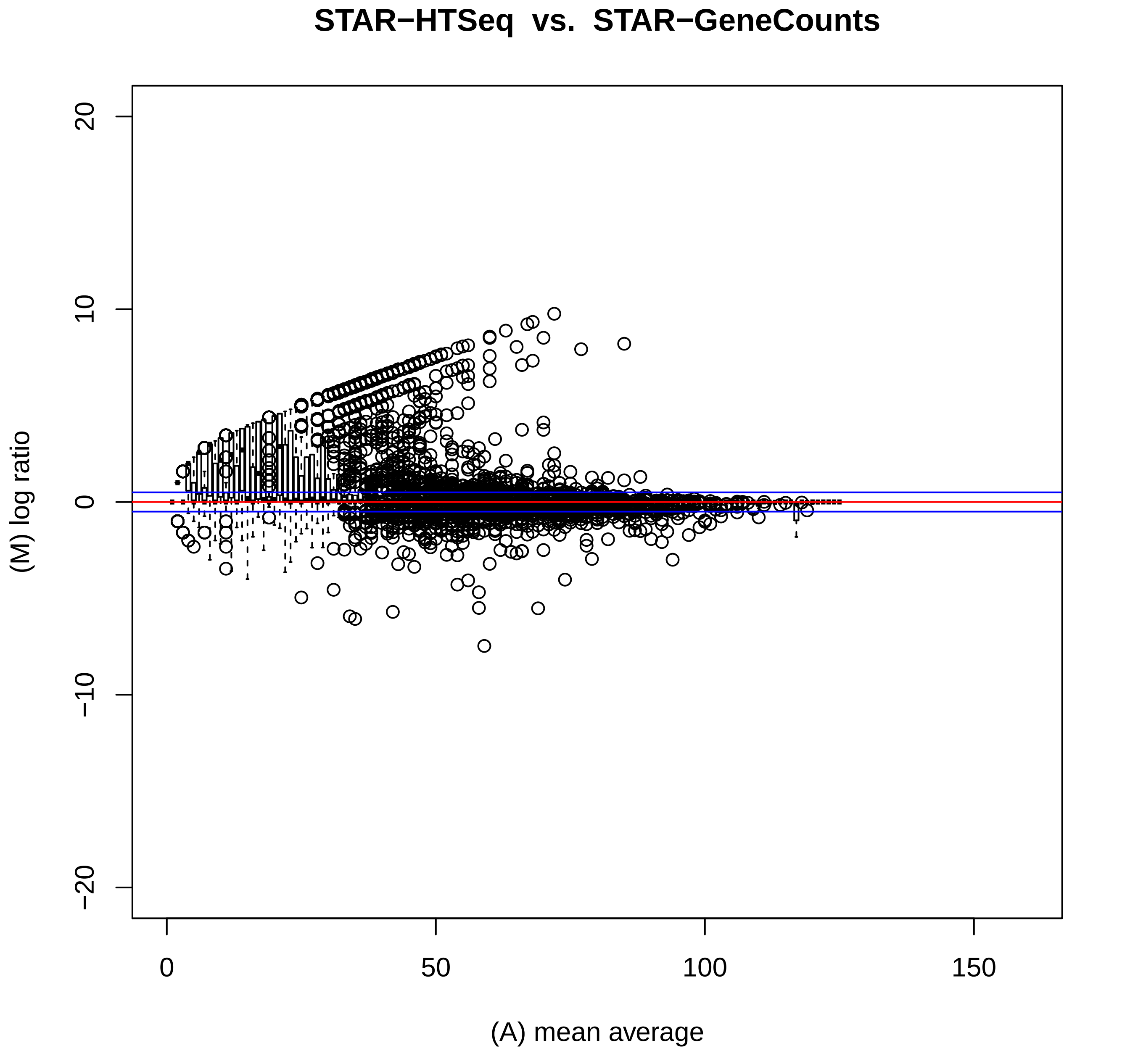
STAR−HTSeq vs. STAR−GeneCounts
(M) log ratio
(A) mean average
20
10
0
−10
−20
0
50
100
150

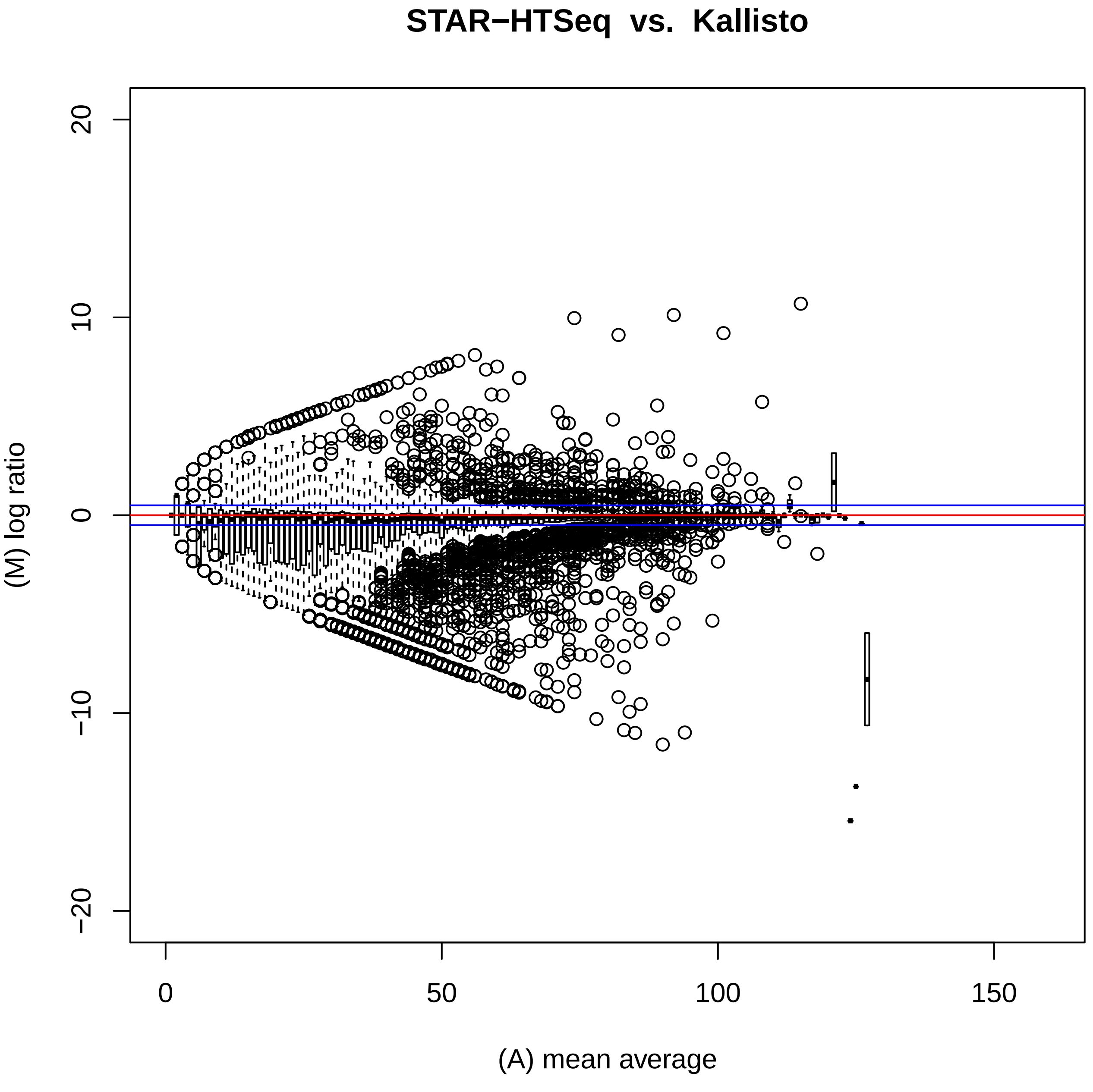

STAR−HTSeq vs. Kallisto
(M) log ratio
(A) mean average
20
10
0
−10
−20
0
50
100
150

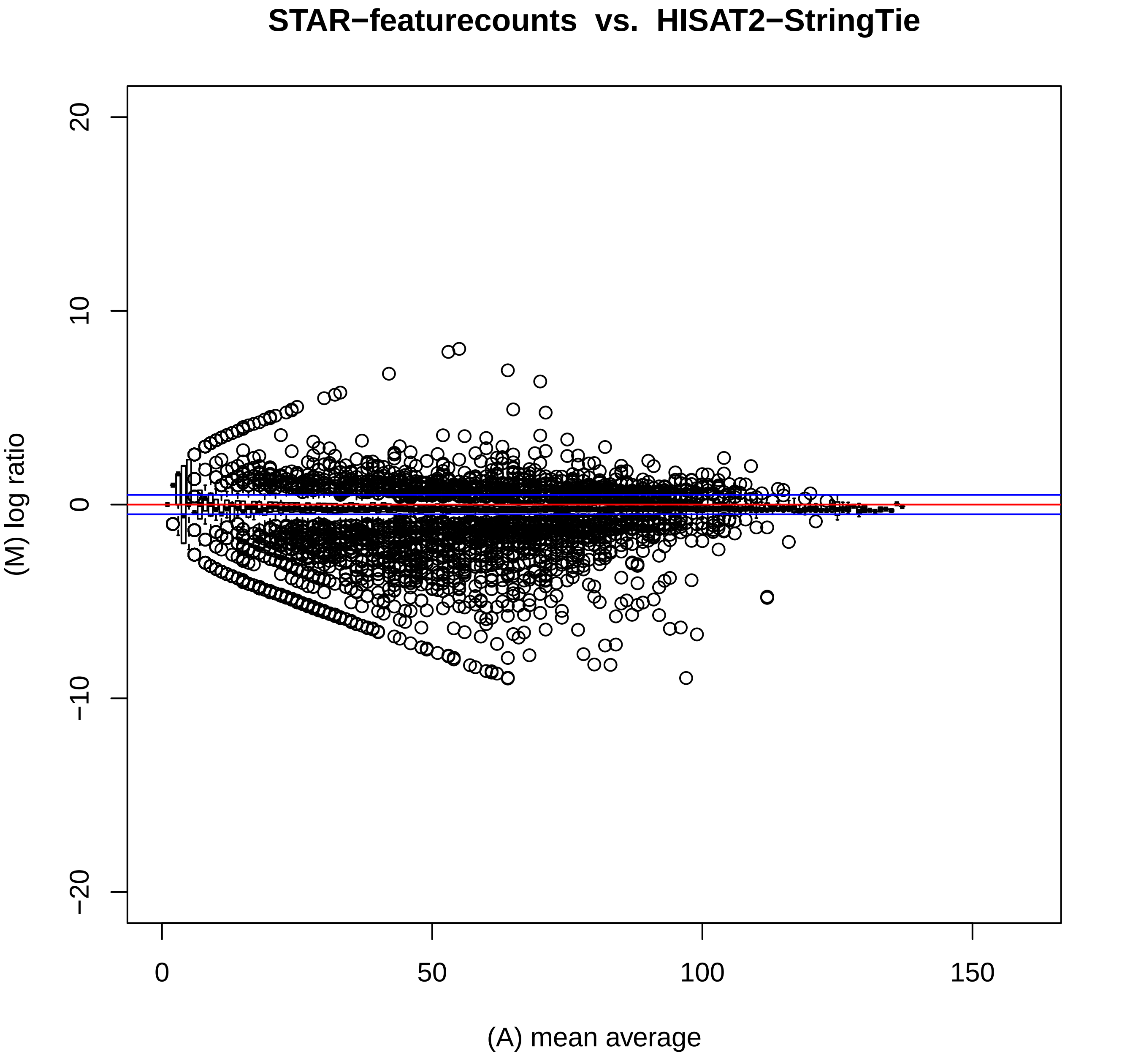

STAR−featurecounts vs. HISAT2−StringTie
20
10
0
−10
−20
(M) log ratio
0
50
100
150
(A) mean average

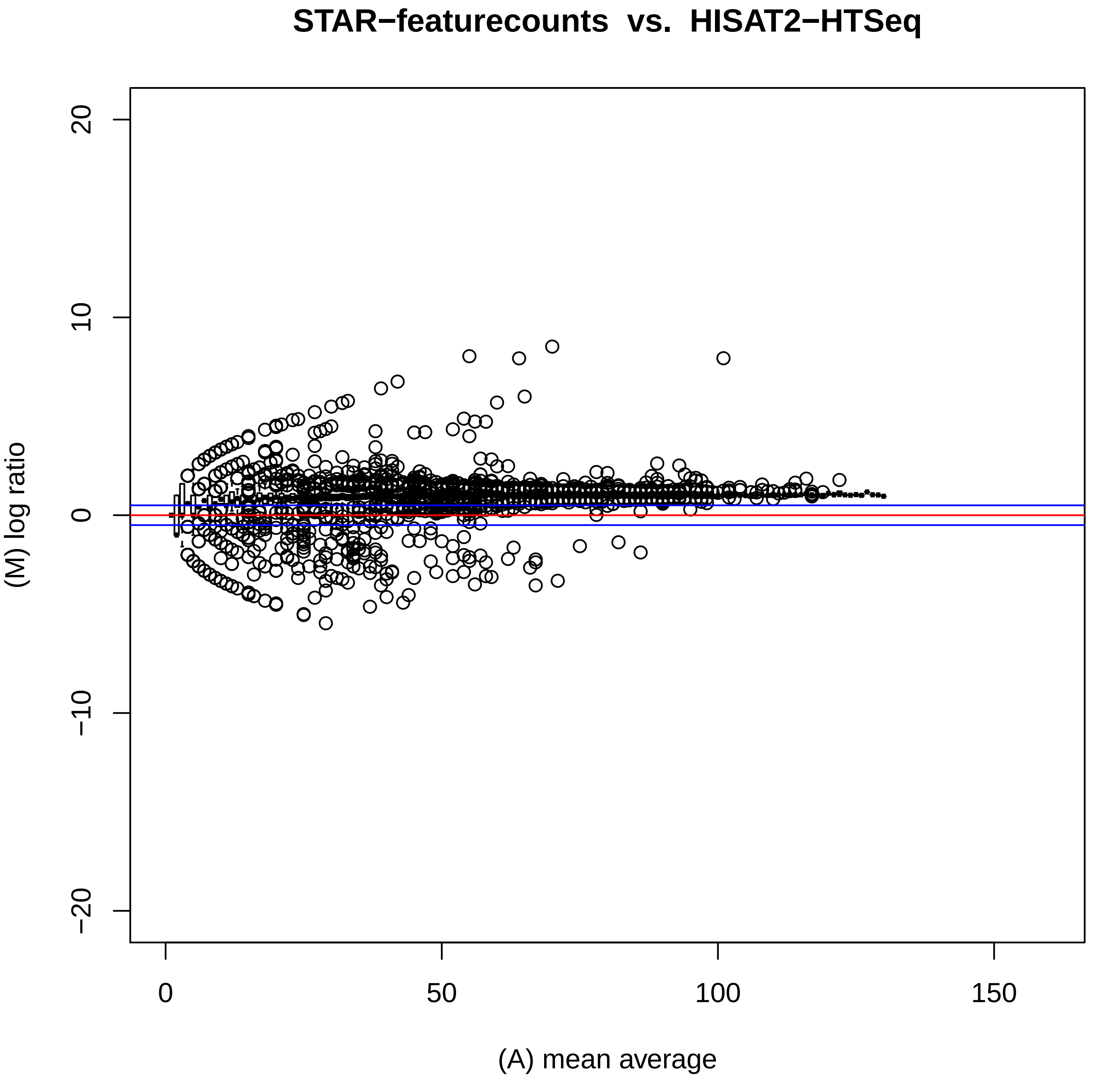

STAR−featurecounts vs. HISAT2−HTSeq
(M) log ratio
20
10
0
−10
−20
0
50
100
150
(A) mean average

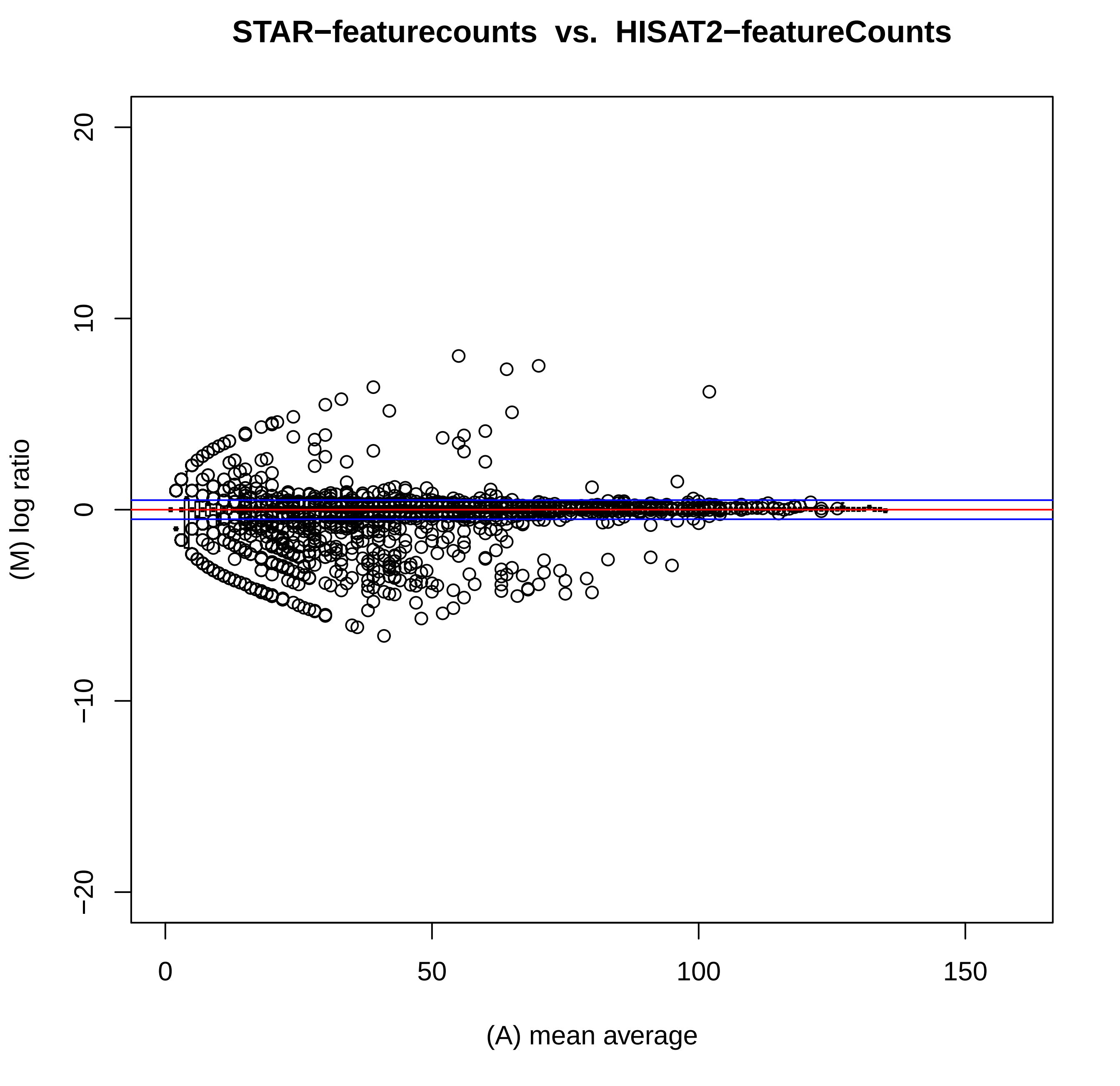

STAR−featurecounts vs. HISAT2−featureCounts
20
10
0
−10
−20
(M) log ratio
0
50
100
150
(A) mean average

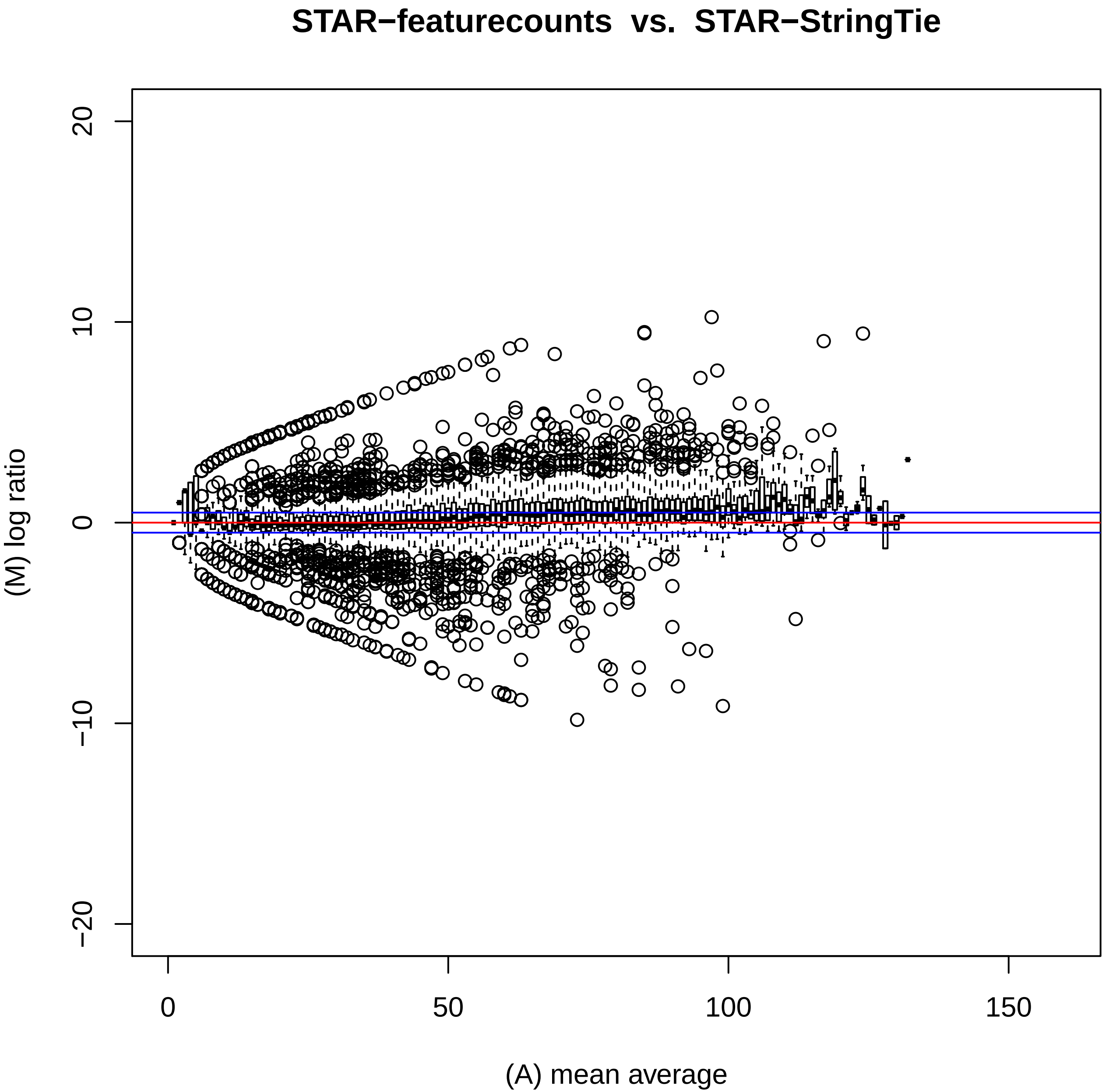
STAR−featurecounts vs. STAR−StringTie
(M) log ratio
(A) mean average
20
10
0
−10
−20
0
50
100
150

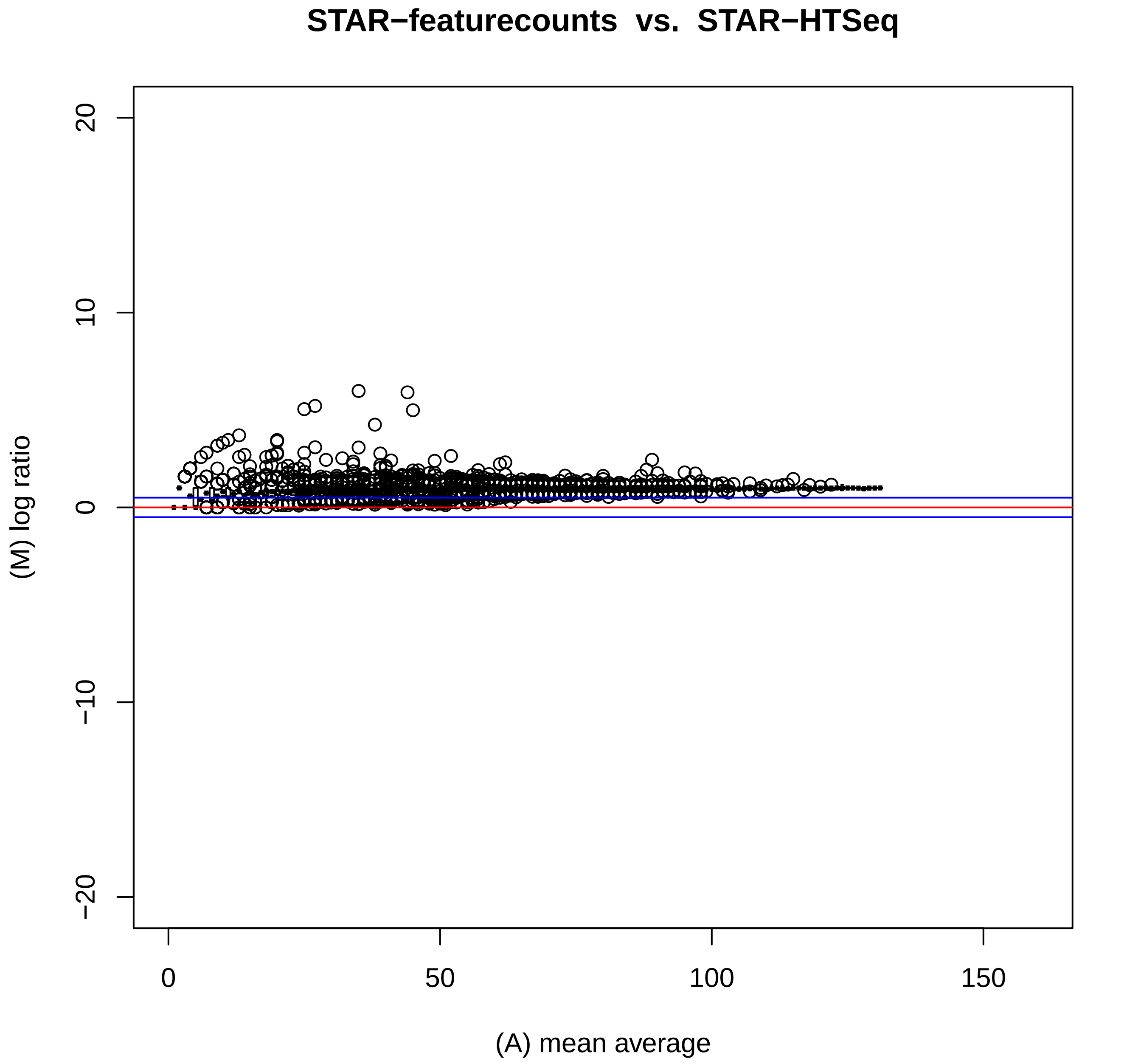
STAR−featurecounts vs. STAR−HTSeq
(M) log ratio
(A) mean average
20
10
0
−10
−20
0
50
100
150

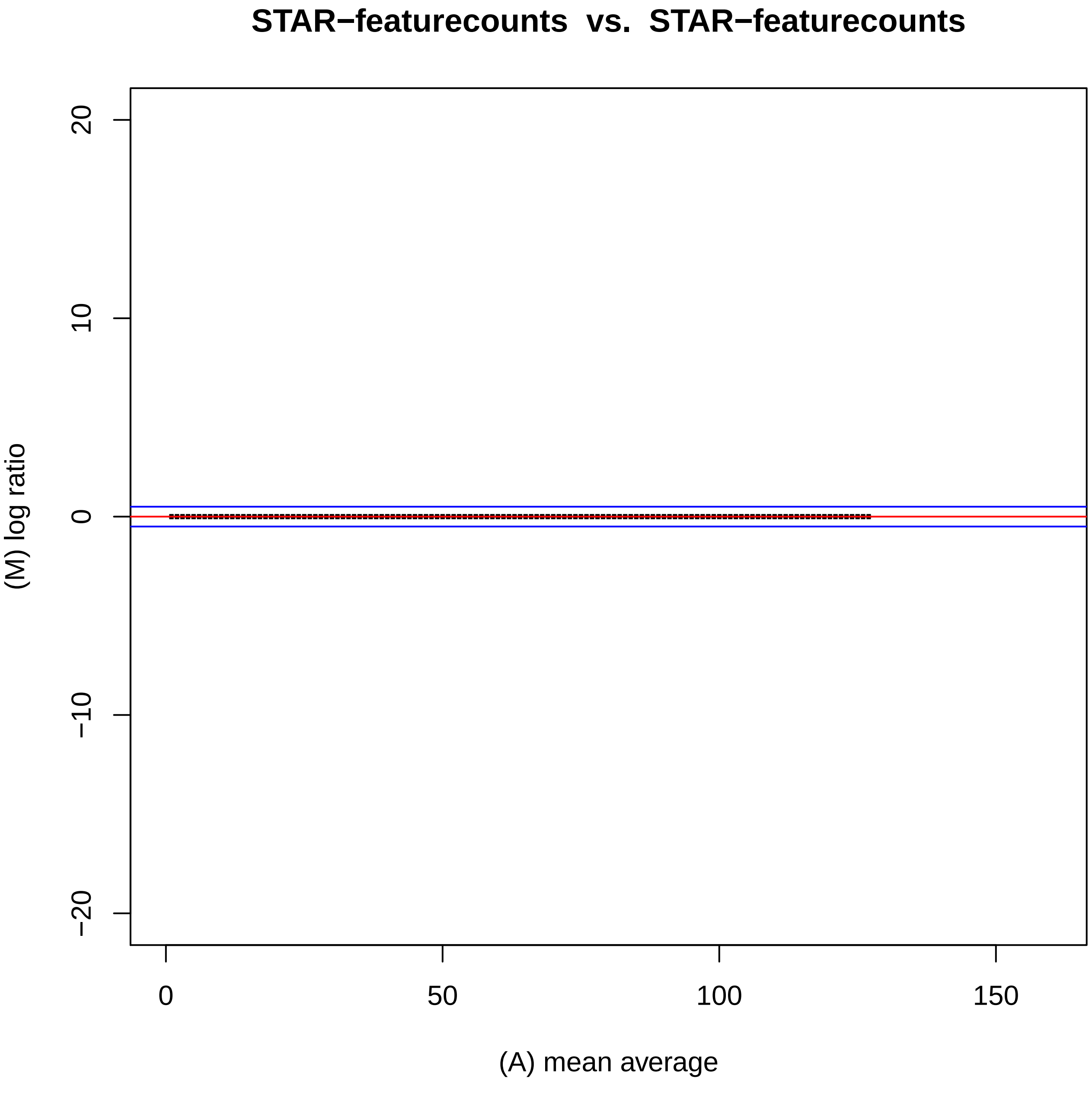
STAR−featurecounts vs. STAR−featurecounts
(M) log ratio
20
10
0
−10
−20
0
50
100
150
(A) mean average

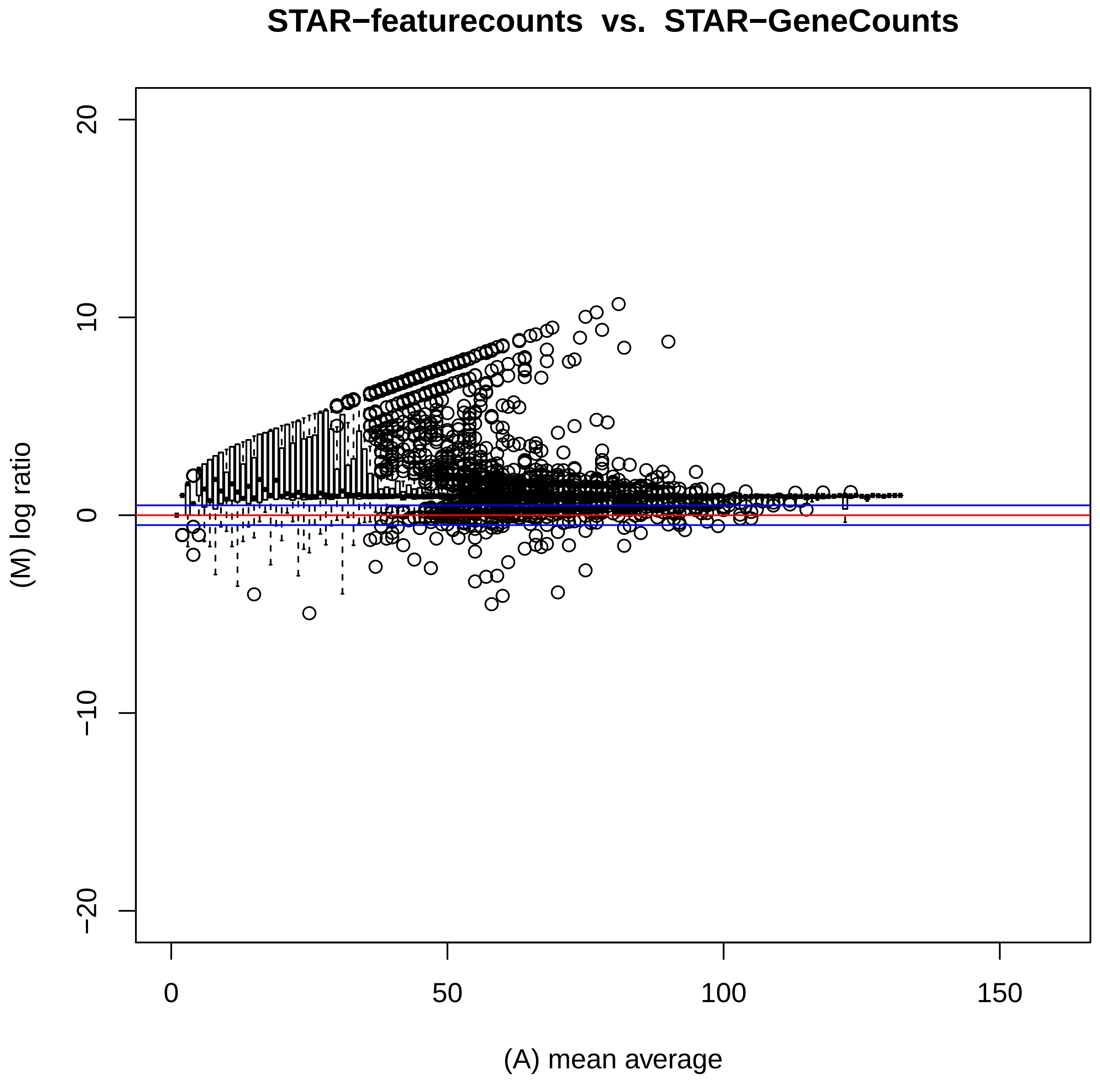

STAR−featurecounts vs. STAR−GeneCounts
(M) log ratio
(A) mean average
20
10
0
−10
−20
0
50
100
150

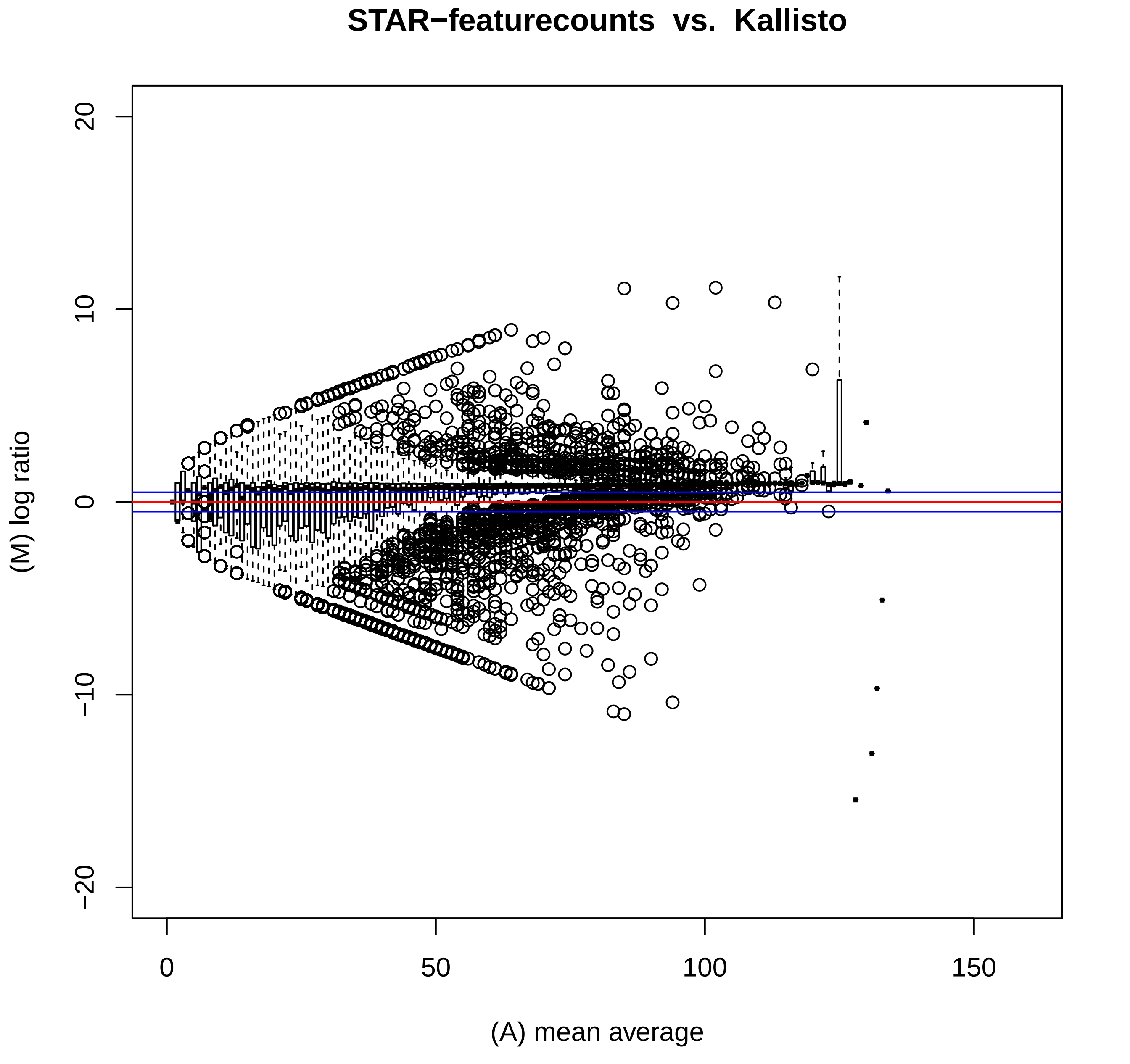

STAR−featurecounts vs. Kallisto
(M) log ratio
(A) mean average
20
10
0
−10
−20
0
50
100
150

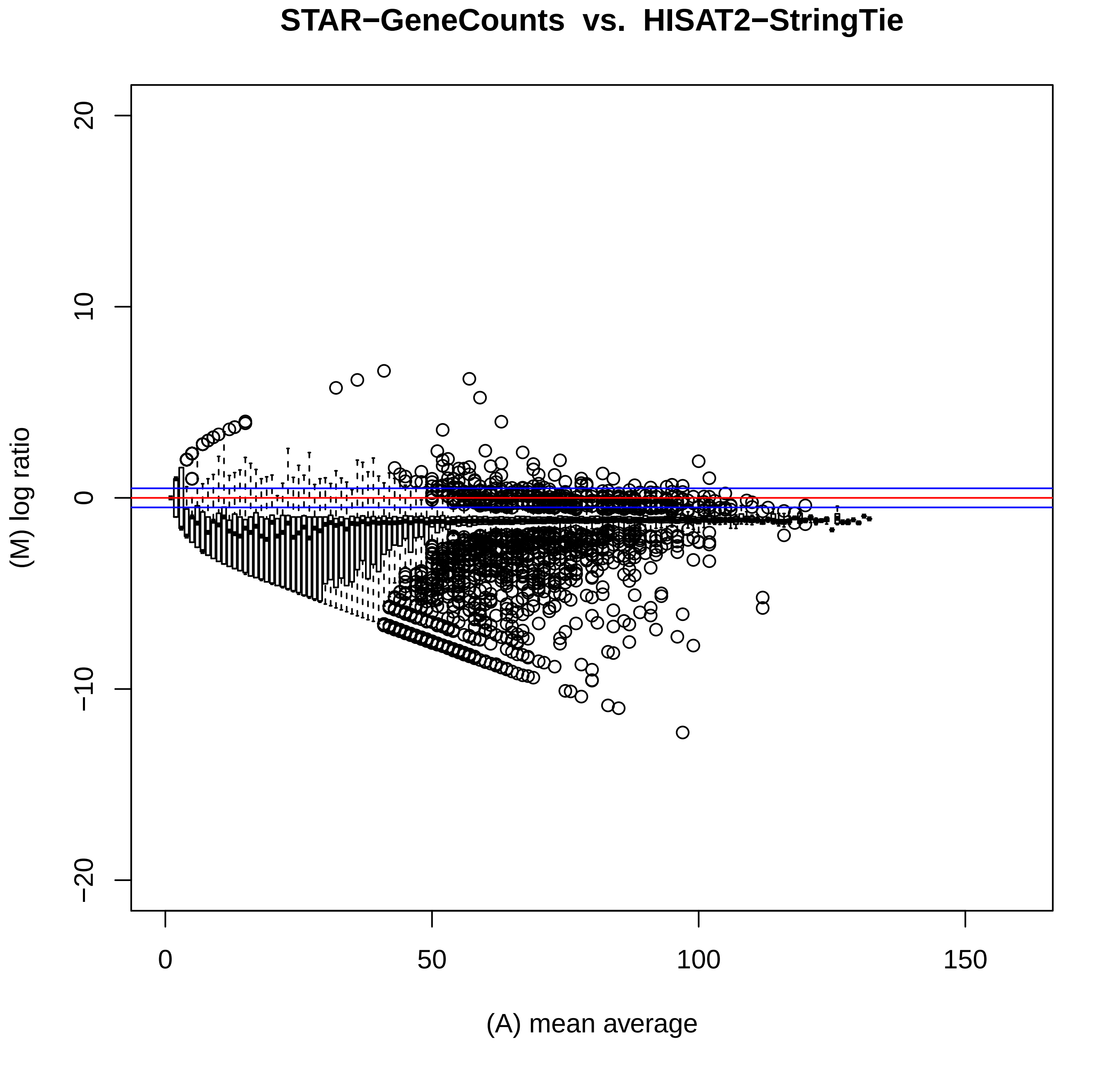

STAR−GeneCounts vs. HISAT2−StringTie
(M) log ratio
20
10
0
−10
−20
0
50
100
150
(A) mean average

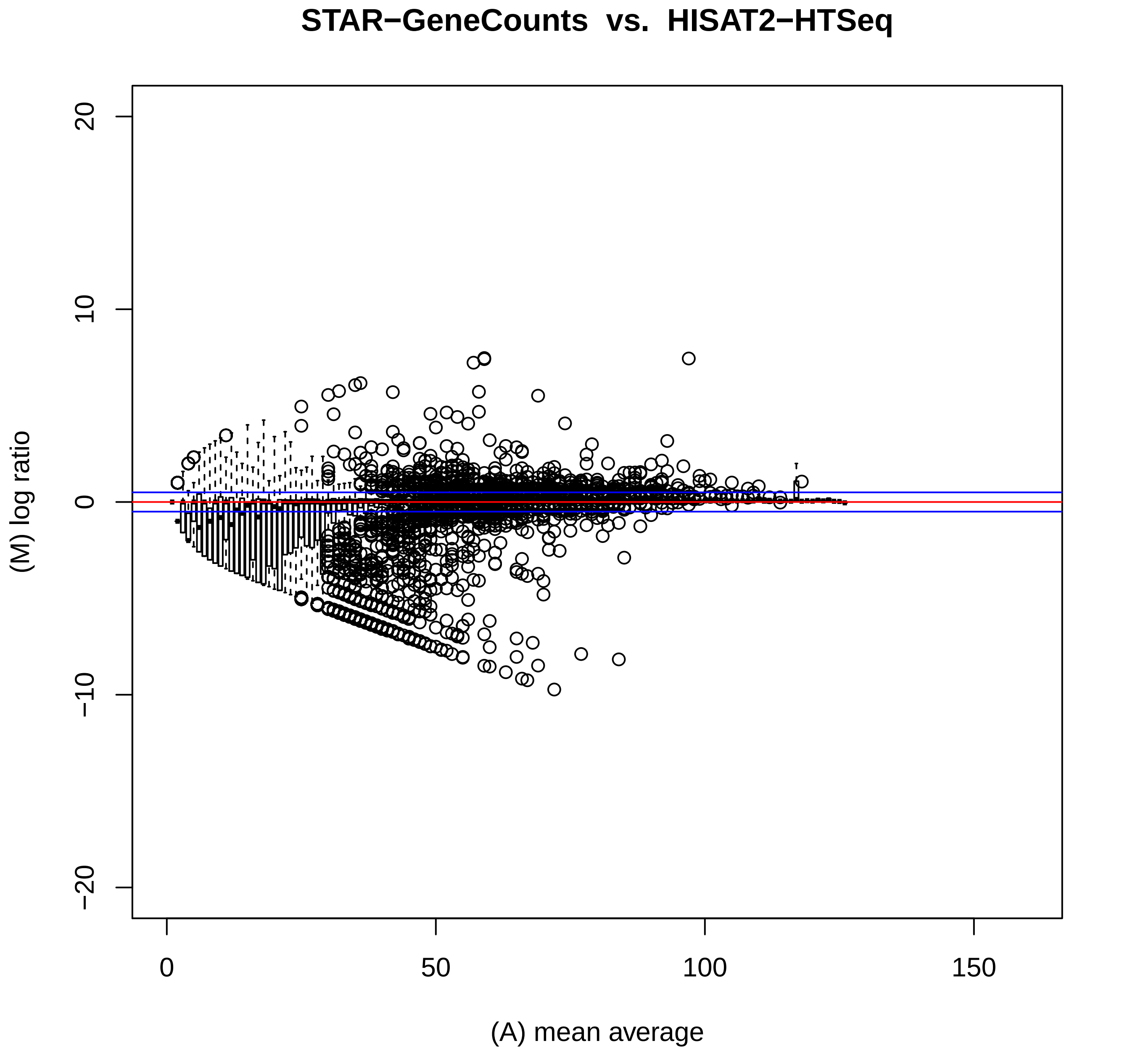

STAR−GeneCounts vs. HISAT2−HTSeq
(M) log ratio
(A) mean average
20
10
0
−10
−20
0
50
100
150

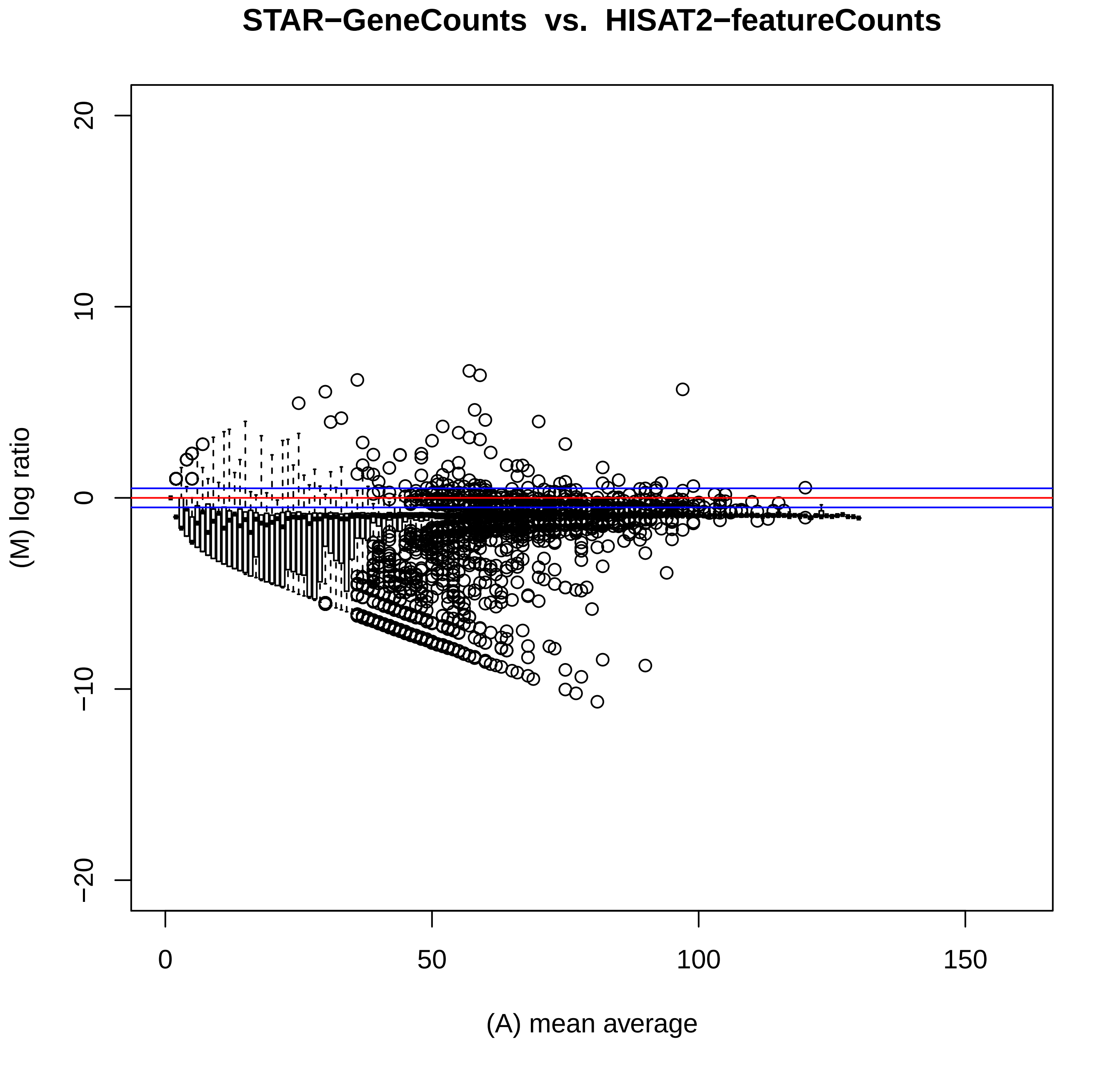

STAR−GeneCounts vs. HISAT2−featureCounts
(M) log ratio
20
10
0
−10
−20
0
50
100
150
(A) mean average

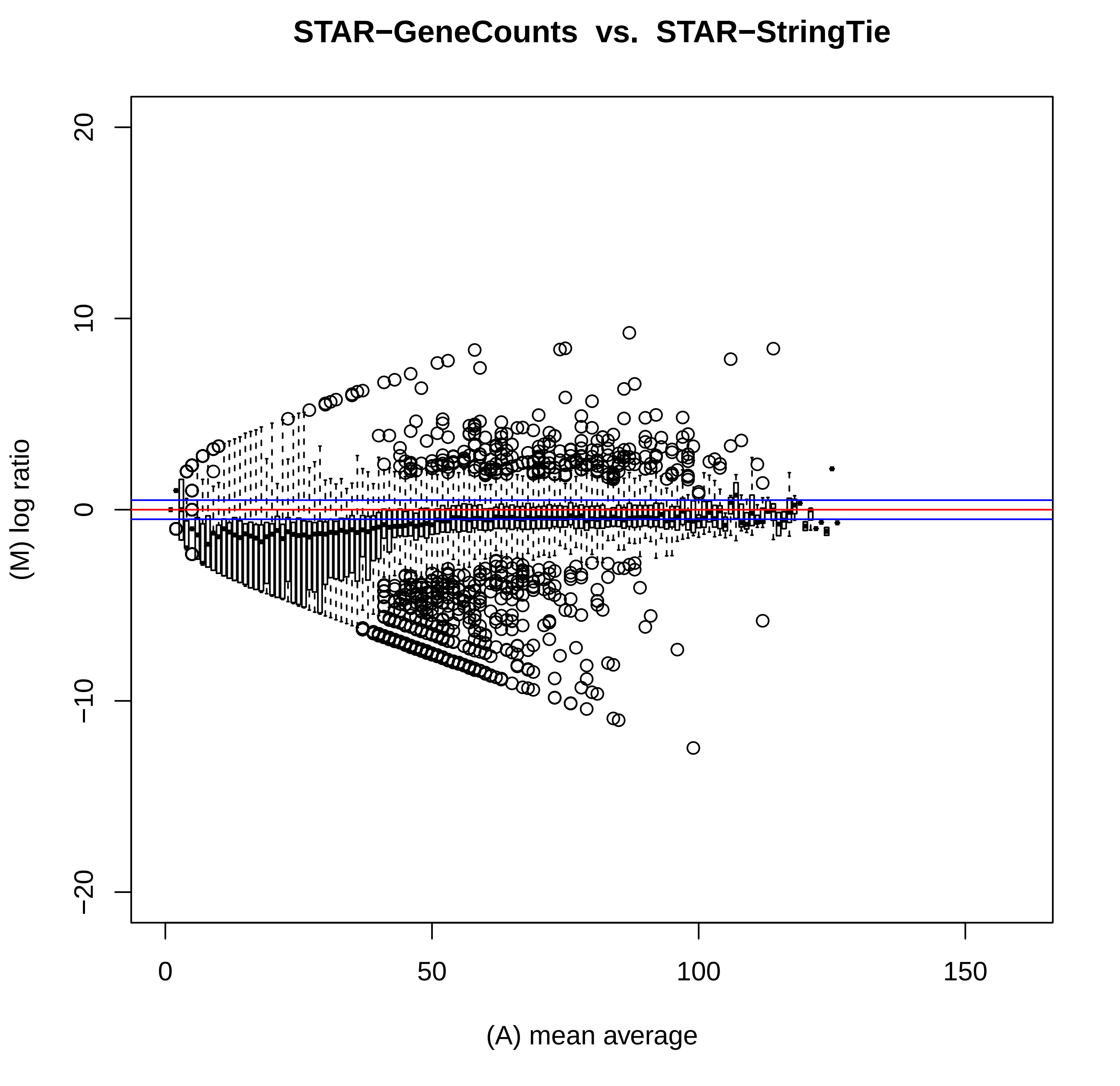
STAR−GeneCounts vs. STAR−StringTie
(M) log ratio
(A) mean average
20
10
0
−10
−20
0
50
100
150

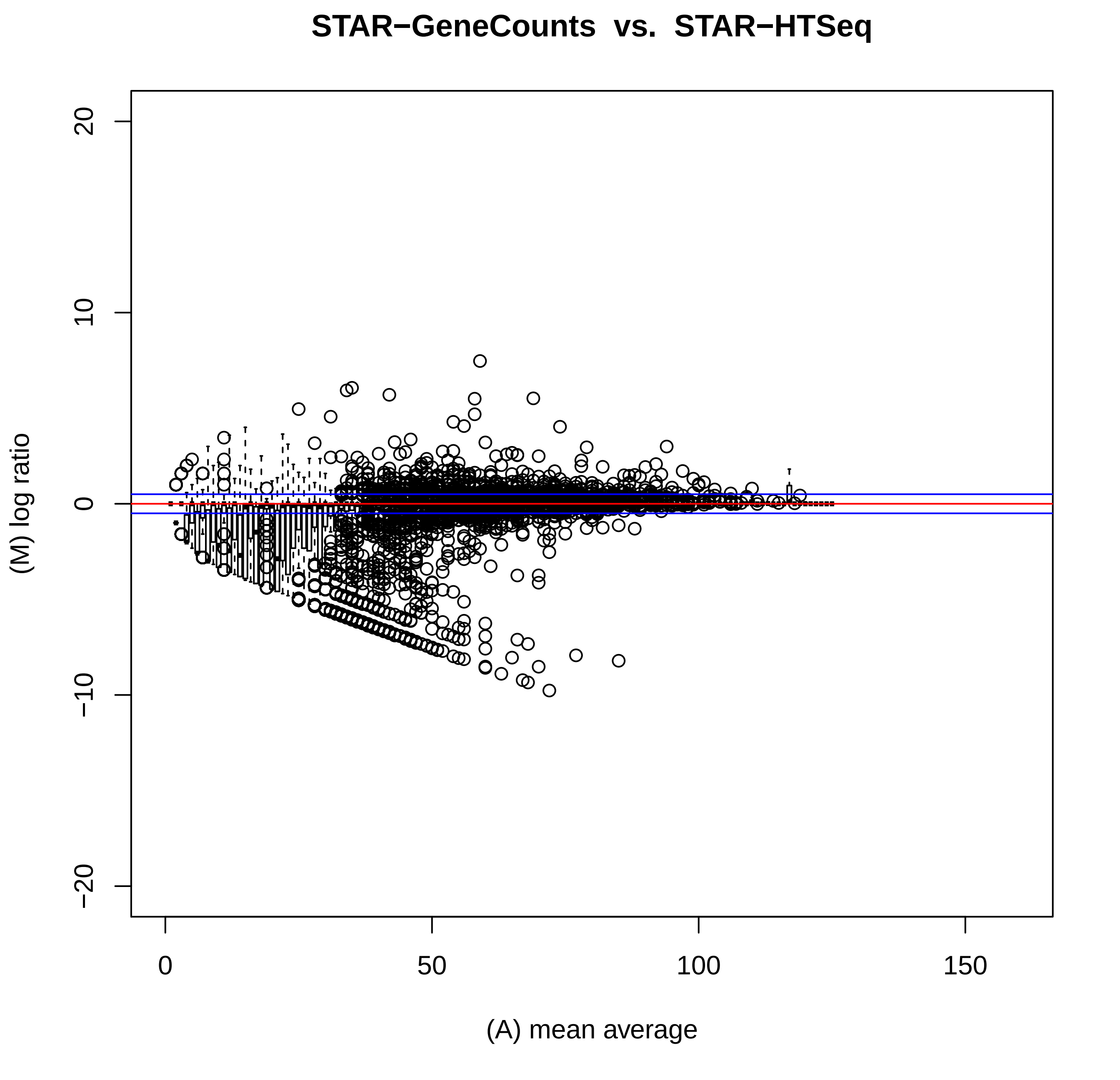
STAR−GeneCounts vs. STAR−HTSeq
(M) log ratio
(A) mean average
20
10
0
−10
−20
0
50
100
150

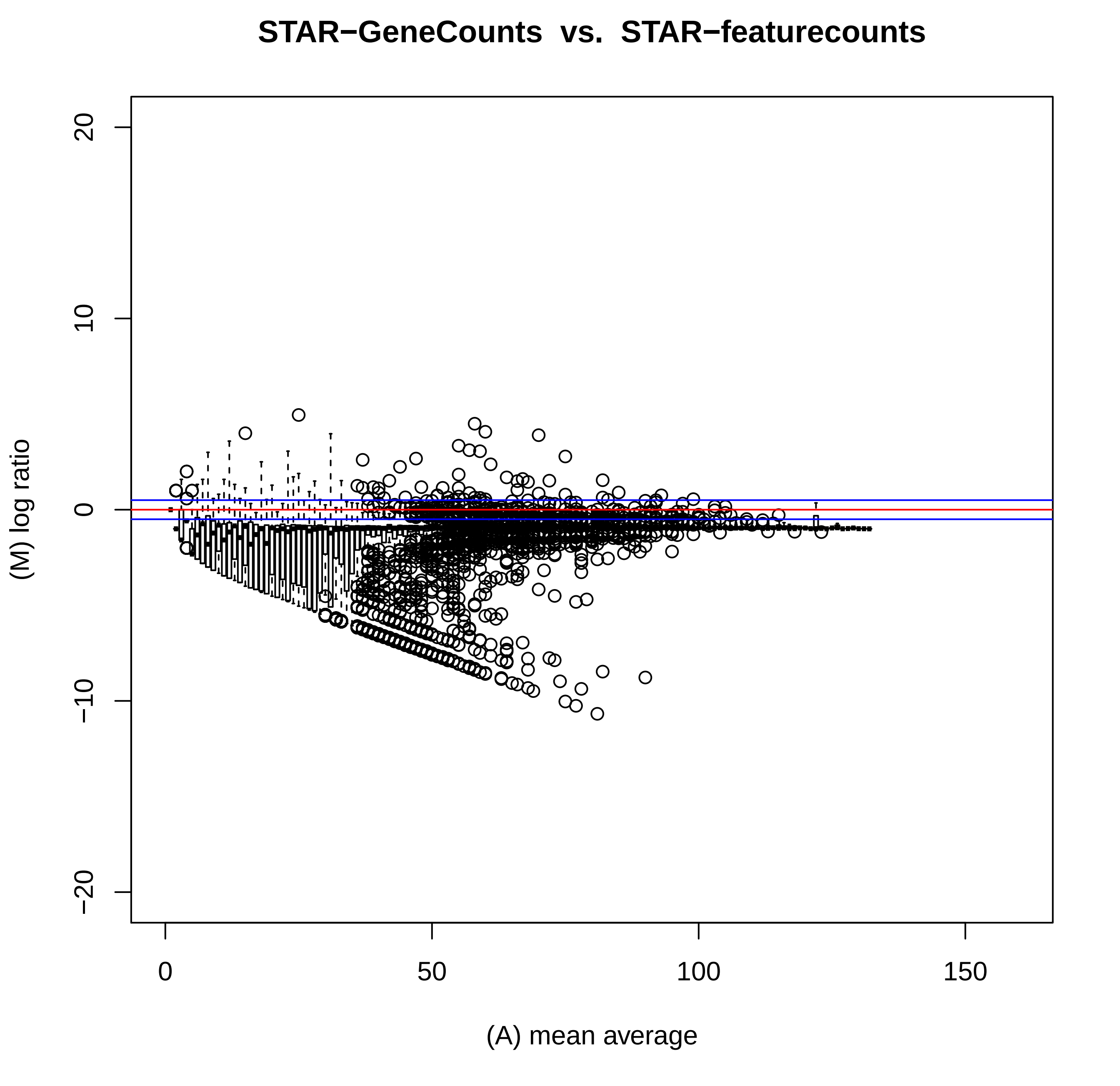

STAR−GeneCounts vs. STAR−featurecounts
(M) log ratio
20
10
0
−10
−20
0
50
100
150
(A) mean average

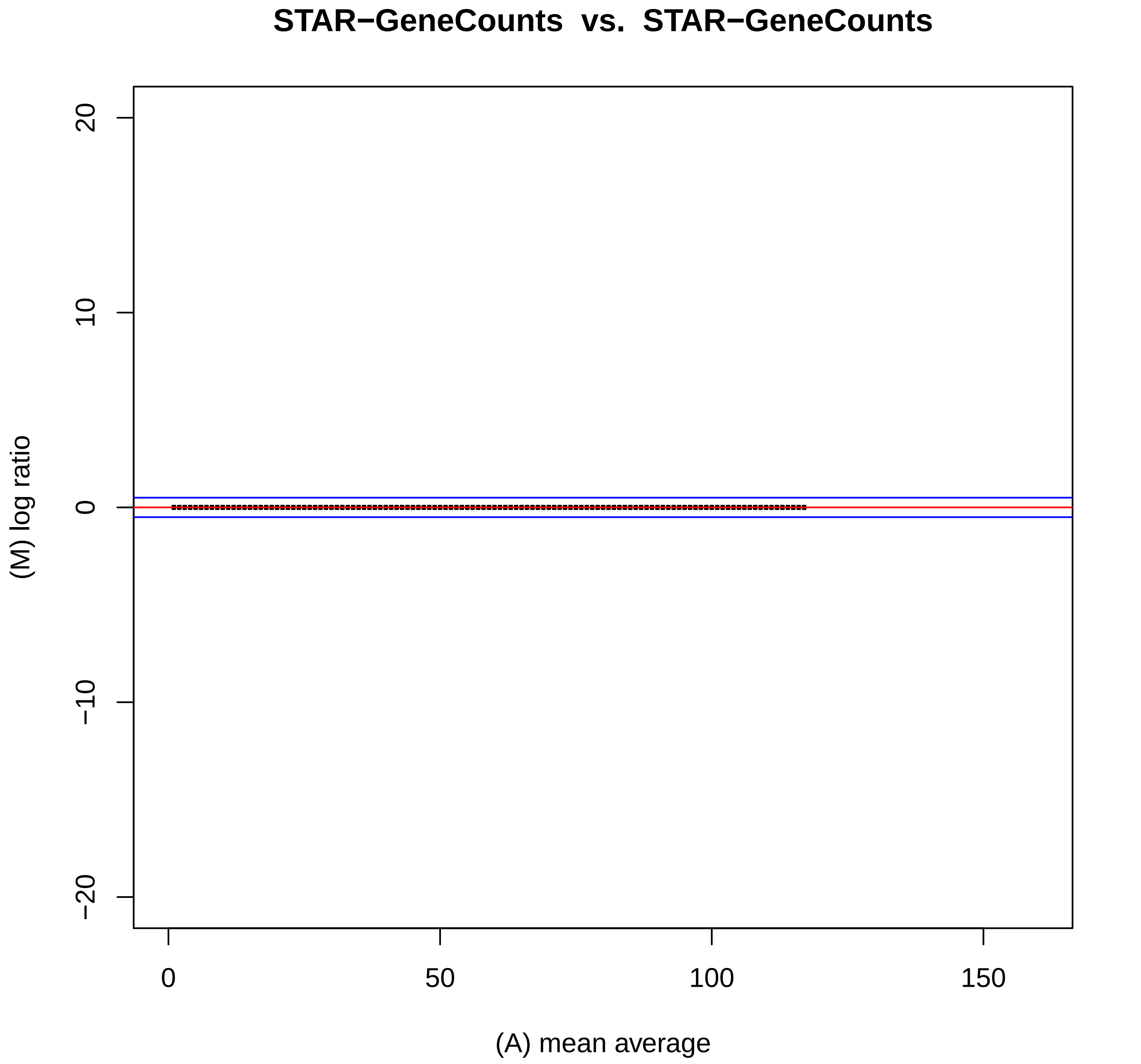
STAR−GeneCounts vs. STAR−GeneCounts
20
10
0
−10
−20
(M) log ratio
0
50
100
150
(A) mean average

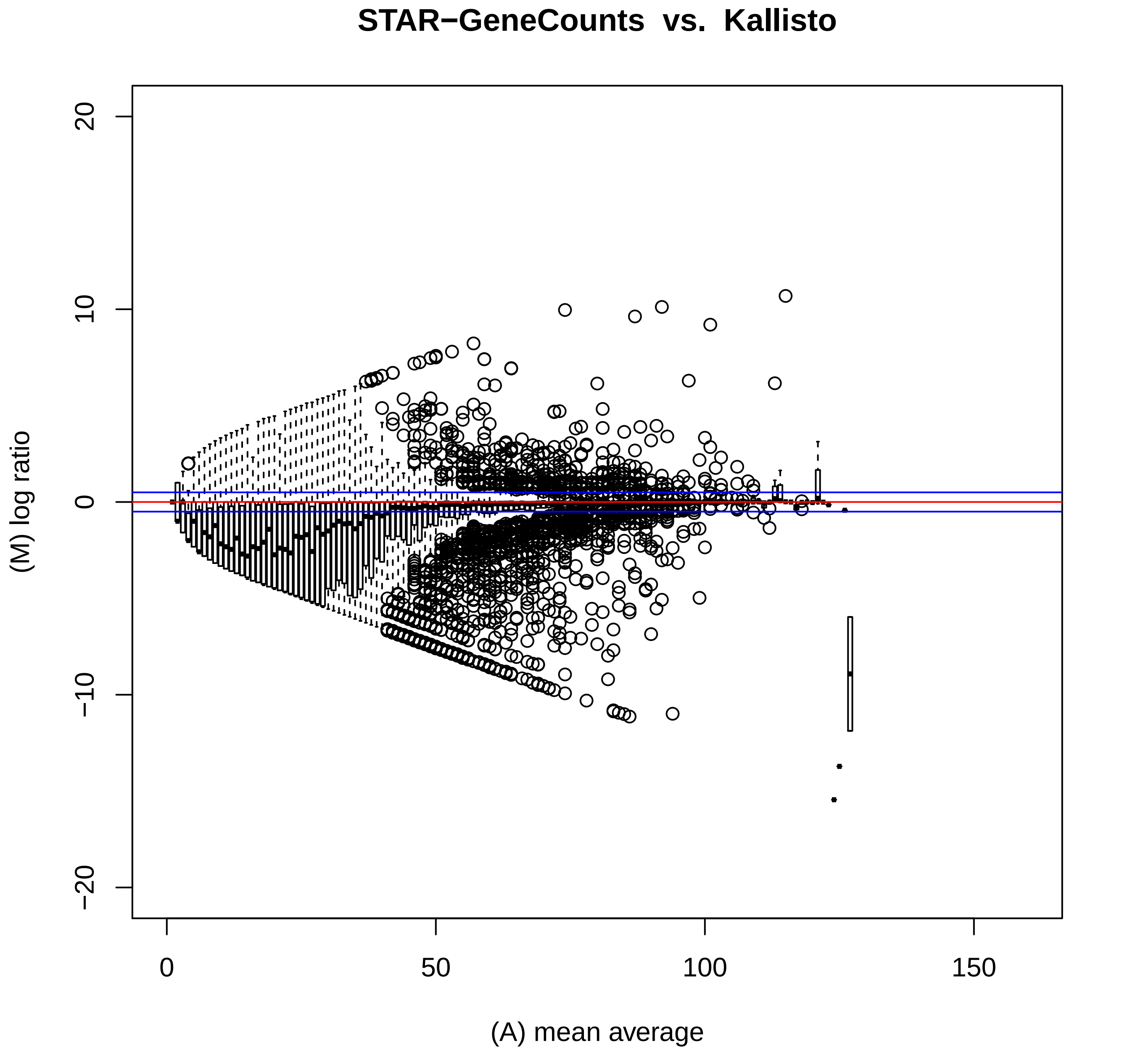
STAR−GeneCounts vs. Kallisto
(M) log ratio
(A) mean average
20
10
0
−10
−20
0
50
100
150

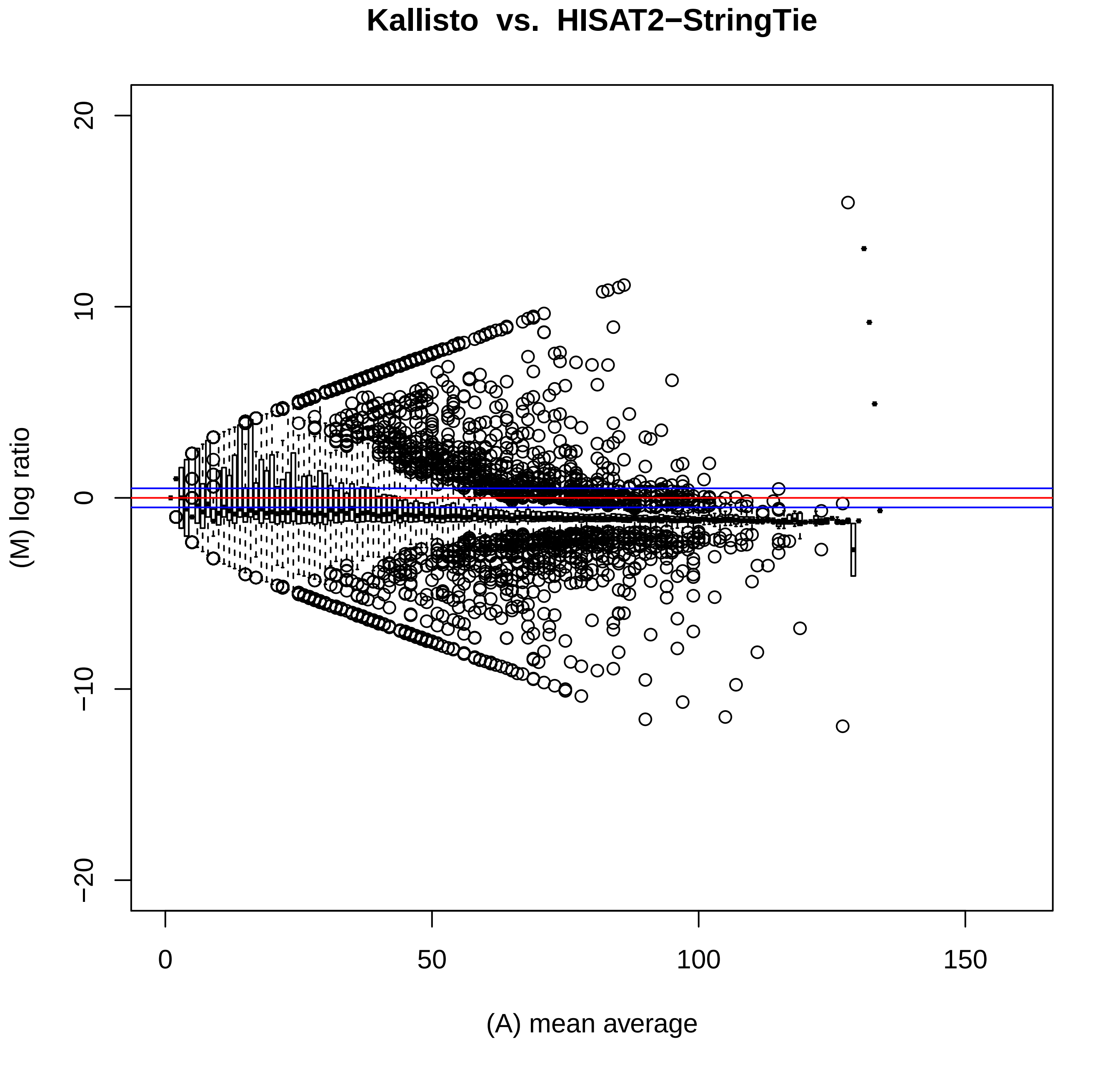
Kallisto vs. HISAT2−StringTie
(M) log ratio
(A) mean average
20
10
0
−10
−20
0
50
100
150

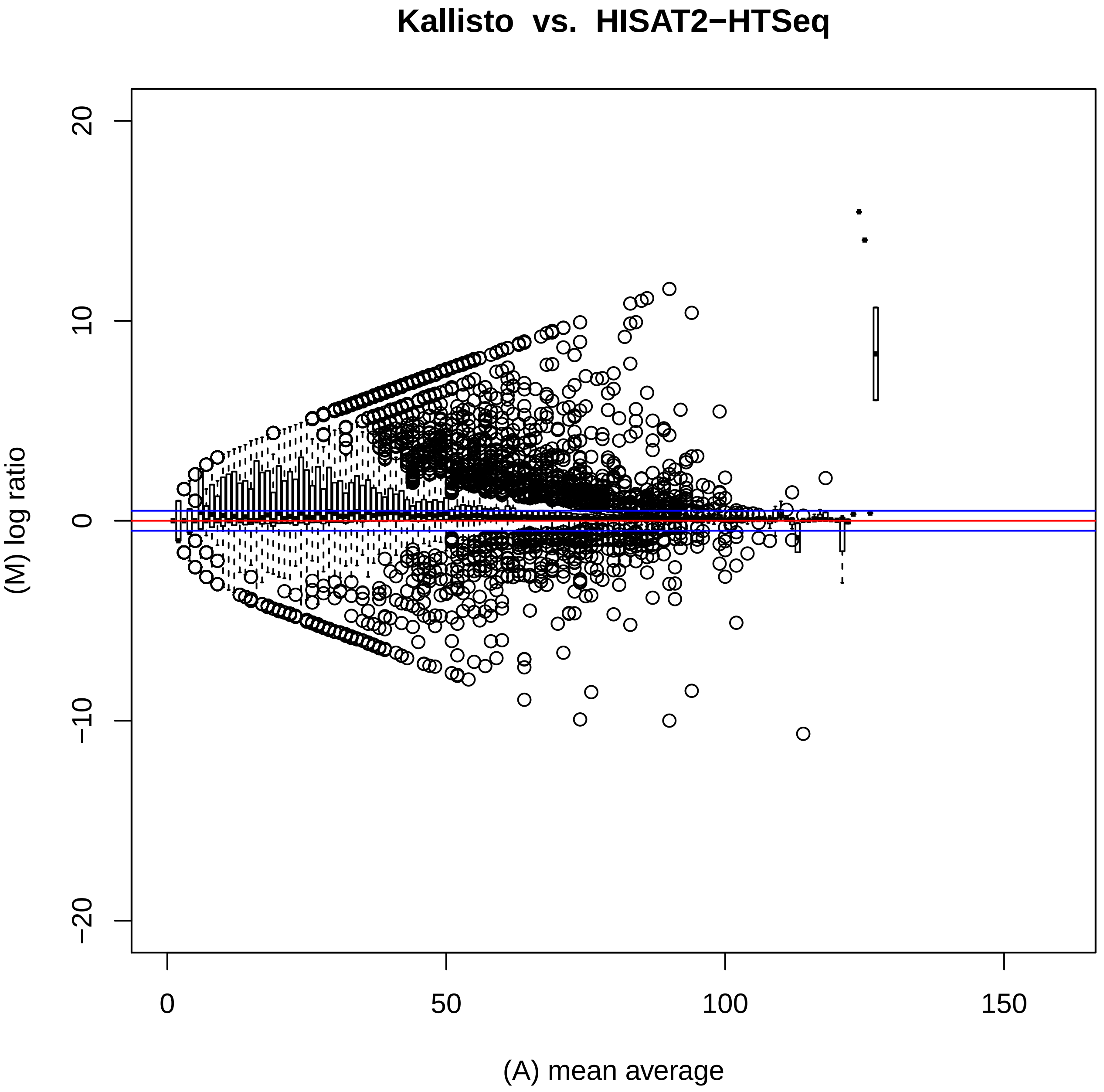
Kallisto vs. HISAT2−HTSeq
(M) log ratio
(A) mean average
20
10
0
−10
−20
0
50
100
150

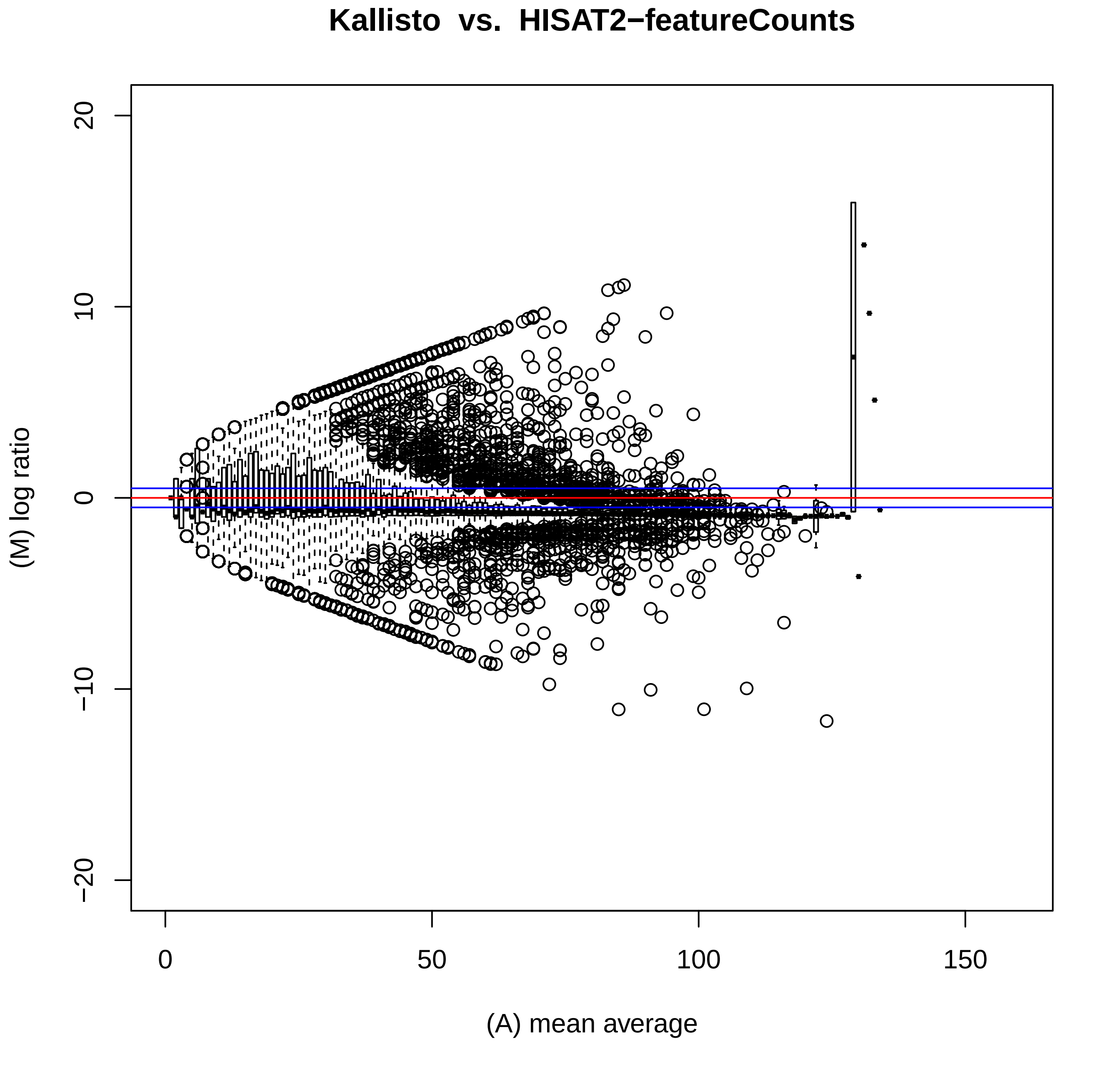

Kallisto vs. HISAT2−featureCounts
(M) log ratio
(A) mean average
20
10
0
−10
−20
0
50
100
150

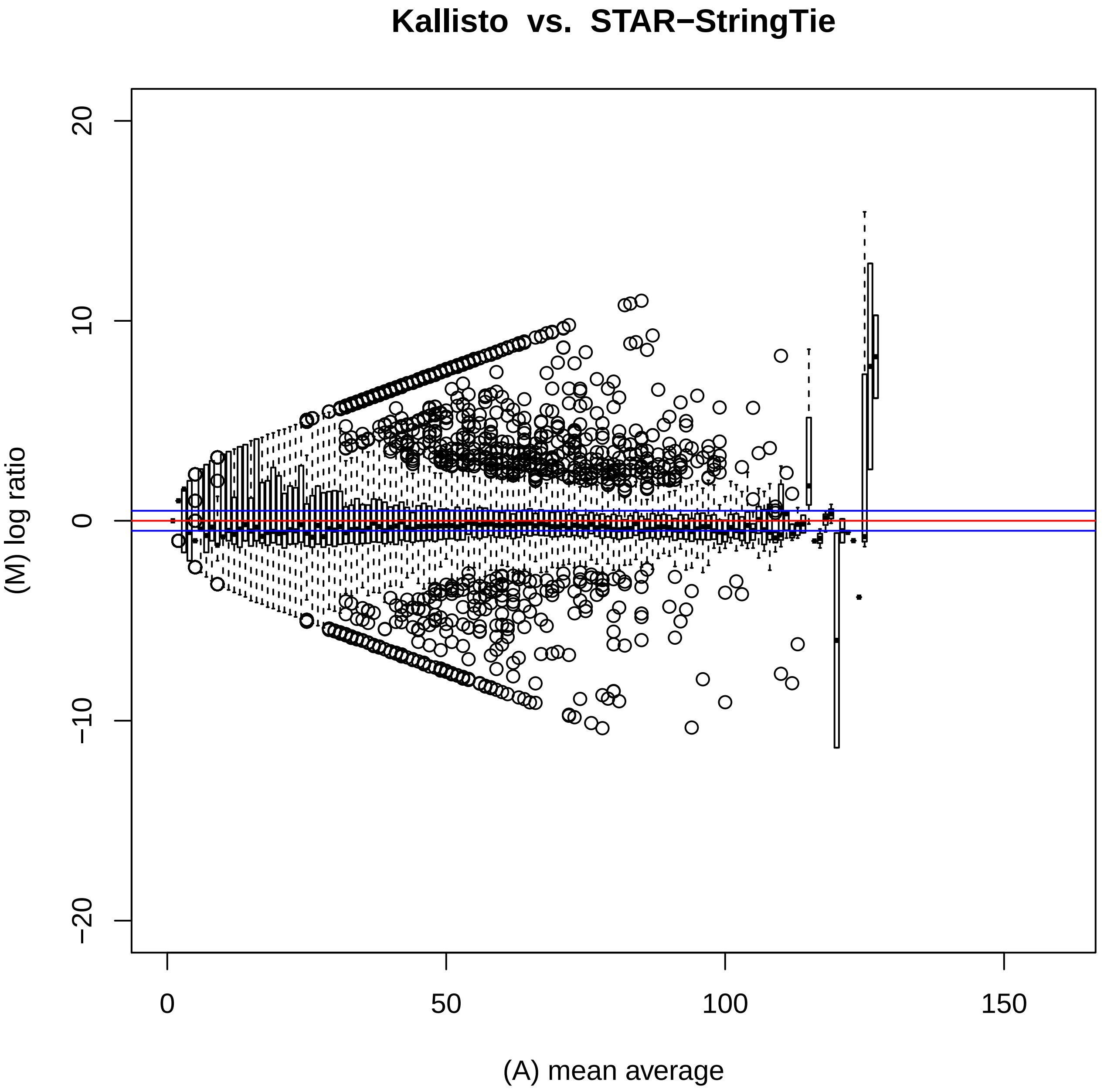
Kallisto vs. STAR−StringTie
(M) log ratio
(A) mean average
20
10
0
−10
−20
0
50
100
150

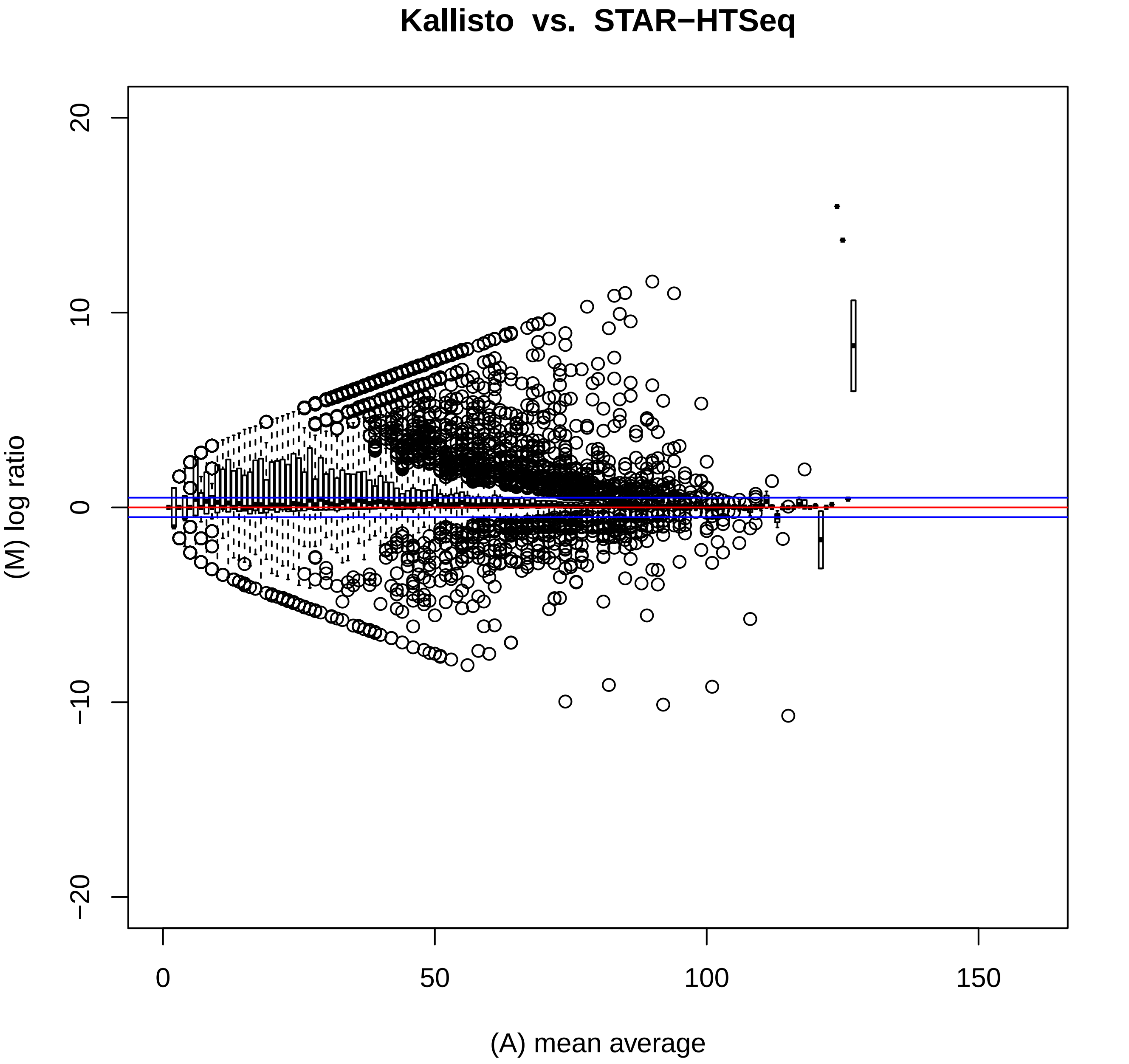

Kallisto vs. STAR−HTSeq
20
10
0
−10
−20
(M) log ratio
0
50
100
150
(A) mean average

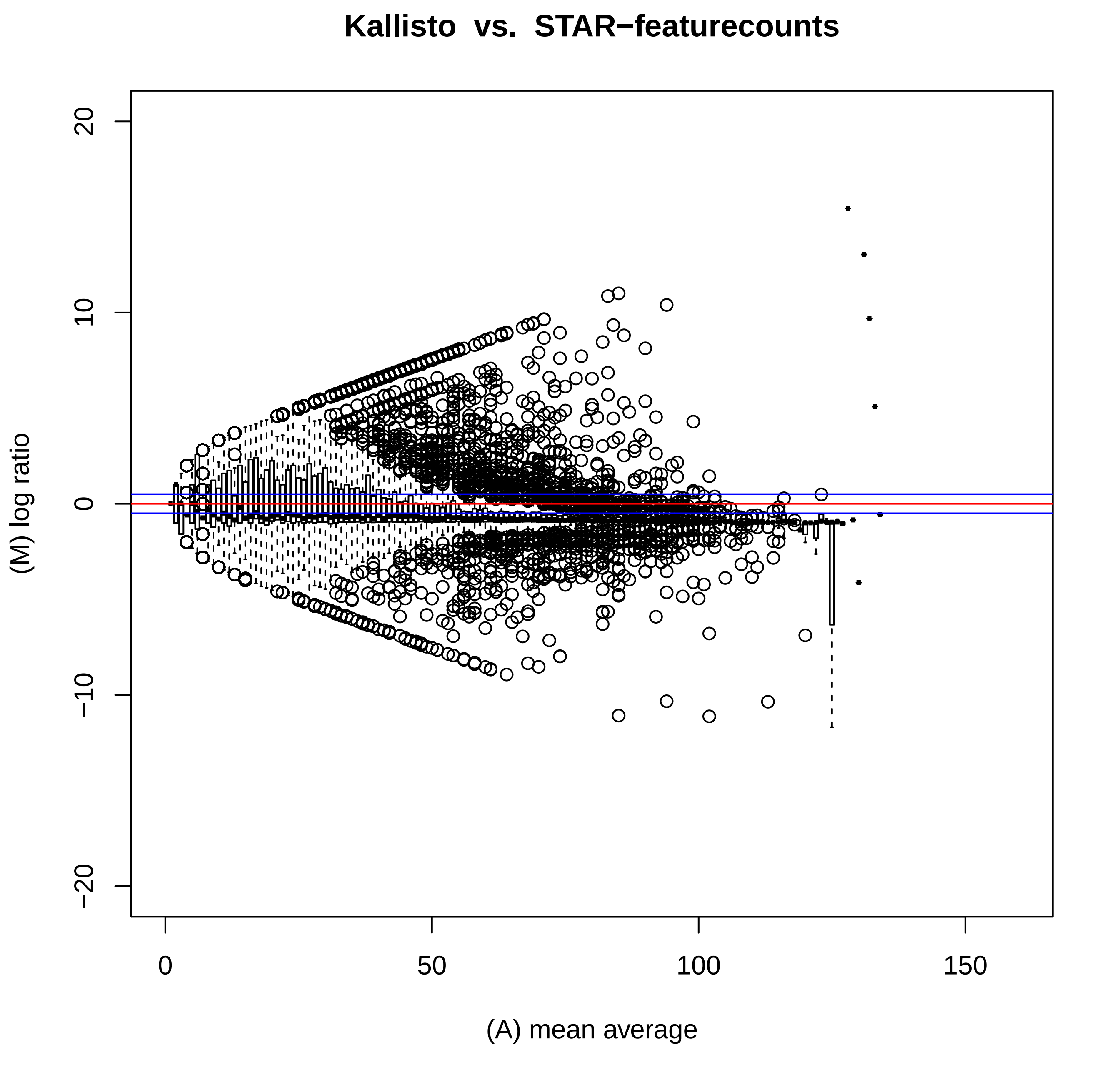

Kallisto vs. STAR−featurecounts
20
10
0
−10
−20
(M) log ratio
0
50
100
150
(A) mean average

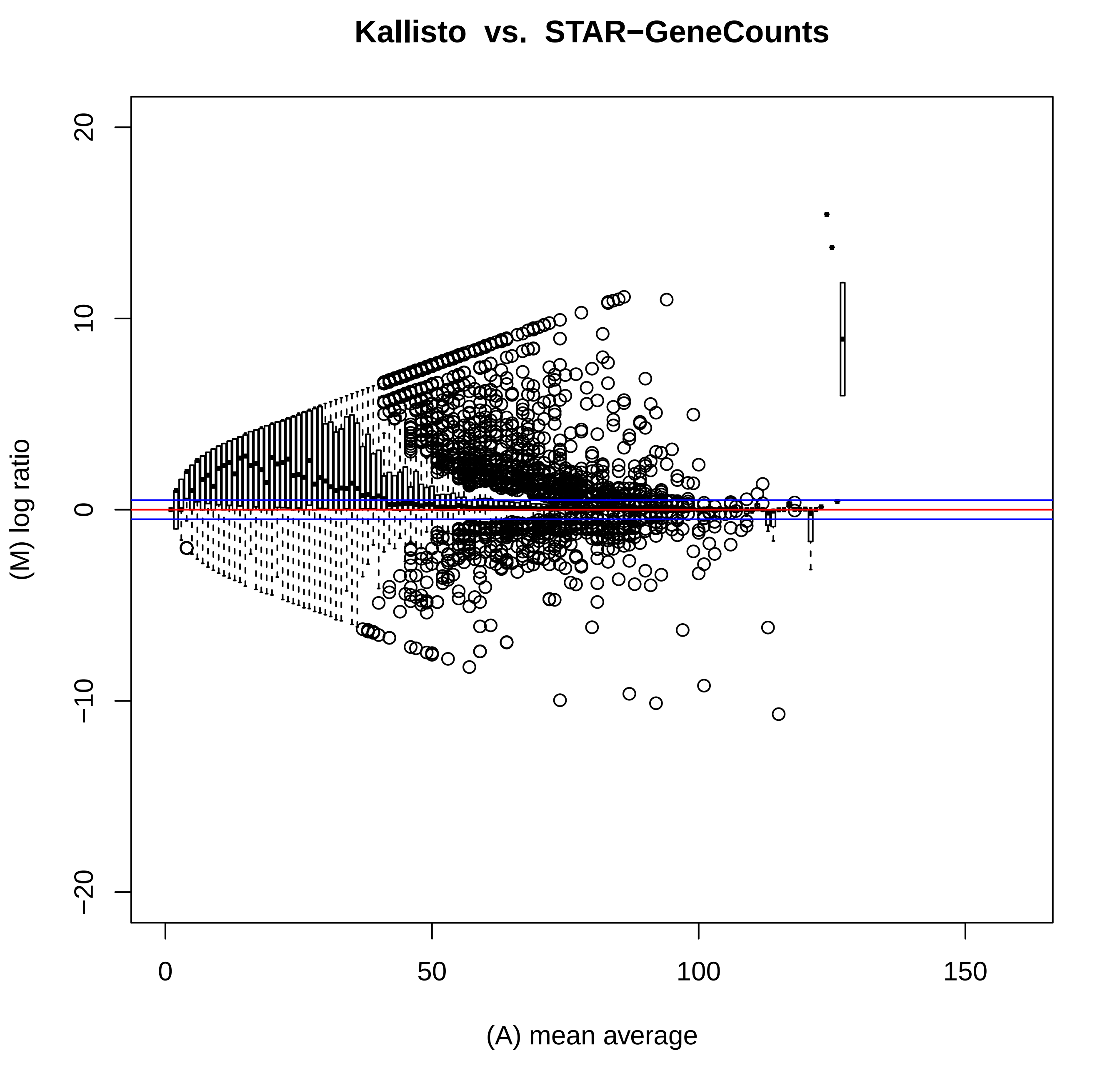
Kallisto vs. STAR−GeneCounts
(M) log ratio
(A) mean average
20
10
0
−10
−20
0
50
100
150

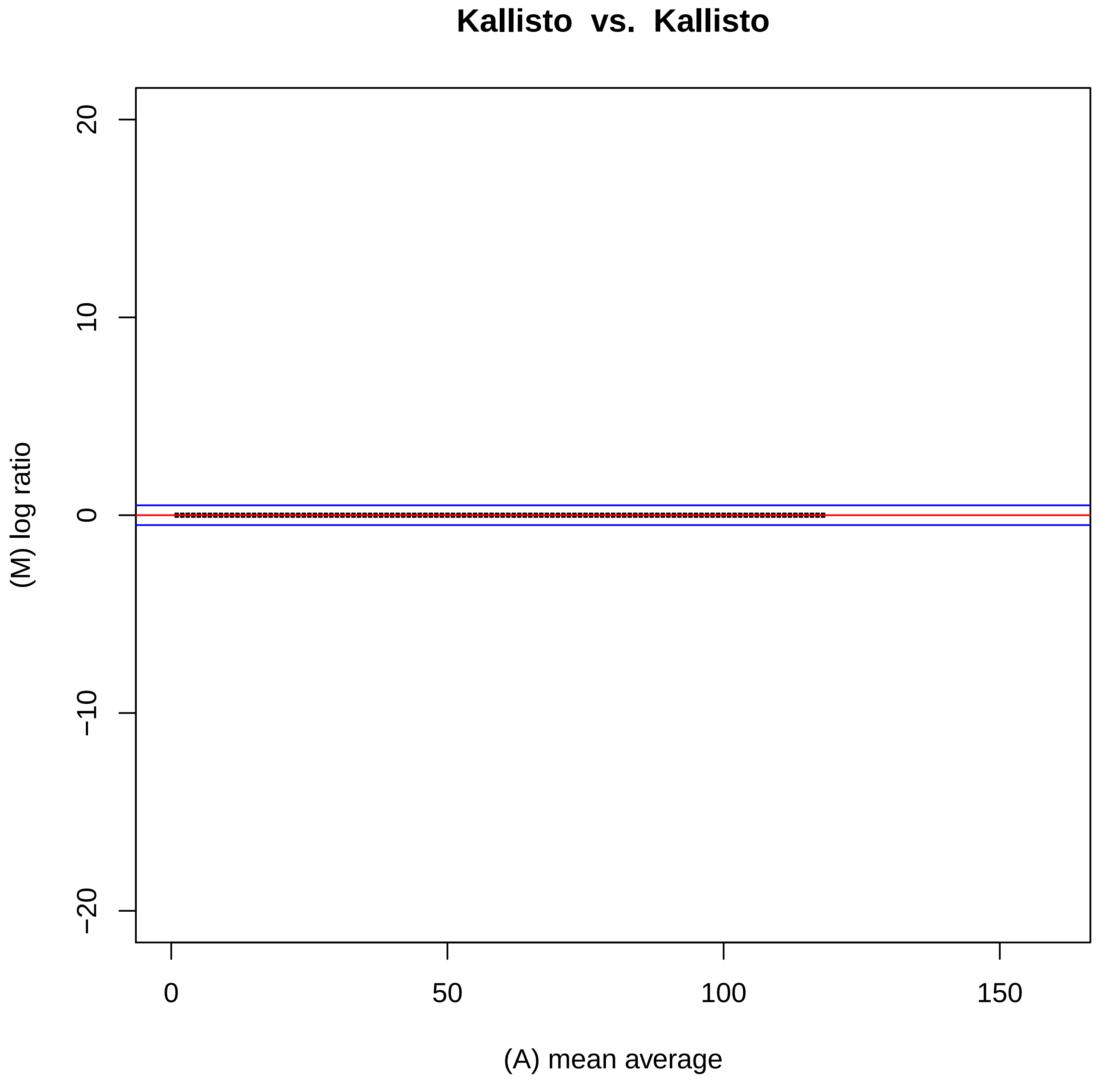
Kallisto vs. Kallisto
(M) log ratio
(A) mean average
20
10
0
−10
−20
0
50
100
150
