## Supplementary figures and images for "Effects of technical noise on bulk RNA-seq differential gene expression inference"

### S2 Data. Supplementary Fig. 13

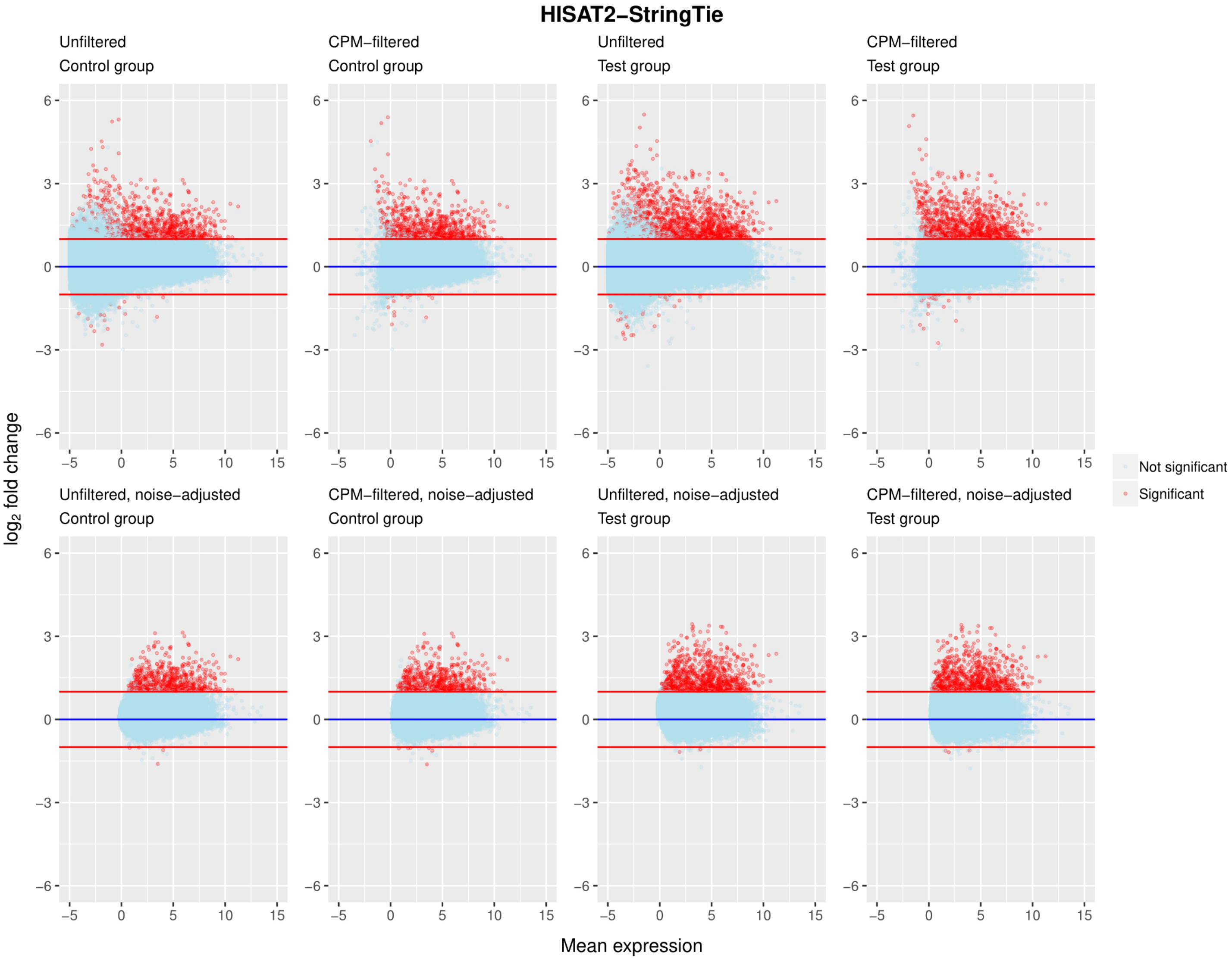

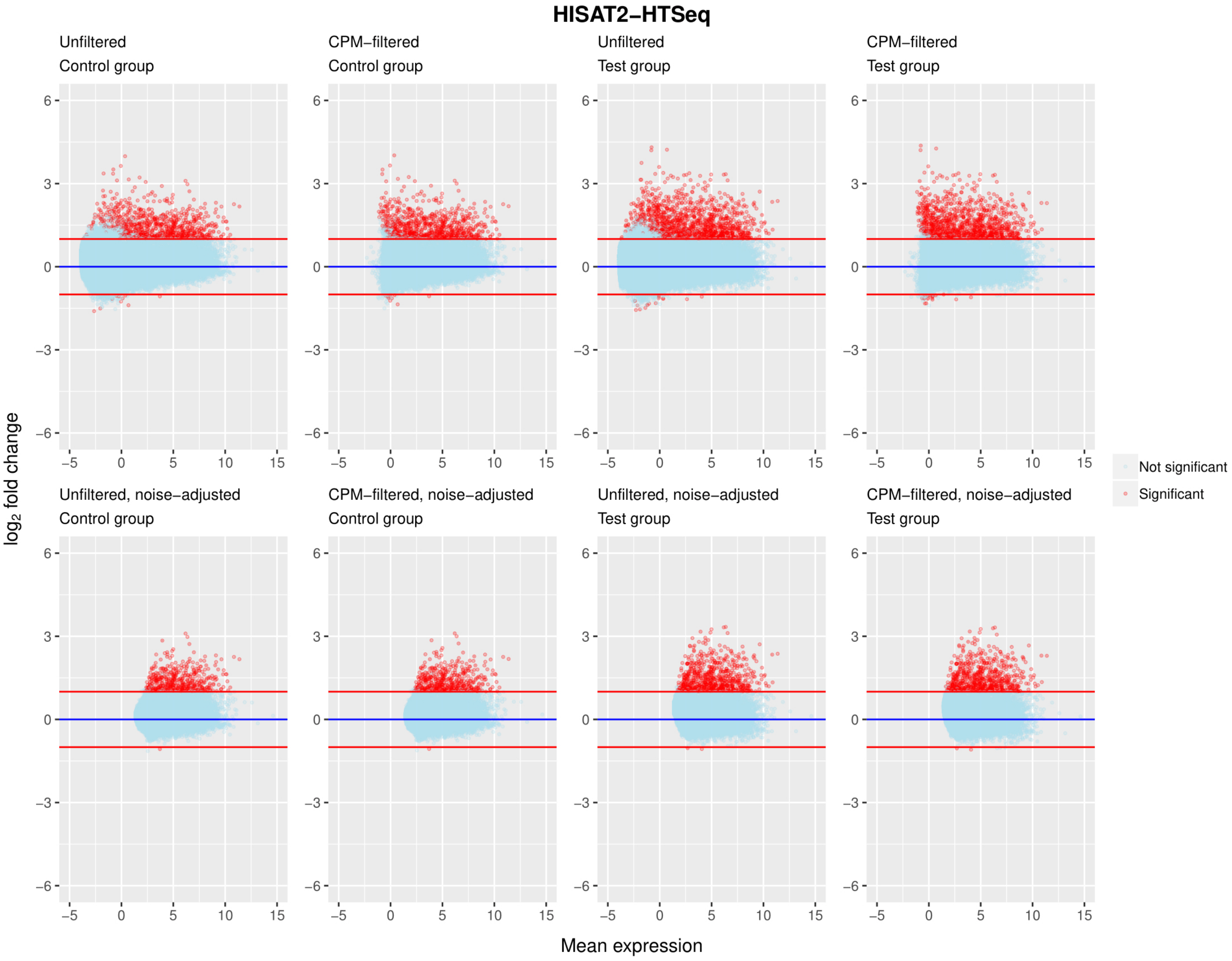

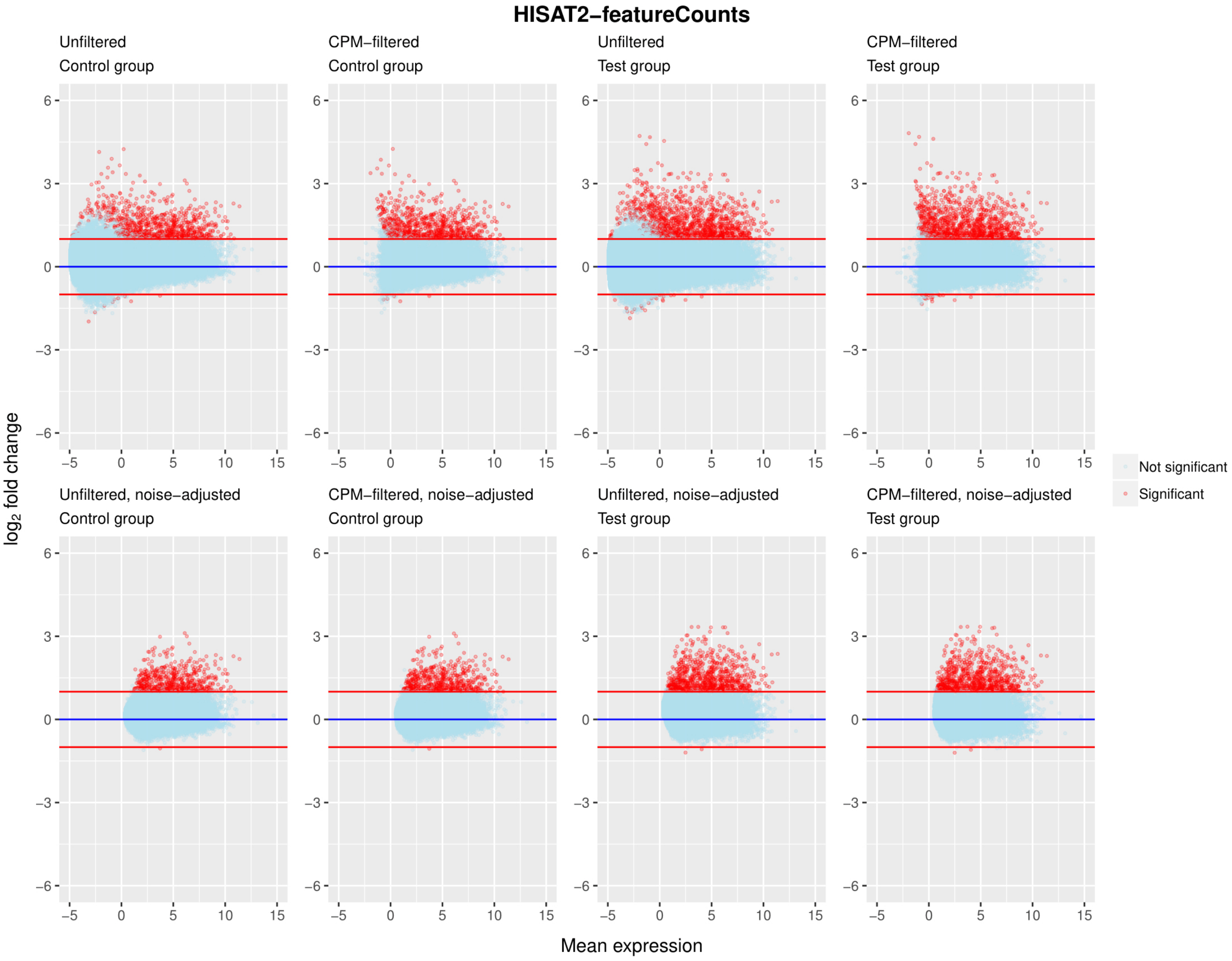

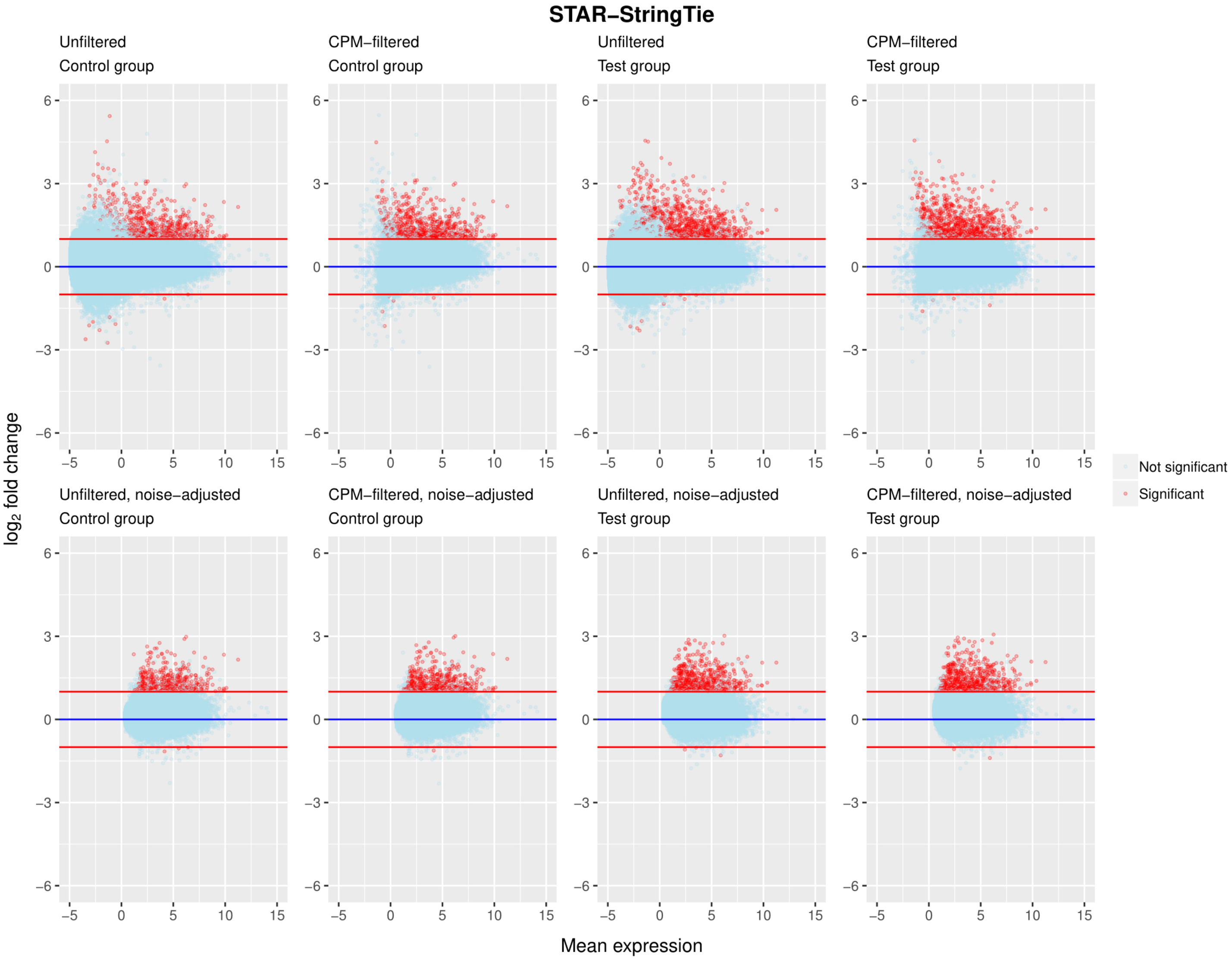

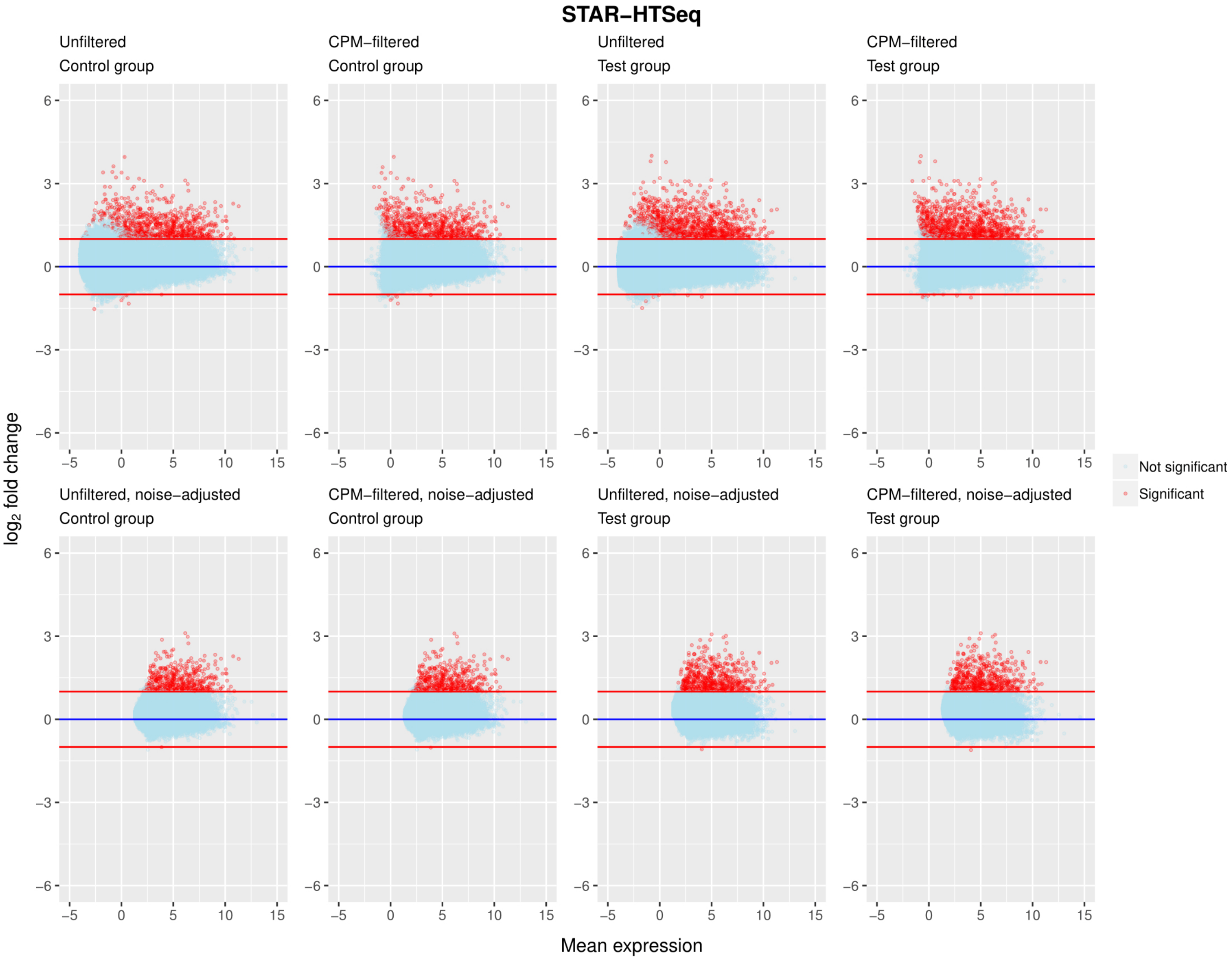

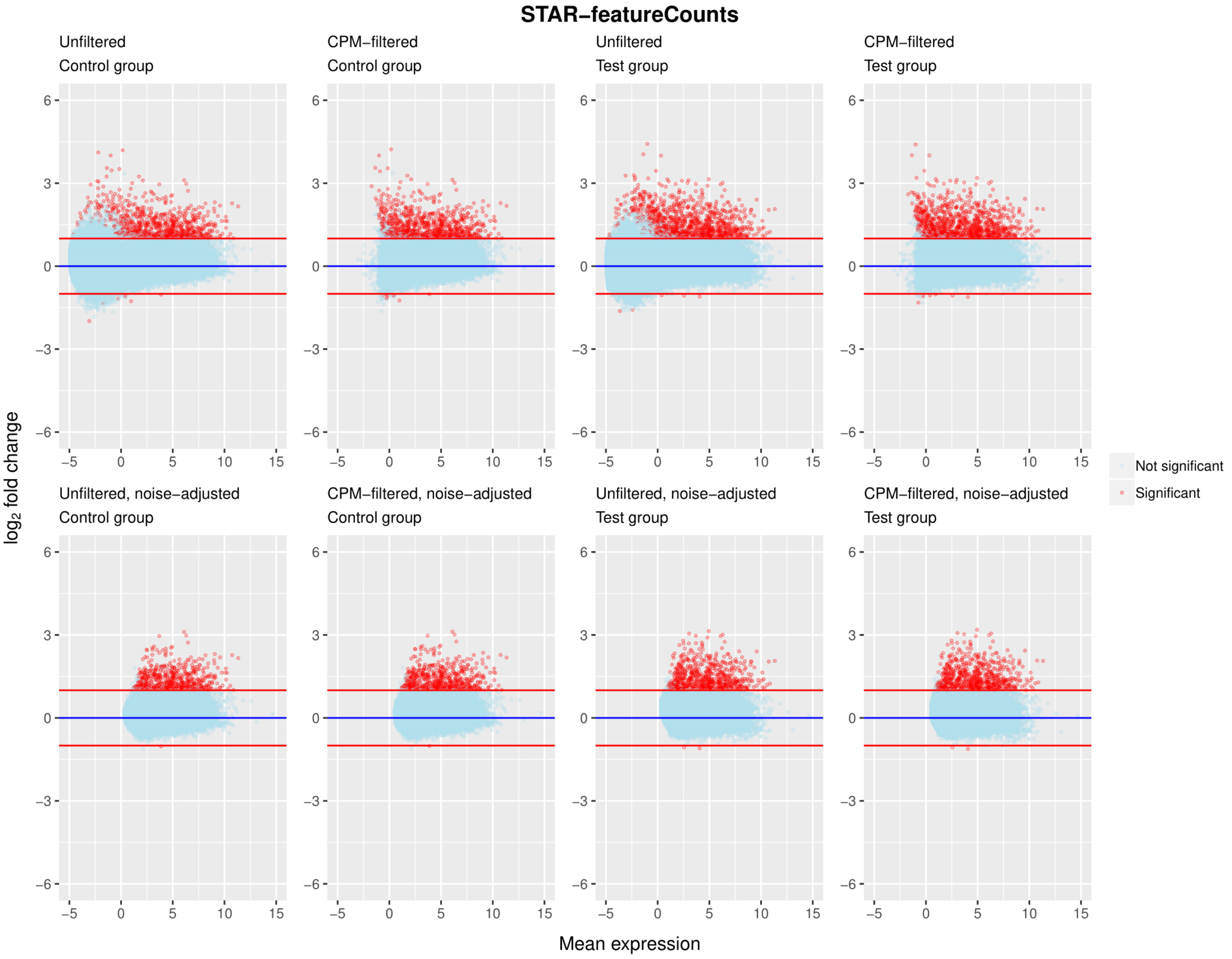

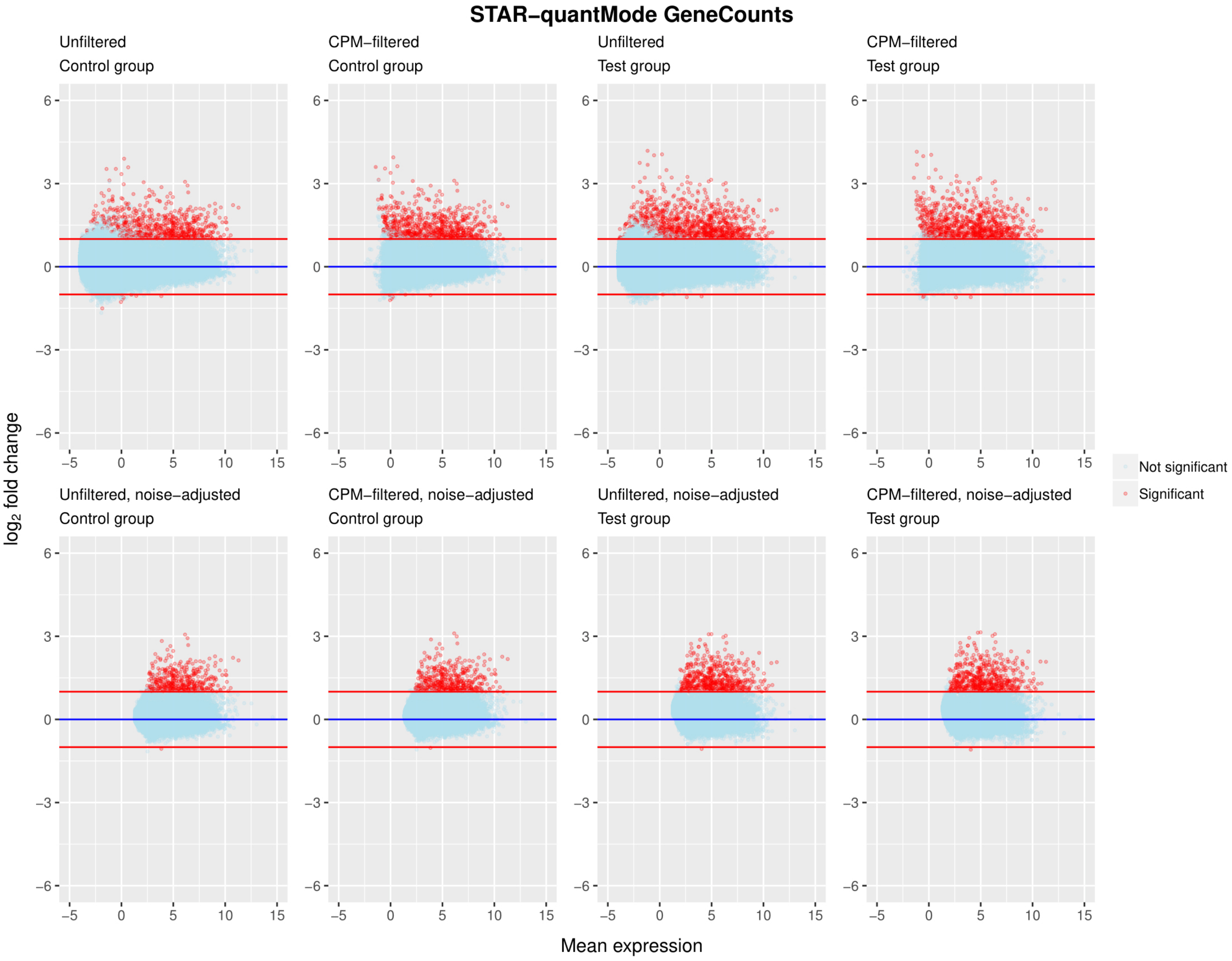

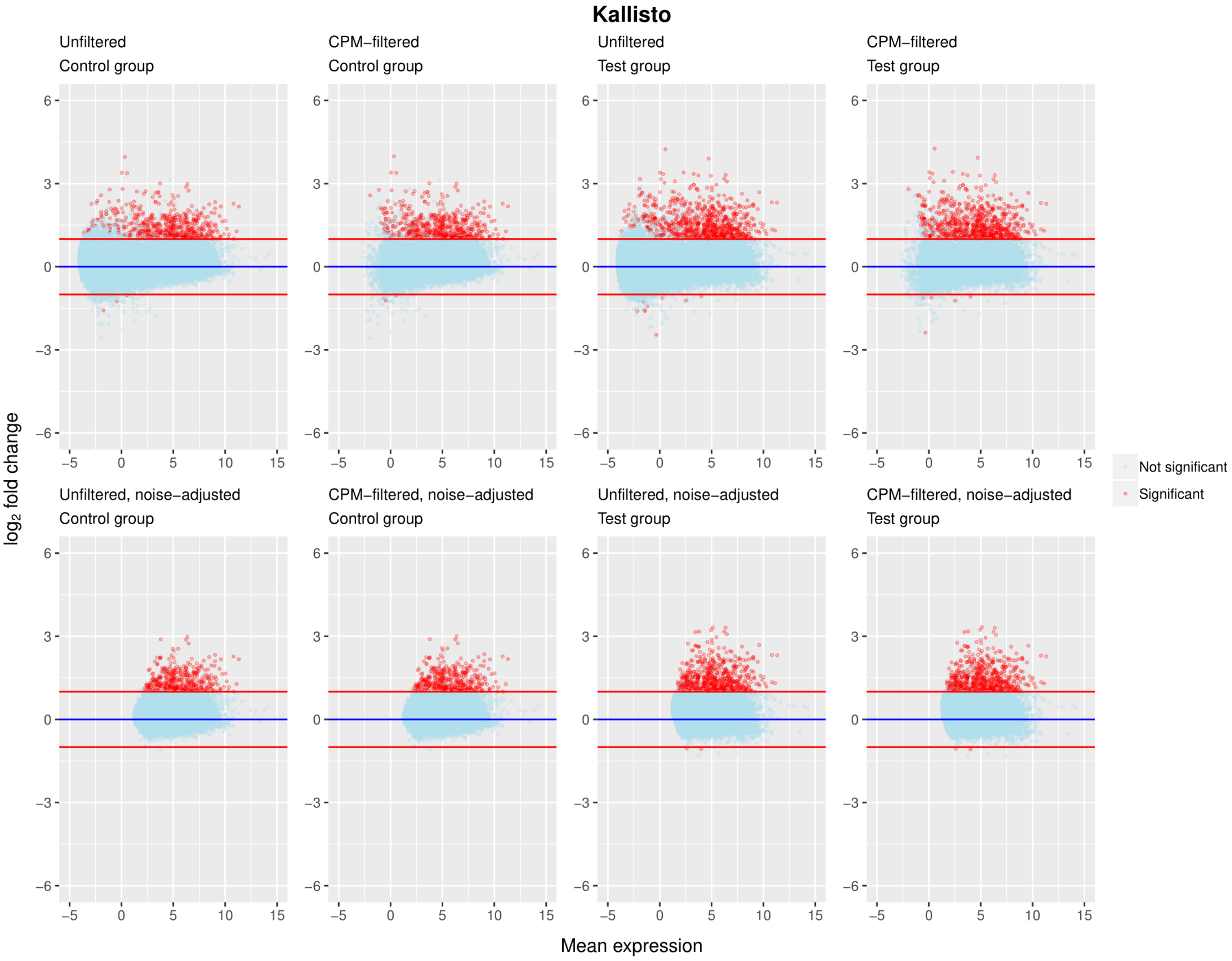
